# Programmable Recruitment of RNA-Binding Proteins Enables Small Molecule-Directed Destabilization of Nuclear Pre-mRNA

**DOI:** 10.64898/2026.08.25.746729

**Authors:** Xiaoxuan Su, Jielei Wang, Takahiro Ishii, Kisu Sung, Ryuichi Sekioka, Xueyi Yang, Patrick R. A. Zanon, Zhonglin Liu, Matthew D. Disney

## Abstract

Chemically induced proximity has not been systematically applied to control RNA fate. Here, a programmable platform was developed to identify RNA-binding proteins (RBPs) that can be recruited by small molecules to destabilize RNA. Using *microtubule-associated protein Tau* (*MAPT*) pre-mRNA as a model target, a heterobifunctional molecule was designed to bind both a ligandable structure in *MAPT* pre-mRNA and FKBP12^F36V^-tagged RBPs. Screening of a library of tagged RBPs identified several proteins that reduced *MAPT* RNA levels, including zinc finger protein 36 (ZFP36) and nanos C2HC-type zinc finger 3 (NANOS3). The approach was then extended from engineered proteins to an endogenous RBP. Using small molecule ligandability maps, a cysteine-reactive ligand for ZFP36 was identified. When this ligand was linked to the *MAPT*-binding small molecule, endogenous ZFP36 was recruited to *MAPT* mRNA, reducing its abundance in cells. Genetic and chemical controls demonstrated that activity was dependent on both RNA binding and ZFP36 recruitment, supporting an induced-proximity mechanism. These studies establish a general strategy for identifying new recruitable RBP effectors and should advance ribonuclease-targeting chimera (RiboTAC) technology by expanding the repertoire of effector proteins that can be harnessed for RNA degradation. More broadly, new effectors can be discovered through model reporter-based screens and translated to endogenous systems by mining known protein binders and ligandability maps, providing a systematic path to develop small molecules that control RNA stability, including RNAs targeted through structured regions of nuclear pre-mRNAs.

**TOC GRAPHIC:** 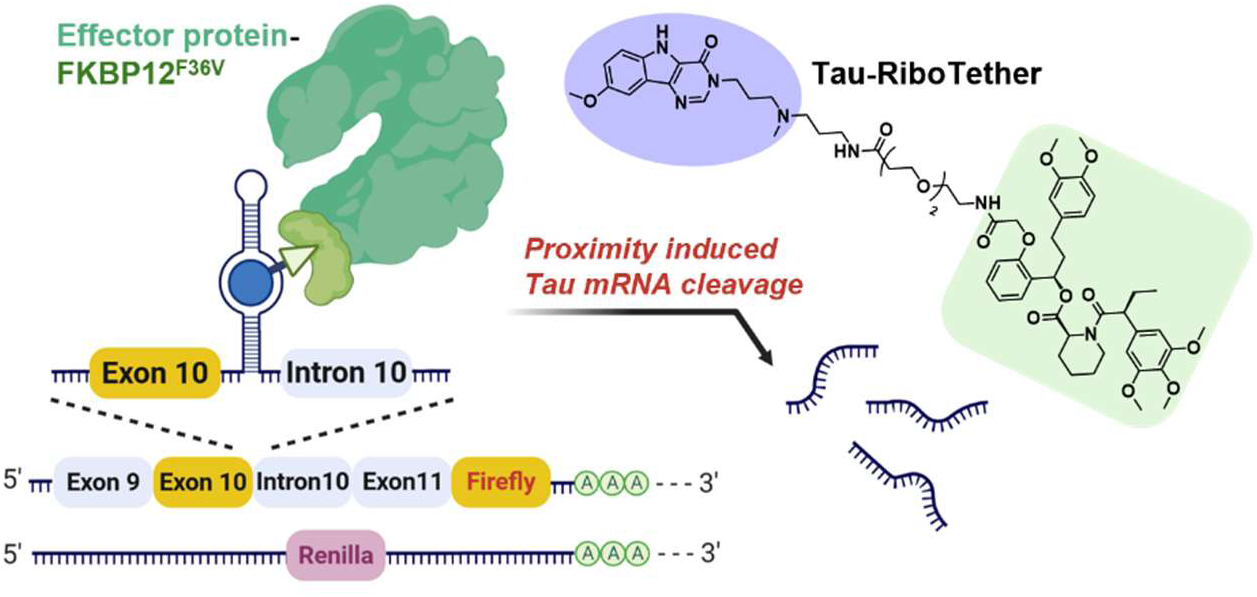

## INTRODUCTION

RNA plays diverse roles in biology, including encoding proteins, catalyzing chemical reactions^1, 2^, and regulating gene expression^3, 4^. Molecular entities that modulate RNA function or stability can provide tools for studying RNA biology and potential medicines for RNA-mediated diseases^5^. Oligonucleotide-based modalities, particularly antisense oligonucleotides (ASOs), recognize RNA through sequence complementarity^6^, which inhibits protein translation by occupying a target RNA^7^ or inducing its degradation by recruiting RNase H to cleave the RNA strand within an RNA-ASO heteroduplex in a catalytic and substoichiometric manner^8^.

Small molecules can also modulate RNA expression by binding folded, functional RNA structures^9, 10^ or by inducing proximity between effector proteins and target RNAs^10, 11^. One class of enzyme-recruiting heterobifunctional compounds, dubbed ribonuclease-targeting chimeras (RiboTACs), links an RNA-binding moiety to a ligand that engages RNase L and directs it to cleave a selected transcript^12^. Recruitment of RNase L has expanded the landscape of druggable RNAs by converting otherwise inactive RNA binders into active degraders in models of cancer^13^ and genetic disease^14^. Moreover, microtubule-associated protein 1 light chain 3 beta (LC3B; MAP1LC3B) was recently identified as a recruitable effector protein that triggers targeted RNA decay^15, 16^, inspired by targeted protein lysosomal degradation technologies^17^ such as autophagosome-tethering compounds (ATTECs)^18, 19^. However, the predominantly cytoplasmic localization of RNase L can limit its ability to act on nuclear RNAs, which are of considerable interest because many disease-driving transcripts are retained or processed in the nucleus^20, 21^. Nuclear RNA targets can also be depleted using direct cleavage approaches in which RNA-binding small molecules are appended to a bleomycin derivative that cleaves RNA oxidatively^22^. An enzymatic recruitment strategy capable of identifying ribonucleases or other RNA-destabilizing proteins compatible with nuclear RNA targets would further expand the scope of the RiboTAC approach.

Tethered-function assays have been developed to investigate the activities of RNA-binding proteins (RBPs) in living cells by artificially recruiting them to reporter RNAs through defined interactions^23^. For example, the well-characterized binding between bacteriophage MS2 stem-loops and the MS2 coat protein ^24^ has been widely used for this purpose^25, 26^. In a recent large-scale study^27^, 690 full-length RBP open reading frames were fused to a C-terminal MS2 coat protein domain and co-expressed with a luciferase reporter mRNA containing multimerized MS2 stem-loops in its 3′ untranslated region (3′ UTR)^24^. The resulting high-affinity MS2-MCP interaction recruits each RBP fusion to the reporter RNA, enabling functional screening based on changes in reporter output. Such approaches have provided important insights into RBP-mediated regulation of RNA stability and translation. However, few studies have demonstrated that RBPs identified through these systems can subsequently be harnessed by small molecules to modulate selected RNA targets.

Here, a model system was developed and implemented to chemically and systematically induce proximity between *microtubule-associated protein Tau* (*MAPT*; Tau) pre-mRNA and RBPs with RNA-stabilizing or RNA-destabilizing functions. The system uses a cell permeable heterobifunctional compound, dubbed **RiboTether**. In this study, the Tau-directed RiboTether (**Tau-RiboTether**) links a known Tau pre-mRNA-binding small molecule^28^ to **ortho-AP1867**, a ligand for the engineered FKBP12^F36V^ protein tag.^29^ A related protein-centric strategy, termed degradation tag (dTAG), uses heterobifunctional small molecules to recruit E3 ubiquitin ligases to FKBP12^F36V^-tagged protein targets, thereby inducing their degradation in cells^29^. In the approach described here, each candidate RBP was fused to FKBP12^F36V^, enabling its chemical recruitment to Tau mRNA upon treatment with **Tau-RiboTether**. Complementing existing genetic tethering strategies, this system provides a framework for determining whether individual RBPs can be chemically recruited to modulate RNA stability, identifying a broad range of effectors compatible with nuclear RNA targets, particularly those that promote RNA destabilization, and guiding the development of small molecules that recruit endogenous RBPs.

## RESULTS

### Design and construction of a programmable system for systematic recruitment of RBPs to nuclear RNAs

The chemical recruitment of specific RNA-binding proteins (RBPs) to nuclear RNA targets was conceptualized as requiring two components of molecular recognition: (i) programmable binding of an effector protein and (ii) robust engagement of the target RNA. An orthogonal chemical genetic recruitment system was adapted from the dTAG platform, in which E3 ligases such as cereblon (CRBN) have been recruited to substrate proteins fused to an engineered protein tag for targeted protein degradation^29–33^. The dTAG strategy is based on an engineered FK506-binding protein 12 (FKBP12) variant, FKBP12^F36V^, in which a high affinity binding cavity for the synthetic ligand **ortho-AP1867** is created by the F36V mutation^29^. FKBP12^F36V^ is selectively engaged by **ortho-AP1867**, while engagement of endogenous FKBP12 is minimized, thereby enabling orthogonal chemical control in cells^29^. Bivalent molecules recruiting FKBP12^F36V^-tagged protein kinase cyclin dependent kinase 9 (CDK9)^34^ or the fusion transcription factor EWSR1::FLI1^35^ to DNA sites bound by the transcriptional regulator BCL6 transcription repressor (BCL6) were also developed; these studies demonstrate proofs-of-concept to relocalize a transcriptional kinase or DNA-bound transcription factor to drive expression of pro-apoptotic genes in cancer cells. In the present study, fusion constructs between FKBP12^F36V^ and selected RBPs were generated (**Figure 1A**). The **ortho-AP1867** was then appended to a characterized RNA-binding small molecule and used to bridge the FKBP12^F36V^-tagged RBPs to the target RNA, thereby enabling proximity-induced changes in RNA abundance to be investigated.

**Figure 1.**
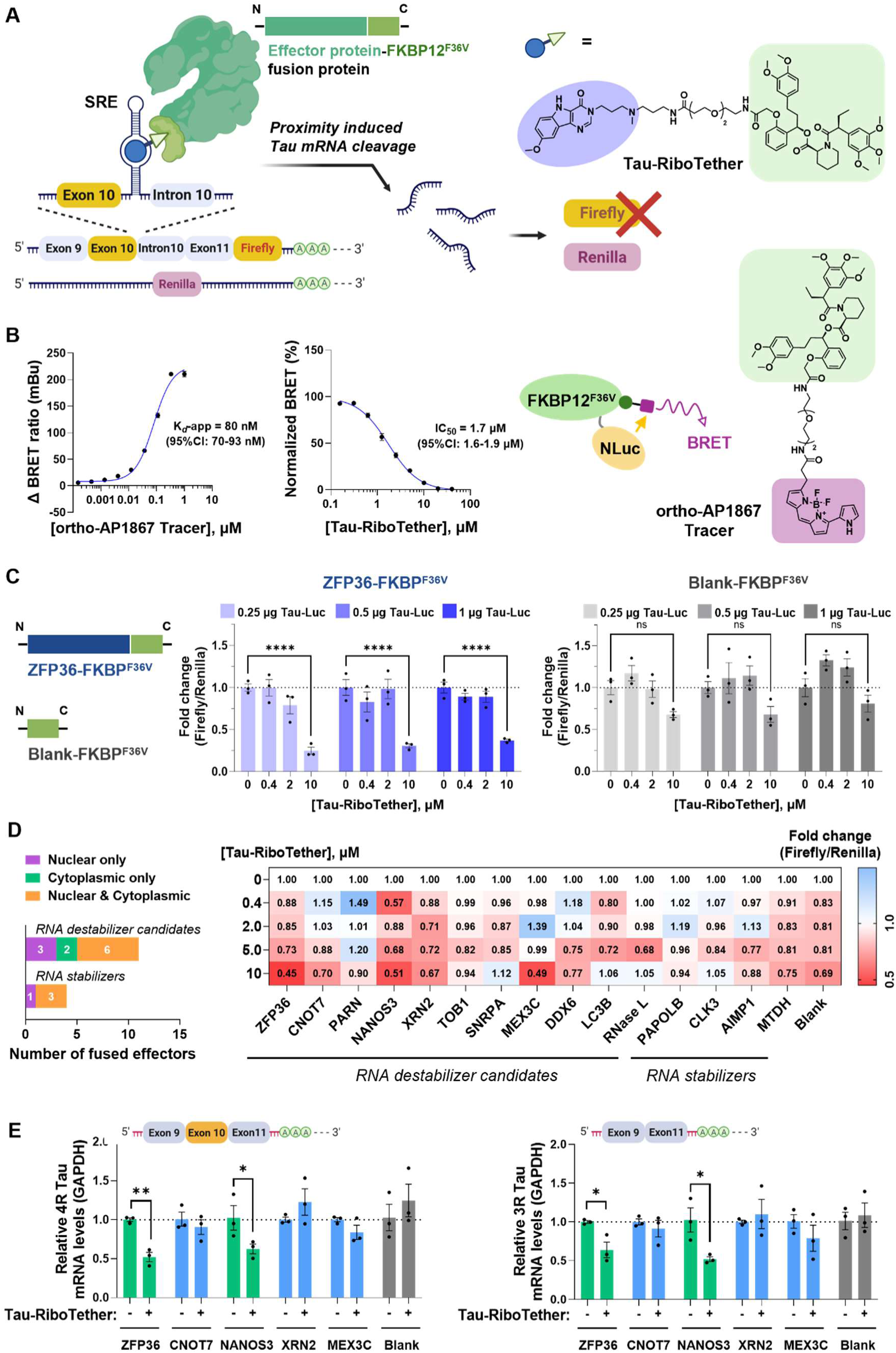
A chemically controlled FKBP12^F36V^-RBP tethering system identifies recruitable effectors that reduce *Tau-Luc* reporter transcript abundance. **(A)** Schematic of the chemically induced proximity system. Candidate RNA-binding proteins (RBPs) were expressed as C-terminal FKBP12^F36V^ fusion proteins. **Tau-RiboTether**, a heterobifunctional compound containing a *Tau* pre-mRNA-binding module and the FKBP12^F36V^ ligand **ortho-AP1867**, was designed to recruit each tagged effector to the splicing regulatory element at the exon 10-intron 10 junction of the *Tau-Luc* reporter transcript. Productive recruitment reduces *Tau-Luc* reporter transcript abundance and firefly luminescence without affecting the co-transfected Renilla luciferase control. **(B)** Cellular engagement of FKBP12^F36V^ by **Tau-RiboTether** was measured using NanoBRET. Binding of the **ortho-AP1867 tracer** to FKBP12^F36V^-NanoLuc produced an apparent affinity of 80 nM (95% CI, 70.0-92.6 nM; left). Tracer engagement was competitively displaced by **Tau-RiboTether** with an IC_50_ of 1.7 µM (95% CI, 1.6-1.9 µM; middle). The NanoBRET assay configuration and chemical structure of the **ortho-AP1867 tracer** are shown on right. **(C)** Benchmarking of the tethering system using ZFP36-FKBP12^F36V^ or Blank-FKBP12^F36V^. HeLa cells were transfected with 0.25, 0.5, or 1 µg of *Tau-Luc* plasmid, as well as Renilla plasmid and corresponding FKBP12^F36V^ plasmid, and treated with **Tau-RiboTether** at the indicated concentrations for 48 h. firefly luminescence was normalized to Renilla luminescence and expressed relative to the corresponding vehicle-treated condition (0.1% (v/v) DMSO). **(D)** Screening of 15 FKBP12^F36V^-tagged effectors, comprising 12 candidate RNA-destabilizing proteins and four RNA-stabilizing proteins. Candidate effectors were classified by reported subcellular localization (left). The heat map shows the mean firefly/Renilla fold change following treatment with **Tau-RiboTether** at the indicated concentrations for 48 h, normalized to the corresponding vehicle-treated condition (0.1% (v/v) DMSO; first row). **(E)** RT-qPCR analysis of the five RNA-destabilizing effectors prioritized from the luminescence screen. Relative levels of exon 10-containing *4R* and exon 10-lacking *3R Tau* reporter transcripts were measured in HeLa cells expressing the indicated FKBP12^F36V^ fusion protein following treatment with vehicle (0.1% (v/v) DMSO) or 10 µM of **Tau-RiboTether** for 48 h. Transcript levels were normalized to *GAPDH* and expressed relative to the corresponding vehicle-treated condition (0.1% (v/v) DMSO). Data are reported as the mean ± SEM from n = 3 biological replicates, with individual measurements shown. ns, not significant; *p < 0.0505, **p < 0.01, and ****p < 0.0001, as determined by two-way ANOVA with multiple-comparisons correction.

An FKBP12^F36V^-RBP library was constructed by selecting RBPs reported to be localized either predominantly within the nucleus or in both the nucleus and cytosol. Both RNA-destabilizing proteins (n = 17) and RNA-stabilizing proteins (n = 4) were selected (**Table S1**). Two additional RNA-destabilizing proteins with predominantly cytosolic localization, RNase L and LC3B, were included based on previous demonstrations of their chemical recruitment for targeted RNA degradation^10, 15, 16^. The mammalian expression vector pCMV6-FLAG was modified by HiFi assembly to incorporate the *FKBP12^F36V^* sequence, thereby generating a blank parent vector (*Blank-FKBP12^F36V^*). Each RBP coding sequence was subsequently inserted in-frame upstream of *FKBP12^F36V^* and separated from the tag by a short spacer, yielding C-terminally tagged fusion proteins. Robust expression of 15 of the 21 constructs was observed in HeLa cells following transient transfection, as confirmed by Western blotting (**Figure S2**).

To extend a chemically induced proximity approach to nuclear RNA targets, a luminescence-based reporter system was established for scalable detection of changes in RNA abundance induced by effector recruitment (**Figure 1A**). A previously characterized small molecule^28^ that binds an adenine bulge within the splicing regulatory element present at the exon 10-intron 10 junction of *MAPT* pre-mRNA^36, 37^ was used as the RNA-targeting module. The SRE controls the alternative splicing of exon 10, producing a transcript with three microtubule binding domains (exon exclusion; 3R isoform) or four domains (exon inclusion; 4R); that latter is aggregation prone and is associated with tauopathies^38, 39^. Upon small molecule binding, the SRE structure is stabilized, U1 small nuclear ribonucleoprotein (snRNP) binding is impaired, and a modest shift in exon 10 splicing is produced, thereby altering the *4R:3R* Tau isoform ratio^28^.

Because the targeted SRE is located at an exon-intron junction in *MAPT* pre-mRNA, it was selected as a model for evaluating effector activity against a nuclear RNA target. A *MAPT* mini-gene reporter in which exon 10 was fused in-frame to firefly luciferase was employed, allowing 4R Tau output to be monitored^40, 41^. We hypothesized that upon treatment with the bifunctional *Tau*-binder-FKBP12^F36V^-recruiter, dubbed **Tau-RiboTether**, proximity between the FKBP12^F36V^-tagged effector and the model *Tau* pre-mRNA could be induced. Thus, a reduction in firefly luciferase activity without a corresponding change in co-transfected Renilla luciferase could be used as an initial indicator of effector-dependent activity. Changes in the abundance of the *3R* and *4R Tau* reporter transcripts could then be subsequently quantified using established Reverse Transcription-quantitative Polymerase Chain Reaction (RT-qPCR) or droplet digital PCR (ddPCR) assays, allowing the effects of chemically recruited RBPs on the model nuclear RNA target to be evaluated directly.

### FKBP12^F36V^-RBP library screening identifies two effectors that destabilize Tau RNA

Cellular engagement of FKBP12^F36V^ by **ortho-AP1867** was validated using a NanoLuciferase bioluminescence resonance energy transfer (NanoBRET) assay. FKBP12^F36V^ was fused to the N-terminus of NanoLuciferase using the same linker design employed in construction of the FKBP12^F36V^-RBP library construction (**Figures 1B, left and S1A**). A marked increase in BRET signal was induced by the **ortho-AP1867 Tracer,** with an apparent binding affinity of 80 nM (95% CI: 70-93 nM, **Figure 1B, left**). This signal was competitively displaced dose dependently by **ortho-AP1867** (IC_50_ = 4.4 µM, 95% CI: 3.9-5.5 μM; **Figure S3A, right**), **Tau-RiboTether** (IC_50_ = 1.7 µM, 95% CI: 1.6-1.9 μM; **Figure 1B, right**), but could not be displaced by **Tau binder** treatment up to 40 μM (**Figure S3B, right**). Collectively, these results indicate specific engagement of FKBP12^F36V^ by the chimeric **Tau-RiboTether** compound.

To determine whether functional proximity could be induced by **Tau-RiboTether**, zinc finger protein 36 homolog (ZFP36) was initially evaluated as a benchmark RNA-destabilizing effector^27, 42^. AU-rich element (ARE)-mediated RNA decay is promoted by ZFP36 through binding to regulatory elements, after which pre-mRNA polyadenylation can be inhibited in the nucleus or deadenylase and decapping complexes can be recruited in the cytoplasm^43, 44^. ZFP36 binding to pre-mRNAs in the nucleus has also been reported, with downstream cytoplasmic decay being licensed after export of ZFP36-containing mRNA-processing ribonucleoprotein complexes^45^. Potential ZFP36 binding sites near the *Tau* SRE were also identified using RBPmap^46^ (**Figure S4**), which maps experimentally defined RBP-binding motifs while accounting for the surrounding sequence context.

HeLa cells were co-transfected with plasmids encoding ZFP36-FKBP12^F36V^, the *MAPT* mini-gene reporter (*Tau-Luc*), and the Renilla luciferase internal control. The *MAPT* mini-gene contains a disease-relevant disinhibition-dementia-parkinsonism-amyotrophy complex (DDPAC) mutation in the SRE^40^, which shifts splicing toward the 4R isoform. Following transient transfection (0.25-1 µg of *Tau-Luc* plasmid), a dose dependent decrease in Tau-Luc firefly luciferase activity was observed after treatment with up to 10 µM of **Tau-RiboTether** (**Figure S5A**). No reduction in Renilla luminescence (**Figure S5B**) or detectable cytotoxicity (**Figure S5C**) was observed in the concentration range used. Consequently, a marked reduction in the normalized firefly/Renilla ratio was observed (**Figure 1C, left**).

To further validate this result, activity was measured across different Tau-Luc expression levels by altering the amount of plasmid transfected. At 10 µM of **Tau-RiboTether**, relative Tau-Luc firefly luminescence (as compared to vehicle-treated cells; not normalized to Renilla luciferase) was reduced to 26 ± 5% following transfection with 0.25 µg of *Tau-Luc* plasmid (p < 0.0001), 36 ± 4% following transfection with 0.5 µg of plasmid (p < 0.0001), and 39 ± 1% following transfection with 1 µg (p < 0.0001) (**Figure S5A**). Following normalization to Renilla luminescence, the firefly:Renilla ratio was relatively unaffected, with reductions to 25 ± 4% (p < 0.0001), 31 ± 2% (p < 0.0001), and 37 ± 1% (p < 0.0001), respectively (**Figure 1C, left**), supporting the robustness of the reporter system.

In cells expressing Blank-FKBP12^F36V^, no significant reduction in firefly luciferase activity was observed after treatment with up to 10 µM **Tau-RiboTether** (**Figure S5D**). A small increase in Renilla signal was observed at 10 µM of **Tau-RiboTether**, reaching 123 ± 2% with 0.25 µg *Tau-Luc* (p < 0.0001), 116 ± 1% with 0.5 µg *Tau-Luc* (p < 0.001), and 19 ± 4% with 1 µg of *Tau-Luc* (p < 0.0001) (**Figure S5E**). Nevertheless, no meaningful change in the normalized firefly ratio was produced (**Figure 1C, right**). These results supported an effector-dependent mechanism for suppression of the Tau reporter.

Screening of the FKBP12^F36V^-RBP library was performed using a fixed input of 1 µg of *Tau-Luc* plasmid, which provided the optimal signal-to-noise (S/N) ratio (**Figure S6A-B**). Following treatment with **Tau-RiboTether** for 48 h, dose dependent reduction in normalized firefly luminescence was observed for five RNA-destabilizing RBP fusions: ZFP36-, CCR4-NOT transcription complex subunit 7 (CNOT7), nanos C2HC-type zinc finger 3 (NANOS3), 5’-3’ exoribonuclease 2 (XRN2), and Mex-3 RNA binding family Member C (MEX3C) (**Figure 1D, right; Figure S7**). In contrast, no significant activity was observed for Blank-FKBP12^F36V^ or the four RNA-stabilizing RBP fusions, poly(A) polymerase beta (PAPOLB), CDC like kinase 3 (CLK3), aminoacyl tRNA synthetase complex interacting multifunctional protein 1 (AIMP1), and metadherin (MTDH) (**Figure 1D, right**). The strongest effects were associated with ZFP36-FKBP12^F36V^ and NANOS3-FKBP12^F36V^, for which normalized luminescence values of 45 ± 6% and 51 ± 4%, respectively, were observed following treatment with 10 µM of **Tau-RiboTether** (p < 0.01). No significant effects were induced by the predominantly cytoplasmic RBPs LC3B and RNase L^47^ (**Figure 1D, right**), consistent with an important role for effector localization in the modulation of nuclear RNA targets.

The five prioritized RNA-destabilizing RBP fusions were subsequently evaluated by RT-qPCR in the same reporter system. Substantial reductions in *Tau* reporter RNA abundance were observed only upon treatment of ZFP36-FKBP12^F36V^- or NANOS3-FKBP12^F36V^-expressing cells with **Tau-RiboTether** (10 µM; 48 h) (**Figure 1E**). Both the exon 10-containing *4R* and exon 10-lacking *3R Tau* reporter transcripts were reduced. In cells expressing ZFP36-FKBP12^F36V^, relative *4R* and *3R* transcript levels were reduced to 48 ± 6% (p < 0.01) and 36 ± 10% (p < 0.05), respectively, as compared to vehicle-treated cells (**Figure 1E**). In cells expressing NANOS3-FKBP12^F36V^, relative *4R* and *3R* transcript levels were reduced to 38 ± 6% (p < 0.05) and 48 ± 3% (p <0.05), respectively (**Figure 1E**). Across all six transfection conditions, changes in the raw C_t_ values were consistent with the normalized changes in Tau RNA abundance (**Figure S8A-B**). In the corresponding no-reverse-transcription controls, C_t_ values were increased by more than 7.5 cycles relative to samples subjected to reverse transcription (**Figure S8C**), indicating efficient removal of *Tau-Luc* plasmid DNA during RNA preparation and minimal interference with the RT-qPCR measurements. No changes in *4R* or *3R* Tau reporter RNA abundance were observed following expression of Blank-FKBP12^F36V^ and treatment with 10 µM of **Tau-RiboTether** for 48 h (**Figure 1E**), consistent with the effector-dependent activity observed in the initial Tau-Luc screen (**Figure 1D, right**). The other three effectors, CNOT7-, XRN2-, and MEX3C-FKBP12^F36V^, reduced firefly luciferase activity but did not reduce *Tau* mRNA levels, which might be attributed to steric inhibition of translation rather than destabilization/cleavage of the transcript. Collectively, ZFP36 and NANOS3 were identified as recruitable effectors whose chemically induced proximity to *Tau* pre-mRNA was associated with reduced reporter output and decreased abundance of both *3R* and *4R* Tau reporter transcripts.

### Reduction of *Tau-Luc* reporter RNA is dependent on induced proximity and effector recruitment

The requirement for induced proximity was assessed using genetic and chemical controls designed to disrupt either RNA engagement or effector recruitment. Dose dependent reductions in *3R* and *4R Tau* reporter transcript abundance were observed following **Tau-RiboTether** treatment in cells expressing ZFP36-FKBP12^F36V^ or NANOS3-FKBP12^F36V^ (**Figures 2A and S9A-B**). In contrast, no significant reductions were observed in cells expressing Blank-FKBP12^F36V^ in cells transfected with a low (0.25 μg; **Figure S9A-B**) or high (1 μg; **Figure 2A**) amount of *Tau-Luc* plasmid. The *3R* and *4R* Tau reporter transcript copy numbers were further confirmed by quantification using ddPCR (**Figure S9C-E**), supporting an effector-dependent decrease in RNA abundance.

**Figure 2.**
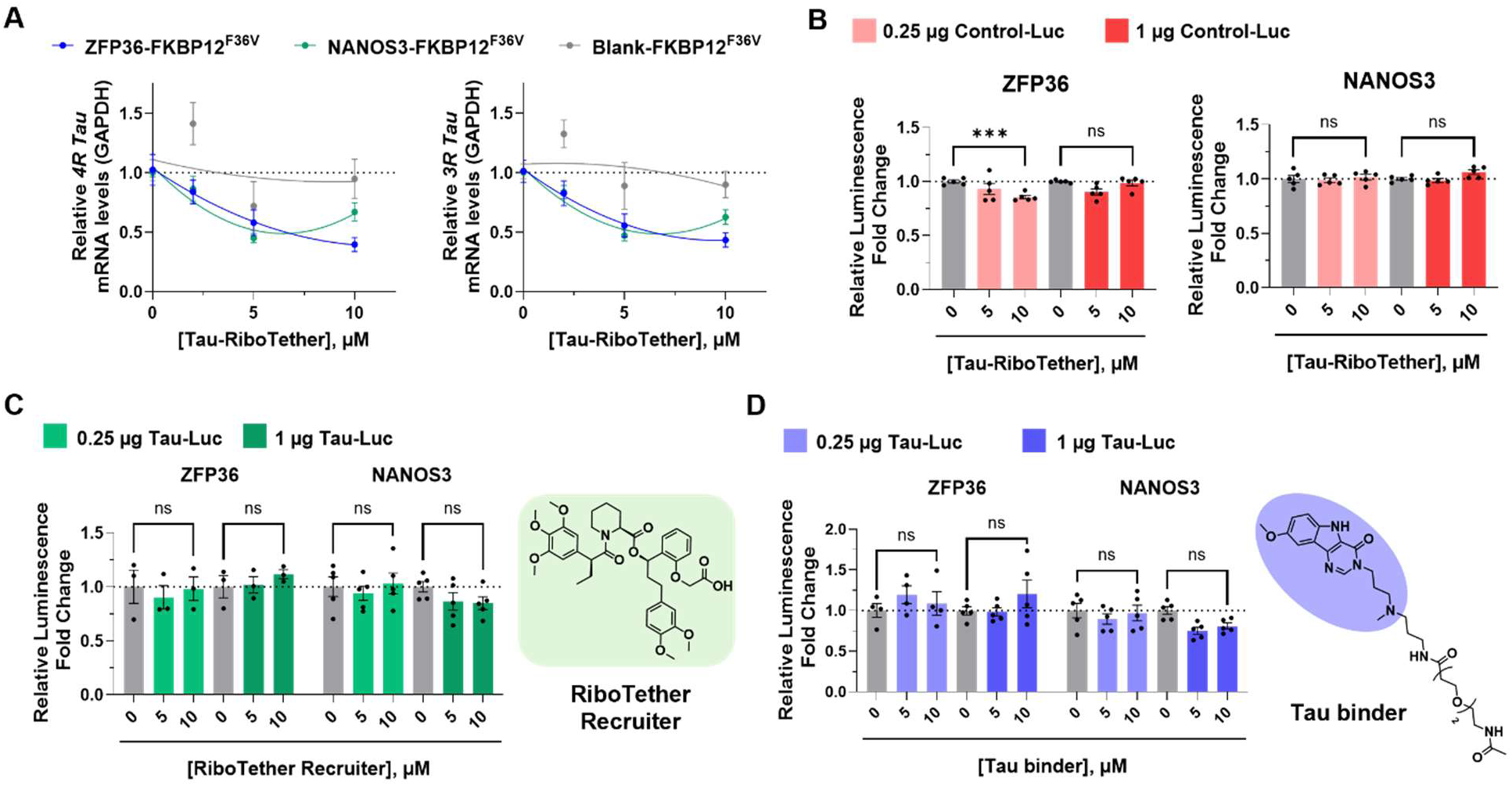
Reduction of *Tau-Luc* transcript abundance requires recognition of the *Tau* SRE and recruitment of an RNA-destabilizing effector. **(A)** Dose dependent effects of **Tau-RiboTether** on *4R* and *3R Tau-Luc* transcript abundance. HeLa cells were co-transfected with 1 μg of *Tau-Luc* plasmid and a plasmid encoding ZFP36-FKBP12^F36V^, NANOS3-FKBP12^F36V^, or Blank-FKBP12^F36V^ followed by treatment with **Tau-RiboTether** at the indicated concentrations for 48 h. Transcript levels were measured by RT-qPCR, normalized to *GAPDH*, and expressed relative to the corresponding vehicle-treated condition (0.1% (v/v) DMSO). Curves are included as visual guides; no potency values were derived from these fits. **(B)** Dependence of **Tau-RiboTether** activity on *Tau* sequence. A control firefly luciferase reporter plasmid lacking the *Tau* sequence (0.25 or 1 μg) was transfected into HeLa cells expressing ZFP36-FKBP12^F36V^ or NANOS3-FKBP12^F36V^. Relative luminescence was measured after treatment with **Tau-RiboTether** at the indicated concentrations for 48 h and expressed relative to the corresponding vehicle-treated condition (0.1% (v/v) DMSO). **(C)** Effects of the monofunctional **RiboTether Recruiter (ortho-AP1867)**, which contains the FKBP12^F36V^-binding module but lacks the *Tau* RNA-binding module. HeLa cells co-expressing Tau-Luc and either ZFP36-FKBP12^F36V^ or NANOS3-FKBP12^F36V^ were treated with the **RiboTether Recruiter** at the indicated concentrations for 48 h. **(D)** Effects of the monofunctional **Tau binder**, which contains the *Tau* RNA-binding module but lacks the FKBP12^F36V^-recruiting module. HeLa cells co-expressing Tau-Luc and either ZFP36-FKBP12^F36V^ or NANOS3-FKBP12^F36V^ were treated with the **Tau binder** at the indicated concentrations for 48 h. For **(B-D)**, data are reported as the mean ± SEM from n = 5 biological replicates, with individual measurements shown. ns, not significant; **p < 0.001, as determined by two-way ANOVA with multiple-comparisons correction.

When the *Tau* mini-gene was replaced with a control firefly luciferase reporter lacking the *Tau* sequence, the activity of **Tau-RiboTether** was largely abolished at concentrations up to 10 μM. Minimal (<15%) or no reduction in normalized luminescence was observed following co-transfection with ZFP36-FKBP12^F36V^ or NANOS3-FKBP12^F36V^ (**Figure 2B**). Similar results were obtained when either 0.25 μg or 1 μg of the control firefly luciferase plasmid was used (**Figure 2B**), indicating that the reduction in reporter signal was dependent on the presence of the ligandable *Tau* sequence.

The requirement for recognition of the *Tau* SRE was further evaluated using a mutant *Tau-I17T-Luc* reporter. In this construct, a uridine is placed opposite the bulged adenine, thereby eliminating the A-bulge required for recognition by the *Tau* RNA-binding small molecule and hence **Tau-RiboTether**^28^. Following transfection of either 0.25 μg or 1 μg of *Tau-I17T-Luc* and treatment with 10 μM of **Tau-RiboTether**, the response was substantially attenuated in cells expressing ZFP36-FKBP12^F36^V (**Figure S10A**) and abolished in cells expressing NANOS3-FKBP12^F36V^ (**Figure S10B**). These results were consistent with a requirement for recognition of the *Tau* SRE to produce efficient transcript reduction.

The requirement for both components of **Tau-RiboTether** was assessed using two monofunctional control compounds: **Tau binder**, which contains only the RNA-binding module and linker, and **RiboTether Recruiter** (**ortho-AP1867**), which contains only the FKBP12^F36V^-recruiting module. In cells expressing ZFP36-FKBP12^F36V^, no significant changes in Tau-Luc luminescence were observed following treatment (48 h) with either control compound at concentrations up to 10 μM, regardless of whether 0.25 μg or 1 μg of *Tau-Luc* was used (**Figure 2C-D**). Similarly, no significant reduction in luminescence was observed in cells expressing NANOS3-FKBP12^F36V^ and transfected with 0.25 μg of *Tau-Luc* mini-gene. When 1 μg of *Tau-Luc* plasmid was transfected, only modest and statistically insignificant reductions of <20% were observed with either control compound (**Figure 2C-D**).

No significant reductions in *3R Tau* transcript abundance were observed following treatment (48 h) with **Tau binder** or **RiboTether Recruiter** at concentrations up to 10 μM (**Figure S11A-B**). Similarly, no significant reductions in *4R Tau* reporter RNA abundance were observed under the same conditions (**Figure S11C-D**). The **Tau binder** control did not alter *4R/3R Tau* alternative splicing produced by a related *Tau* pre-mRNA ligand containing the same SRE-binding core^28^. This loss of activity could have resulted from replacement of its *N*-methylcyclohexylamine group^28^ with the longer linker used in **Tau binder** and **Tau-RiboTether**. The PEG2 linker and terminal amide might reduce productive recognition of the SRE through steric or solvation effects according to the established pharmacophore model^28^, which was further suggested by a modest reduction in binding affinity after appending a diazirine module via a short linker to the SRE-binding core^28^.

Collectively, reductions in *Tau-Luc* transcript abundance were found to require both recognition of the *Tau* SRE and recruitment of a functional effector. The concurrent reduction of the *3R* and *4R* reporter transcripts was consistent with a mechanism distinct from modulation of exon 10 splicing alone. A binding site-dependent, induced proximity mechanism involving recruitment of ZFP36 or NANOS3 to the *Tau* reporter transcript is therefore supported.

### Endogenous ZFP36-recruiting Tau RNA degraders using cysteine-reactive ligands

Whether endogenous ZFP36 could be recruited by a chimeric small molecule to reduce *Tau* reporter transcript abundance was evaluated using two cysteine-reactive tryptoline acrylamide ligands, **WX-01-02** and **WX-01-10**^48^. These ligands were identified through reanalysis of publicly available chemoproteomic data in Ramos cells reported by Cravatt and co-workers^48^ (**Figures 3A, left, and S12A-B**). Two sets of stereoprobes, **WX-01-01/02/03/04** and **WX-01-09/10/11/12**, were also examined for stereoselective ZFP36 engagement based on the competition observed at reactive cysteines. The **WX-01-02** achieved >50% ZFP36 engagement at C67, whereas its three stereoisomers (**WX-01-01/03/04**) showed <25% engagement at this position, indicating strong stereoselective engagement by **WX-01-02** (**Figure 3A, right**). In contrast, both **WX-01-10** and **WX-01-12** showed >50% engagement at C67 and C253, respectively, indicating lower stereoselectivity than **WX-01-02** (**Figure 3A, right**).

**Figure 3.**
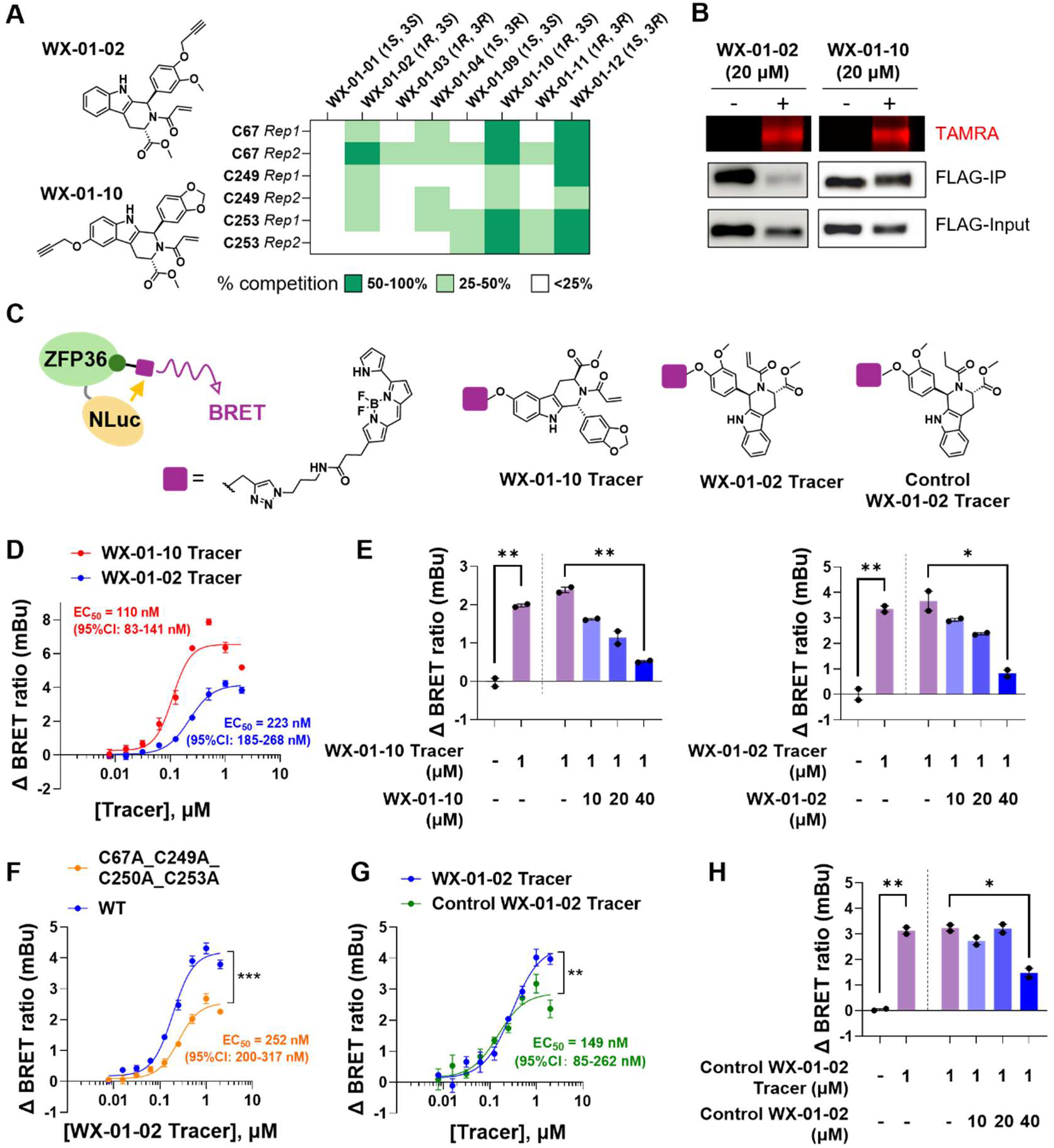
Cellular engagement of ZFP36 by cysteine-reactive tryptoline ligands and a noncovalent analog. **(A)** Structures of **WX-01-10** and **WX-01-02** that were previously identified by cysteine-directed activity-based protein profiling as tryptoline acrylamide probes that engage endogenous ZFP36 in Ramos cells (left)^48^. Heatmap showing ZFP36 engagement by the full stereoprobe sets of **WX-01-10** and **WX-01-02**, with values representing engagement (%) determined by competitive chemical proteomics. **(B)** Engagement of ZFP36-NLuc-FLAG by **WX-01-10** and **WX-01-02** via forced expression of a plasmid in HeLa cells. Following transfection with ZFP36-NLuc-FLAG for 48 h, cells were treated with the indicated probe (20 μM; 3 h). ZFP36-NLuc-FLAG was enriched by FLAG immunoprecipitation, and probe-labeled proteins were conjugated to TAMRA-azide by click chemistry. TAMRA, in-gel fluorescence of probe-labeled ZFP36-NLuc-FLAG following FLAG enrichment; FLAG-IP: anti-FLAG immunoblot following FLAG immunoprecipitation; FLAG-Input: anti-FLAG immunoblot before enrichment (n = 2 biological replicates). **(C)** NanoBRET assay used to measure engagement of ZFP36-NLuc by fluorescent tracers derived from **WX-01-10** or **WX-01-02**, or the noncovalent **control WX-01-02** analog. The structures of the three tracers are shown. **(D)** Concentration dependent NanoBRET responses produced by the **WX-01-10** and **WX-01-02** tracers in live HeLa cells expressing ZFP36-NLuc after incubation for 2 h. Apparent NanoBRET engagement values of 110 nM (95% CI: 83-141 nM) for the **WX-01-10 Tracer** and 223 nM (95% CI: 185-268 nM) for the **WX-01-02 Tracer** were obtained. Because both tracers contain irreversible electrophiles, these values are reported as apparent engagement measurements rather than equilibrium binding constants (n = 3 biological replicates). **(E)** Competitive NanoBRET assays performed in lysates containing ZFP36-NLuc. BRET signals produced by 1 μM of **WX-01-10 Tracer** or **WX-01-02 Tracer** were reduced by the corresponding unlabeled compound at the indicated concentrations (n = 2 biological replicates). **(F)** Engagement of wild-type ZFP36-NLuc and the C67A_C249A_C250A_C253A quadruple ZFP36-NLuc mutant by the **WX-01-02 Tracer** in live HeLa cells after incubation for 2 h. An attenuated NanoBRET response was observed with the four-cysteine mutant relative to wild-type ZFP36-NLuc (n = 3 biological replicates). **(G)** Comparison of ZFP36-NLuc engagement by the covalent **WX-01-02 Tracer** and the noncovalent **control WX-01-02 Tracer** in live HeLa cells after incubation for 2 h. Apparent NanoBRET engagement value was 149 nM (95% CI: 85-262 nM) for the noncovalent **control tracer** (n = 3 biological replicates). **(H)** Competitive NanoBRET assay performed in lysates containing overexpressed ZFP36-NLuc. The signal produced by 1 μM of noncovalent **control WX-01-02 tracer** was reduced by the corresponding unlabeled noncovalent **control WX-01-02** at the indicated concentrations (n = 2 biological replicates). Data are reported as the mean ± SEM. ns, not significant; *p < 0.05, **p < 0.01, and ***p < 0.001, as determined by two-tailed Student’s t-test.

Engagement of cellular ZFP36 was first assessed using click chemistry-based labeling as an orthogonal validation method. HeLa cells overexpressing ZFP36-FLAG were treated with **WX-01-02** or **WX-01-10** (20 μM; 3 h), after which the probes were conjugated to TAMRA-azide and ZFP36-FLAG was enriched by immunoprecipitation. Increased TAMRA signal was detected in the FLAG-enriched samples, supporting covalent engagement of cellular ZFP36 by both probes (**Figure 3B**). Interestingly, both compounds moderately reduced ZFP36 abundance at 20 μM, with a more pronounced effect observed for **WX-01-02**, particularly in the pull-down fraction (**Figure 3B**). This may reflect broader cellular protein engagement that affects protein translation and/or compound-induced conformational changes in the antibody-recognized ZFP36 epitope that reduce antibody recognition.

A NanoBRET assay was also developed to measure ZFP36 engagement in live cells (**Figure 3C**). The ZFP36 was fused at its C-terminus to NanoLuciferase (ZFP36-NLuc, **Figure S1B**), and engagement by fluorescently labeled **WX-01-10** or **WX-01-02** tracers detected through proximity-dependent BRET. Following incubation for 2 h in live HeLa cells, a dose dependent increase in BRET signal was observed for both probes, with apparent NanoBRET engagement values of 110 nM (95% CI: 83-141 nM) for **WX-01-10** and 223 nM (95% CI: 185-268 nM) for **WX-01-02** (**Figure 3D**). Because these probes contain irreversible electrophiles, these values were treated as apparent engagement measurements rather than equilibrium binding constants. In cell lysates containing overexpressed ZFP36-NLuc, dose dependent displacement of the tracer signal was observed when excess **WX-01-02** was added in the presence of 1 μM of **WX-01-02 Tracer** or when excess **WX-01-10** was added in the presence of 1 μM of **WX-01-10 Tracer** (**Figure 3E, left**). These results further support engagement of ZFP36 by both compounds.

Based on modification sites identified in previous activity-based protein profiling studies^48^, four cysteine-to-alanine mutations (C67A, C249A, C250A, and C253A) were introduced into ZFP36-NLuc (**Figure 3A**). Following expression of the mutant protein, an attenuated but saturable NanoBRET response was produced by **WX-01-02 Tracer** relative to wild-type ZFP36-NLuc (**Figure 3F**). This result was consistent with partial contributions from the mutated cysteine residues, while suggesting that additional cysteines or noncovalent interactions might also contribute to engagement.

To evaluate the contribution of the tryptoline scaffold, a noncovalent control analog of **WX-01-02** and its corresponding fluorescent tracer were synthesized by replacing the acrylamide warhead with a propanamide, thereby eliminating covalent reactivity (**Figure 3C, right**). A dose dependent NanoBRET response was retained by the noncovalent control tracer, with an apparent engagement value comparable to that of the covalent **WX-01-02 Tracer** (149 nM, 95% CI: 85-262 nM) but with a modestly reduced maximal response (2.5 mBu vs. 4.2 mBu). The BRET signal was also displaced by excess noncovalent **control WX-01-02** (**Figure 3G**). Thus, a substantial contribution to ZFP36 engagement was provided by the tryptoline scaffold, and engagement was not dependent exclusively on the acrylamide warhead. Collectively, **WX-01-10** and **WX-01-02** were identified as cellular ZFP36 ligands suitable for evaluation as ZFP36-recruiting modules in *Tau*-targeting chimeric compounds.

To evaluate functional activity, the *Tau* RNA-binding moiety was conjugated to **WX-01-10** or **WX-01-02** through a PEG2 linker, yielding **Tau-WX-01-10** and **Tau-WX-01-02**, respectively (**Figure 4A**). In HeLa cells transfected with the *Tau-Luc* mini-gene, dose dependent reductions in Tau-Luc reporter output and *Tau* reporter transcript abundance were produced by **Tau-WX-01-02** (**Figures 4B-C**). At 20 μM, maximal reductions of 32 ± 5% in reporter output, 38 ± 8% in *4R Tau-Luc* RNA, 36 ± 7% in *3R Tau-Luc* RNA, and 26 ± 3% in total *Tau-Luc* transcript abundnace were observed. Interestingly, attaching the noncovalent **control WX-01-02** to the *Tau* RNA-binding moiety via the same PEG2 linker did not afford a functional degrader capable of decreasing Tau-Luc reporter levels (**Tau-control-WX-01-02**, **Figure S13**), indicating the importance of covalent acrylamide warhead to effective ZFP36 engagement.

**Figure 4.**
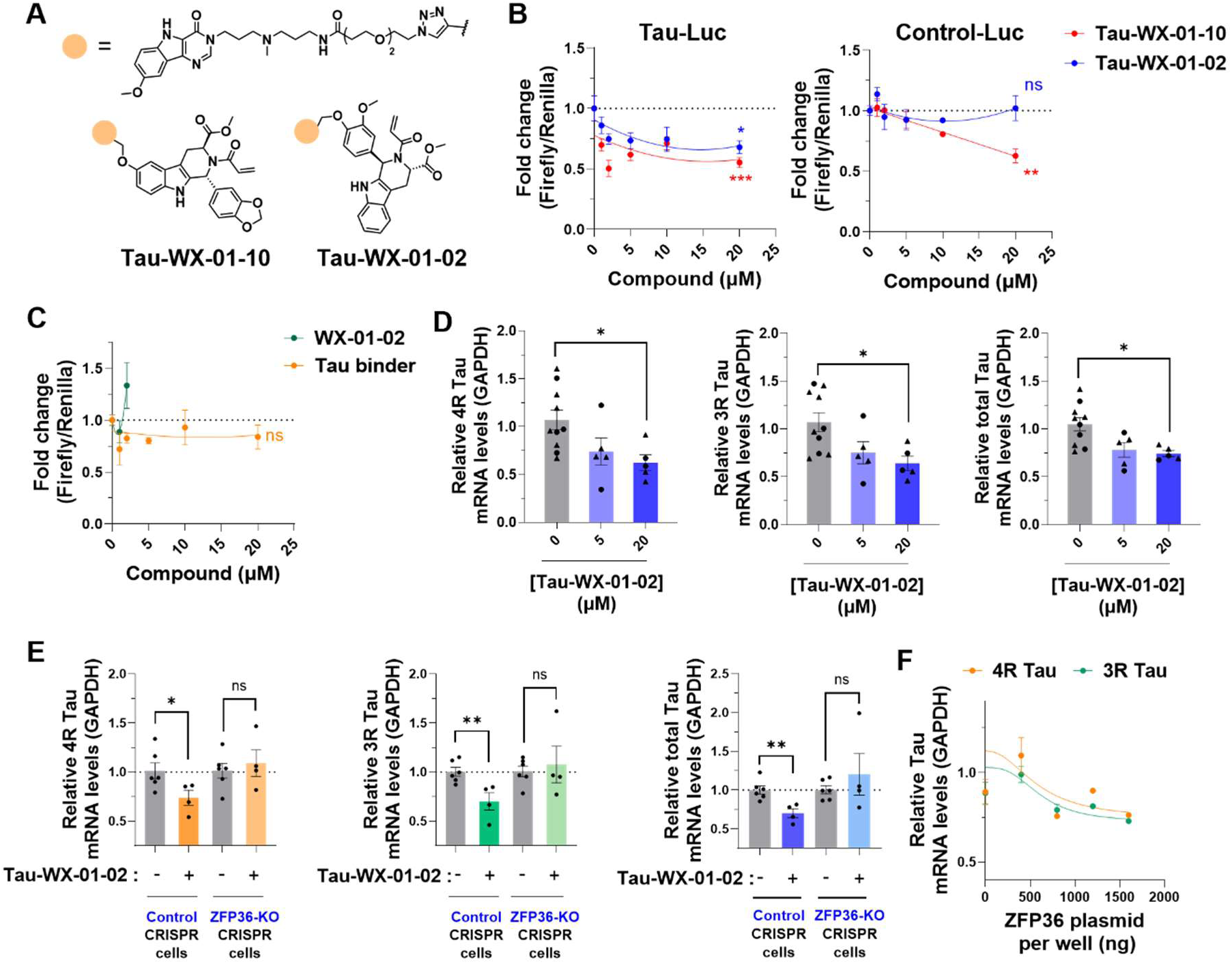
ZFP36-recruiting chimeric compounds reduce *Tau-Luc* transcript abundance in a ZFP36-dependent manner. **(A)** Structures of **Tau-WX-01-10** and **Tau-WX-01-02**. The Tau pre-mRNA-binding module was conjugated to the cysteine-reactive ZFP36 ligands **WX-01-10** or **WX-01-02** via a PEG2-containing linker using click chemistry. **(B)** Concentration dependent effects of **Tau-WX-01-10** and **Tau-WX-01-02** on Tau-Luc and Control-Luc reporter output. HeLa cells transfected with the *Tau* mini-gene reporter or a control firefly luciferase reporter lacking the *Tau* sequence were treated with the indicated compounds for 48 h. Firefly luminescence was normalized to Renilla luminescence and expressed relative to the corresponding vehicle-treated condition (0.2% (v/v) DMSO). **Tau-WX-01-02** reduced Tau-Luc output without significantly affecting Control-Luc, whereas **Tau-WX-01-10** reduced both reporters, indicating a *Tau*-sequence-independent component of its activity. **(C)** Effects of the monofunctional **Tau binder** and **WX-01-02** controls on Tau-Luc reporter output. Cells were treated with **Tau binder** at concentrations up to 20 μM or **WX-01-02** at concentrations up to 2 μM for 48 h. **(D)** Effect of **Tau-WX-01-02** on *4R*, *3R*, and total *Tau-Luc* transcript abundance. HeLa cells transfected with the *Tau* mini-gene were treated with **Tau-WX-01-02** at the indicated concentrations for 48 h. Transcript levels were measured by RT-qPCR, normalized to *GAPDH*, and expressed relative to vehicle-treated cells (0.2% (v/v) DMSO). Circles and triangles denote measurements obtained from two independent experiments. **(E)** Dependence of **Tau-WX-01-02** activity on endogenous ZFP36. Relative *4R*, *3R*, and total *Tau-Luc* transcript abundances were measured in HeLa cells transduced with a non-targeting CRISPR control or ZFP36-targeting CRISPR and treated with vehicle (0.2% (v/v) DMSO) or 20 μM of **Tau-WX-01-02** for 48 h. Transcript levels were normalized to *GAPDH* and expressed relative to the corresponding vehicle-treated cells (0.2% (v/v) DMSO). **(E)** Restoration of **Tau-WX-01-02** activity following re-expression of ZFP36 in ZFP36-depleted cells. Increasing amounts of the ZFP36 expression plasmid were transfected per well, and relative *4R* and *3R Tau-Luc* transcript levels were measured by RT-qPCR and normalized to *GAPDH*. Curves are included as visual guides; no potency values were derived from these fits. Data are reported as the mean ± SEM, with individual measurements shown where applicable. ns, not significant; *p < 0.05, **p < 0.01, and ***p < 0.001, as determined by two-tailed Student’s t-test.

No reduction in the control firefly luciferase reporter lacking the *Tau* sequence was produced by **Tau-WX-01-02** at concentrations up to 20 μM. In contrast, comparable reductions were produced by **Tau-WX-01-10** in both the control firefly luciferase system (37.3 ± 5.6%; **Figure 4B, right**) and the Tau-Luc luciferase system (44.6 ± 4.0%; **Figure 4B, left**). Greater dependence on the *Tau* sequence was therefore demonstrated by **Tau-WX-01-02**, and thus this compound was selected for subsequent mechanistic studies.

No reductions in Tau-Luc reporter output (**Figure 4D**) nor *Tau-Luc* transcript abundance (**Figure S14A**) were observed following treatment with the **Tau binder** alone or with **WX-01-02** alone at tolerated concentrations (0-20 μM for **Tau binder**, 0-2 μM for **WX-01-02**). Treatment with **WX-01-02** for 48 h was tolerated in HeLa cells only at concentrations up to 2 μM, whereas no loss of cell viability was observed following treatment with **Tau-WX-01-02** at concentrations up to 20 μM (**Figure S14B**). Thus, cellular tolerability was altered by incorporation of **WX-01-02** into the chimeric compound, although the basis for this difference was not determined.

ZFP36-dependence was evaluated by CRISPR-mediated depletion of endogenous ZFP36. An approximately 90% reduction in ZFP36 protein abundance was achieved (**Figure S15A-B**), and the reduction of *Tau-Luc* transcript abundance by **Tau-WX-01-02** was abolished relative to cells treated with a non-targeting CRISPR control (**Figure 4E**). When ZFP36 was re-expressed in the ZFP36-depleted cells, sensitivity to **Tau-WX-01-02** was restored (**Figure 4F**). Dependence of **Tau-WX-01-02** activity on endogenous ZFP36 was therefore supported by both loss-of-function and re-expression experiments. Together with the chemical controls studies described above, these results support a requirement for both RNA engagement and ZFP36 recruitment and thus an induced-proximity mechanism.

### Improved potency of Tau-ZFP36 degraders by linker optimization

Although reduction of *Tau-Luc* transcript abundance was achieved with the PEG2-linked **Tau-WX-01-02** compound (**Tau-PEG2-WX-01-02**), a relatively high concentration of 20 μM was required to produce moderate activity in HeLa cells. Because degrader activity can be strongly influenced by linker architecture, as previously demonstrated for RNase L-recruiting degraders^49^, a focused series of **Tau-[linker]_x_-WX-01-02** chimeras was synthesized. Linker length, hydrophobicity, and conformational rigidity were varied across the series (**Figure 5A**). All chimeras affected cell viability by <25% after treatment with up to 20 μM for 48 h (**Figure S16**).

**Figure 5.**
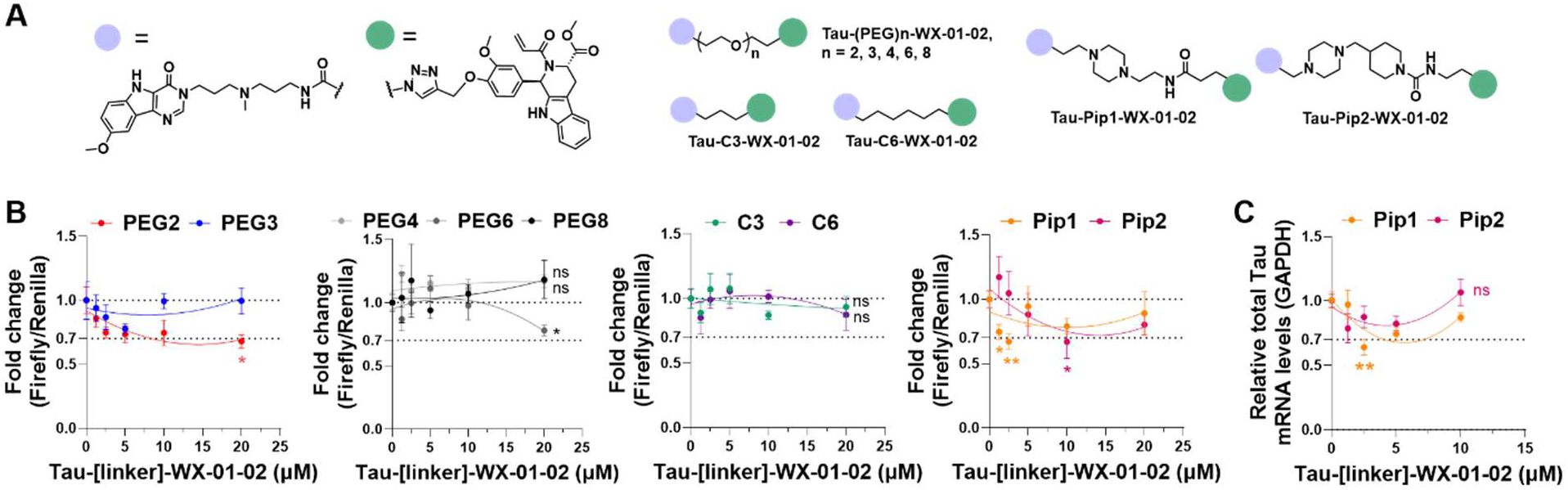
Potencies of the ZFP36-recruiting Tau degraders following linker optimization. **(A)** Structures of the **Tau-[linker]-WX-01-02** chimeric compounds. The focused series comprised two alkyl linkers (C3 and C6), two piperazine-containing linkers (Pip1 and Pip2), and five polyethylene glycol linkers containing two, three, four, six, or eight ethylene glycol units (PEG2, PEG3, PEG4, PEG6, and PEG8, respectively). Note that **Tau-PEG2-WX-01-02 is the same as Tau-WX-01-02 in** Figure 4**. (B)** Effects of the nine **Tau-[linker]-WX-01-02** compounds on Tau-Luc reporter output. HeLa cells transfected with the *Tau* mini-gene reporter plasmid were treated with the indicated compounds for 48 h. Firefly luminescence was normalized to Renilla luminescence and expressed relative to the corresponding vehicle-treated cells (0.2% (v/v) DMSO) (n = 5 biological replicates). **(C)** Effects of **Tau-Pip1-WX-01-02** and **Tau-Pip2-WX-01-02** on total *Tau-Luc* transcript abundance following treatment for 48 h. Transcript levels were measured by RT-qPCR, normalized to *GAPDH*, and expressed relative to the corresponding vehicle-treated condition (0.2% (v/v) DMSO) (n = 3 biological replicates). For **(B)** and **(C)**, curves are included as visual guides; no potency values were derived from these fits. Data are reported as the mean ± SEM, with individual measurements shown where applicable. ns, not significant; *p < 0.05 and **p < 0.01, as determined by two-tailed Student’s t-test.

The greatest improvement in potency was produced by incorporation of piperazine linker 1 into **Tau-Pip1-WX-01-02**. Following treatment with a 2.5 μM dose for 48 h, reductions of 33 ± 6% in Tau-Luc reporter output (p < 0.01; **Figure 5B**) and 36 ± 6% in total *Tau-Luc* transcript abundance (p < 0.01; **Figure 5C**) were observed. In contrast, there was no significant reduction in Tau-Luc reporter activity upon treatment with 2.5 μM of first-generation **Tau-PEG2-WX-01-02 (Figure 5B)**. Improved activity was also observed with a second piperazine-containing compound, **Tau-Pip2-WX-01-02**. At the 10 μM dose, Tau-Luc reporter output was reduced by 33 ± 13% (p < 0.05; vs. 25 ± 4% for 20 μM of **Tau-PEG2-WX-01-02**; **Figure 5B**), although no significant reduction in *Tau-Luc* transcript abundance was detected (**Figure 5C**). In contrast, no appreciable improvement in activity was produced by PEG-based linkers longer than PEG2 (**Figure 5B**). Quantification by ddPCR further showed that the reduction in total *Tau-Luc* transcript abundance produced by 2.5 μM of **Tau-Pip1-WX-01-02** was comparable to that produced by 20 μM of **Tau-PEG2-WX-01-02** (**Figure S17**). Thus, an approximately 8-fold increase in potency was achieved through incorporation of piperazine linker 1.

For **Tau-Pip1-WX-01-02**, an apparent hook effect is observed at higher concentrations for both reporter output and RNA abundance (**Figure 5B-C**). This nonmonotonic concentration response affords a relatively narrow activity window. Similar responses have been reported for irreversible covalent proteolysis-targeting chimeras containing related acrylamide warheads at low micromolar concentrations^50^. Collectively, these results demonstrated that the potency and concentration-dependent activity of the ZFP36-recruiting Tau degrader are strongly influenced by linker composition, which could affect geometry, cellular uptake, and subcellular localization, among other factors.

## DISCUSSION

The **RiboTether** platform enables discovery of productive RNA-effector relationships, and protein ligandability maps enable these relationships to be converted into endogenous effector RiboTACs. By combining a library of FKBP12^F36V^-tagged effector proteins with a cell permeable heterobifunctional recruiter, individual RBPs could be systematically evaluated for their ability to alter the abundance of a particular RNA target. ZFP36 and NANOS3 were identified as recruitable effectors that reduced both *3R* and *4R Tau-Luc* transcript abundance. This finding was subsequently translated to recruitment of endogenous ZFP36 using a cysteine-reactive ligand, with activity further improved through linker optimization. Thus, a general workflow was established in which productive effector function can first be identified in a programmable tethering system and subsequently translated into a chimera that recruits an endogenous effector protein.

The importance of effector recruitment is supported by previous studies demonstrating that occupancy of an RNA by a small molecule is often insufficient to produce a biological response^13, 51, 52^. RNA-binding compounds with little or no intrinsic activity have been converted into active degraders through recruitment of RNase L, with RNA degradation abolished upon depletion of the recruited effector^13^. Similar effector dependence was observed for a *MYC*-targeting RiboTAC in multiple myeloma, where reduction of *MYC* RNA and antiproliferative activity required both expression of the target RNA and functional RNase L^53, 54^. These studies illustrate a central feature of induced-proximity approaches to RNA: productive activity requires not only engagement of a ligandable RNA structure but also recruitment of an effector protein whose localization, abundance, and mechanism are compatible with the targeted transcript. The **RiboTether** platform provides a systematic means to identify such productive RNA-effector relationships before development of small molecules that recruit the endogenous effector protein.

Expansion of the recruitable effector repertoire should increase the range of RNAs and cellular compartments that can be addressed. RNase L has been successfully recruited to cytoplasmic RNA targets^55–57^, whereas LC3B recruitment has been used to induce autophagy-dependent degradation of *collagen type XV alpha 1 chain* (*COL15A1)* mRNA^16^. In the latter study, Unbiased RNA Degrader Identification (URID) was used to integrate transcriptome-wide RNA engagement with RNA-seq measurements of transcript depletion, allowing productive RNA-binding and effector recruitment events to be distinguished from nonproductive occupancy. In the present study, ZFP36 and NANOS3 were identified as effectors capable of acting on an RNA element located within nuclear pre-mRNA. Collectively, these findings indicate that different effectors can provide distinct substrate preferences, mechanisms, and subcellular access. A broader effector palette could therefore be used to match an RNA target with an effector whose localization, expression, and mechanism are compatible with the targeted transcript.

Although RNA destabilization was evaluated here, the tethering platform need not be limited to RNA decay. Libraries of tagged RBPs could be screened for effects on RNA stabilization, translation, localization, processing, or expression. Epitranscriptomic writers, erasers, and readers could also be recruited to determine whether installation, removal, or recognition of RNA modifications can be chemically controlled. Reporter outputs could be adapted to measure RNA abundance, translation, splice-isoform production, localization, or modification dependent signaling. The model system could also be expanded using RNA aptamers or other orthogonal RNA-ligand pairs, allowing defined RNA recruitment sites and geometries to be installed in different reporter transcripts. Such systems would permit effector functions to be systematically evaluated independently of the availability of a ligand for a particular endogenous RNA.

The discovery of recruitable effectors could also be accelerated by integration with proteome-wide ligand-discovery studies. Chemical proteomics maps^48, 58–60^ can be used to identify ligandable sites within RBPs and other RNA-regulatory proteins, while transcriptome-wide RNA-binding methods^55^ can be used to identify the RNAs and structural sites engaged by small molecules. Integration of these datasets with approaches such as URID could provide a systematic route for pairing RNA-binding modules with recruitable effector ligands, after which linker composition and recruitment geometry could be optimized. As additional RNA-targeting small molecules are discovered through transcriptome-wide studies, an expanded collection of effector ligands should allow otherwise silent RNA-binding events to be converted into changes in RNA stability, processing, modification, localization, or translation.

A limitation of the present ZFP36-recruiting compounds is that the ZFP36 binders are only modestly selective and potent **(Figure 3A, Supplementary Tables S1 and S2)**. Broad proteome remodeling, protein aggregation, proteasome activation, and stress-granule formation have been observed with less selective cysteine-reactive fragments^61^. Although overt cytotoxicity was not observed with the optimized Tau-ZFP36 chimera under the conditions examined here, the absence of cytotoxicity does not exclude more subtle electrophile-associated stress or off-target protein engagement^62^. These possibilities will need to be carefully addressed during future optimization. Improved potency, moderated electrophile reactivity, proteome-wide assessment of target engagement, and monitoring of cellular stress responses should be considered^63^. Ultimately, development of selective noncovalent ZFP36 recruiters would be preferred and could provide a clearer pharmacological path while retaining the induced-proximity mechanism established here.

## CONCLUSION

The present study establishes a general strategy in which chemically controlled tethering is used to discover productive RNA-effector relationships before development of recruiters for endogenous effector proteins. As the chemical space of RNA-binding molecules and protein ligands continues to expand, increasingly diverse aspects of RNA biology should become accessible to small molecule control^64, 65^. A broader repertoire of recruitable effectors could ultimately enable RNA fate to be manipulated across cellular compartments through targeted degradation, stabilization, expression enhancement, processing, localization, and epitranscriptomic modification^66, 67^. Future implementations of this platform could incorporate direct measurements of RNA fate, including long-read and direct RNA sequencing approaches such as nanopore sequencing^68^, to systematically identify effector proteins capable of producing these distinct outcomes. Together, these advances expand RiboTAC technology beyond a limited set of known ribonucleases toward a more general framework for chemically programming RNA function through induced proximity.

## Supporting information

Tau-RiboTether_supplementary_information

## DATA AVAILABILITY

All data can be found in Supplemental Documents associated with this manuscript.

## ACKNOWLEDGMENT

This work was supported by the Tau Consortium and the Rainwater Charitable Fund (to M.D.D.) and the National Institute of Neurological Disorders and Stroke R35 grant (R35 NS116846 to M.D.D.). We thank Jessical L. Childs-Disney for advice and critical review of the manuscript.

## Notes

### Competing Interest Statement

M.D.D. is a co-founder of Ribonaut Therapeutics.

