## Supplementary material for "Programmable Recruitment of RNA-Binding Proteins Enables Small Molecule-Directed Destabilization of Nuclear Pre-mRNA": Tau-RiboTether_supplementary_information

#### TABLE OF CONTENTS

##### Supplementary Tables


| Item | Title | Page |
|---|---|---|
| Table S1 | FKBP12$^{F36V}$-fused RNA-binding proteins used in this study | S4 |
| Table S2 | Primer and oligonucleotide sequences used in this study | S5 |


##### Supplementary Figures


| Item | Title | Page |
|---|---|---|
| Figure S1 | Validation of expression of NLuc-fused proteins used in NanoBRET assays | S7 |
| Figure S2 | Validation of expression of FKBP12$^{F36V}$-fused RNA-binding proteins | S8 |
| Figure S3 | FKBP12$^{F36V}$ cellular engagement by ortho-AP1867 and Tau binder controls | S9 |
| Figure S4 | Potential ZFP36-binding sites near the Tau pre-mRNA SRE | S10 |
| Figure S5 | Firefly and Renilla luminescence following **Tau-RiboTether** treatment | S11 |
| Figure S6 | Signal-to-noise analysis of the Tau-Luc reporter system | S12 |
| Figure S7 | Dose-response analysis of the FKBP12$^{F36V}$-tagged effector screen | S13 |
| Figure S8 | Raw $C_t$ values and no-RT controls for RT-qPCR measurements | S14 |
| Figure S9 | **Tau-RiboTether**-mediated reduction of *Tau-Luc* transcript abundance by ZFP36 and NANOS3 | S15 |
| Figure S10 | Effect of **Tau-RiboTether** following ablation of the Tau binder recognition site | S16 |
| Figure S11 | Effects of **Tau binder** and **RiboTether Recruiter** controls on *Tau-Luc* transcript abundance | S17 |
| Figure S12 | Structures of **WX-01-02** and **WX-01-10** stereoisomers | S18 |
| Figure S13 | Effects of **Tau-control-WX-01-02** on Tau-Luc reporter output | S19 |
| Figure S14 | Effects of **Tau binder** and **WX-01-02** on *Tau-Luc* transcript abundance and cell viability | S20 |
| Figure S15 | Validation of ZFP36 CRISPR depletion | S21 |
| Figure S16 | Effects of **Tau-[linker]-WX-01-02 compounds** on cell viability | S22 |
| Figure S17 | ddPCR quantification of *Tau-Luc* transcript abundance following treatment with ZFP36-recruiting chimeras | S23 |


##### Materials and Methods


| Title | Page |
|---|---|
| Plasmid construction | S24 |
| Cell culture | S24 |

| Title | Page |
| --- | --- |
| Luciferase assay | S24 |
| Measurement of *Tau-Luc* transcript abundance by RT-qPCR | S25 |
| Validation of fusion protein expression by Western blotting | S25 |
| Gel-based ABPP validation of ZFP36 engagement | S26 |
| ZFP36 NanoBRET assay | S28 |
| FKBP12$^{F36V}$ NanoBRET assay | S29 |
| Construction of control and ZFP36 CRISPR knockout cells | S30 |
| Proteomics data reanalysis | S30 |

**Synthesis and Characterization of Compounds**

| Item | Title | Page |
| --- | --- | --- |
| — | Abbreviations | S31 |
| — | General synthetic methods | S31 |
| Scheme 1 | Synthesis of **Tau-RiboTether** | S32 |
| Scheme 2 | Synthesis of **Tau-binder** | S33 |
| Scheme 3 | Synthesis of control **WX-01-02** | S34 |
| Scheme 4 | Synthesis of **WX-01-10 Tracer** | S35 |
| Scheme 5 | Synthesis of **WX-01-02 Tracer** | S36 |
| Scheme 6 | Synthesis of **Control WX-01-02 Tracer** | S37 |
| Scheme 7 | Synthesis of **Tau-WX-01-10** | S38 |
| Scheme 8 | Synthesis of **Tau-WX-01-02** | S39 |
| Scheme 9 | Synthesis of **Tau-[linker]-WX-01-02** | S40 |

**Compound Characterization**

| Title | Page |
| --- | --- |
| $^{1}$H NMR, $^{13}$C NMR, analytical HPLC, and HR-MS spectra | S42 |

**References**

| Title | Page |
| --- | --- |
| References | S71 |

#### TABLES & FIGURES

**Table S1. FKBP$^{F36V}$ fused RNA-binding proteins used in this study.**

| | Protein | Accession No. | Function | Subcellular location* | Expressible ? |
|---|---|---|---|---|---|
| 1 | ZFP36 | NM_003407.3 | RNA Destabilizer | Nuc & Cyt | Yes |
| 2 | CNOT7 | NM_001322090.1 | RNA Destabilizer | Nuc & Cyt | Yes |
| 3 | PARN | NM_002582.3 | RNA Destabilizer | Nuc & Cyt | Yes |
| 4 | NANOS3 | NM_001098622.2 | RNA Destabilizer | Nuc & Cyt | Yes |
| 5 | XRN2 | NM_012255.5 | RNA Destabilizer | Nuc | Yes |
| 6 | TOB1 | NM_001243877.1 | RNA Destabilizer | Nuc & Cyt | Yes |
| 7 | SNRPA | NM_004596.5 | RNA Destabilizer | Nuc | Yes |
| 8 | MEX3C | NM_016626.5 | RNA Destabilizer | Nuc & Cyt | Yes |
| 9 | DDX6 | NM_001257191.2 | RNA Destabilizer | Nuc & Cyt | Yes |
| 10 | CNOT1 | NM_016284.4 | RNA Destabilizer | Nuc & Cyt | No |
| 11 | LC3B | NM_022818 | RNA Destabilizer | Cyt | Yes[1] |
| 12 | EXOSC2 | BC000747.1 | RNA Destabilizer | Nuc & Cyt | No |
| 13 | EXOSC10 | BC073788.1 | RNA Destabilizer | Nuc & Cyt | No |
| 14 | RNase L | NM_021133.3 | RNA Destabilizer | Cyt | Yes |
| 15 | SMG6 | NM_017575.4 | RNA Destabilizer | Nuc & Cyt | No |
| 16 | SMG7 | NM_173156.2 | RNA Destabilizer | Nuc & Cyt | No |
| 17 | DIS3 | NM_014953.4 | RNA Destabilizer | Nuc & Cyt | No |
| 18 | PAPOLB | NM_020144.4 | RNA Stabilizer | Nuc | Yes |
| 19 | CLK3 | XM_017021909.1 | RNA Stabilizer | Nuc & Cyt | Yes |
| 20 | AIMP1 | NM_001142415.1 | RNA Stabilizer | Nuc & Cyt | Yes |
| 21 | MTDH | NM_178812.3 | RNA Stabilizer | Nuc & Cyt | Yes |

* **"**Nuc" denotes nuclear location, and "Cyt" denotes cytosolic localization, according to UniProt[2].

**Table S2. Primer and other oligonucleotides sequences used in this study.** **"**Fwd" denotes a forward primer and "Rev" denotes a reverse primer.

| **Oligonucleotide** | **Sequence (5' to 3')** | **Experiment** |
|---|---|---|
| *4R Tau-mini* Fwd | GAGGCGGGAAGGTGCAGATAATTAATAAGA | qPCR |
| *3R Tau-mini* Fwd | CAGCCGGGAGGCGGGAAGGTGCAAATAG | qPCR |
| *4R/3R Tau-mini* Rev | GCCTTATGCAGTTGCTCTCC | qPCR |
| Total *Tau-mini* Fwd | ATCCGCTGGAAGATGGAACC | qPCR |
| Total *Tau-mini* Rev | TAGCTTCTGCCAACCGAACG | qPCR |
| *GAPDH* Fwd | TGCACCACCAACTGCTTAG | qPCR |
| *GAPDH* Rev | GATGCAGGGATGATGTTC | qPCR |
| *ZFP36* sgRNA1 | **mC*mC*mA*rCrArArCrCrCrUrArGrCrGrArArGrArCrCr**GrUrUrUrArGrArGrCrUrArGrArArArUrArGrCrArArGrUrUrArArArArUrArArGrGrCrUrArGrUrCrCrGrUrUrArUrCrArArCrUrUrGrArArArArArGrUrGrGrCrArCrCrGrArGrUrCrGrGrUrGrCmU*mU*mU*rU | CRISPR knockout |
| *ZFP36* sgRNA2 | **mG*mC*mU*rArCrArArGrArCrUrGrArGrCrUrArUrGrU**rGrUrUrUrArGrArGrCrUrArGrArArArUrArGrCrArArGrUrUrArArArArUrArArGrGrCrUrArGrUrCrCrGrUrUrArUrCrArArCrUrUrGrArArArArArGrUrGrGrCrArCrCrGrArGrUrCrGrGrUrGrCmU*mU*mU*rU | CRISPR knockout |
| Non-targeting sgRNA1 | **mG*mG*mA*rGrUrUrArArGrGrCrCrUrCrGrUrCrUrArG**rGrUrUrUrArGrArGrCrUrArGrArArArUrArGrCrArArGrUrUrArArArArUrArArGrGrCrUrArGrUrCrCrGrUrUrArUrCrArArCrUrUrGrArArArArArGrUrGrGrCrArCrCrGrArGrUrCrGrGrUrGrC | CRISPR knockout |
| Non-targeting sgRNA2 | **mG*mU*mG*rCrGrGrGrGrGrCrArUrGrGrCrCrCrCrGrCr**GrUrUrUrArGrArGrCrUrArGrArArArUrArGrCrArArGrUrUrArArArArUrArArGrGrCrUrArGrUrCrCrGrUrUrArUrCrArArCrUrUrGrArA | CRISPR knockout |

| | | |
|---|---|---|
| | rArArArGrUrGrGrCrArCrCrGrArGrUr CrGrGrUrGrC | |
| *ZFP36* Fwd | CGCCACCCCAAATACAAGAC | qPCR |
| *ZFP36* Rev | GTCTTCGCTAGGGTTGTGGA | qPCR |
| Oligonucleotides were purchased from Integrated DNA Technologies (IDT).<br>"*" denotes phosphorothioate linkage.<br>"r" denotes a ribose sugar (RNA nucleoside).<br>**Bold** text denotes spacer sequences in sgRNAs. | | |

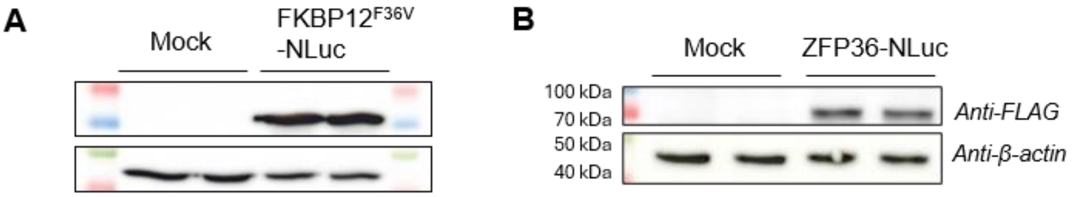


**Figure S1. Validation of expression of NLuc-fused proteins used in NanoBRET assays. (A)** Expression of FKBP12$^{F36V}$-NLuc protein in HeLa cells, measured 48 h post-transfection. **(B)** Expression of ZFP36-NLuc fusion protein in HeLa cells, measured 48 h post-transfection. Immunoblotting was performed using an anti-FLAG, and anti-β-actin primary antibodies.

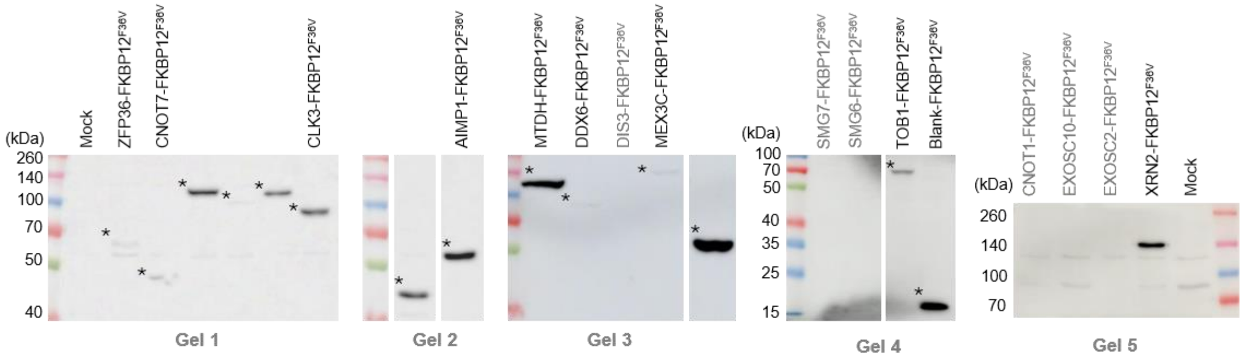


**Figure S2. Validation of the expression of FKBP$^{F36V}$-fused RNA-binding proteins in HeLa cells.** Immunoblotting was performed using an anti-FLAG primary antibody. Grey indicates no detectable expression; black indicates validated overexpression relative to a mock-transfected control.

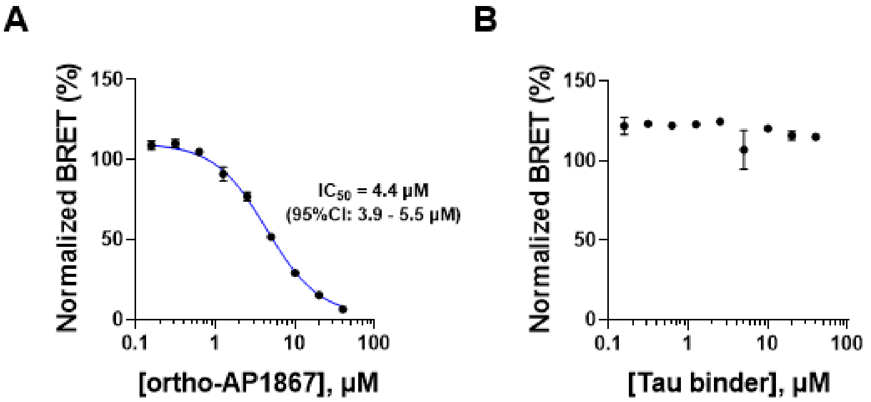


**Figure S3. FKBP12$^{F36V}$ cellular engagement by ortho-AP1867 and Tau binder was measured using competitive NanoBRET. (A)** Engagement of the **ortho-AP1867** tracer was competitively displaced by **ortho-AP1867** with an $IC_{50}$ of 4.4 μM (95% CI, 3.9-5.5 μM; middle). (B) **Tau binder** did not displace **ortho-AP1867 tracer** with treatment up to 40 μM. All data are reported as the mean ± SEM (n = 3 biological replicates).

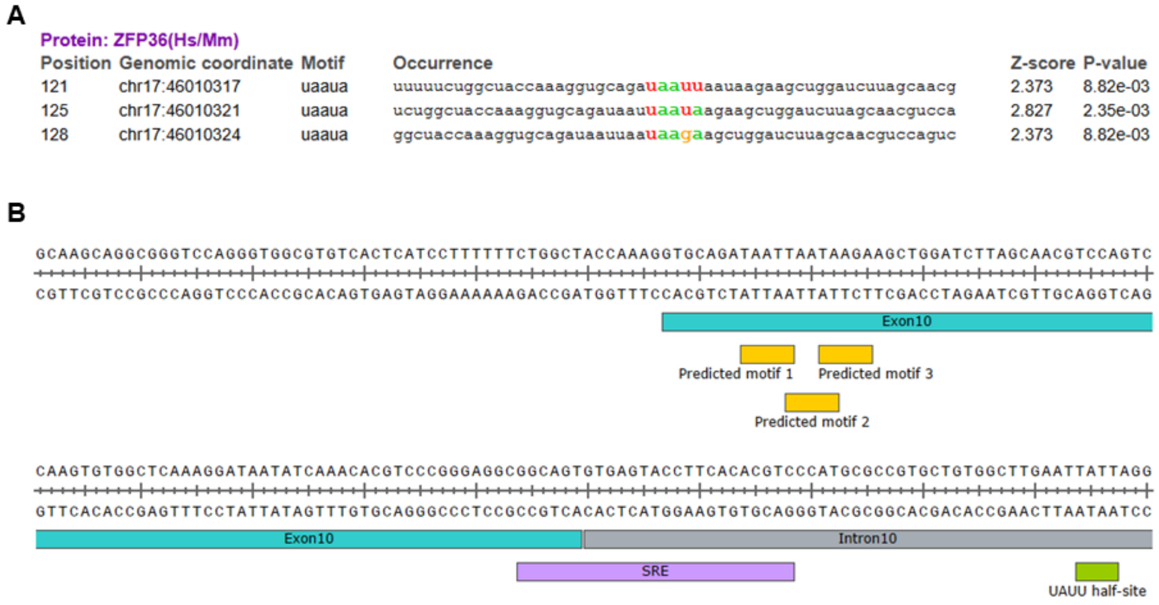


**Figure S4. Potential ZFP36-binding sites near SRE region of *Tau* pre-mRNA. (A)** RBPmap[3] predicted three potential ZFP36 binding motifs when inputting a 425 nt fragment from *MAPT* gene nearby the SRE (genomic coordinates 115,644-116,068). Prediction parameters: genome: human (hg38); selected motifs: all human/mouse motifs; stringency level: default; conservation filter: off. **(B)** The three predicted ZFP36 binding motifs (yellow) are located within exon 10 (~70 nt upstream of SRE) of *Tau* pre-mRNA. Another known ZFP36 RNA regulatory element (RRE), a single UAUU half-site[4], is located 25 nt downstream of *Tau* SRE.

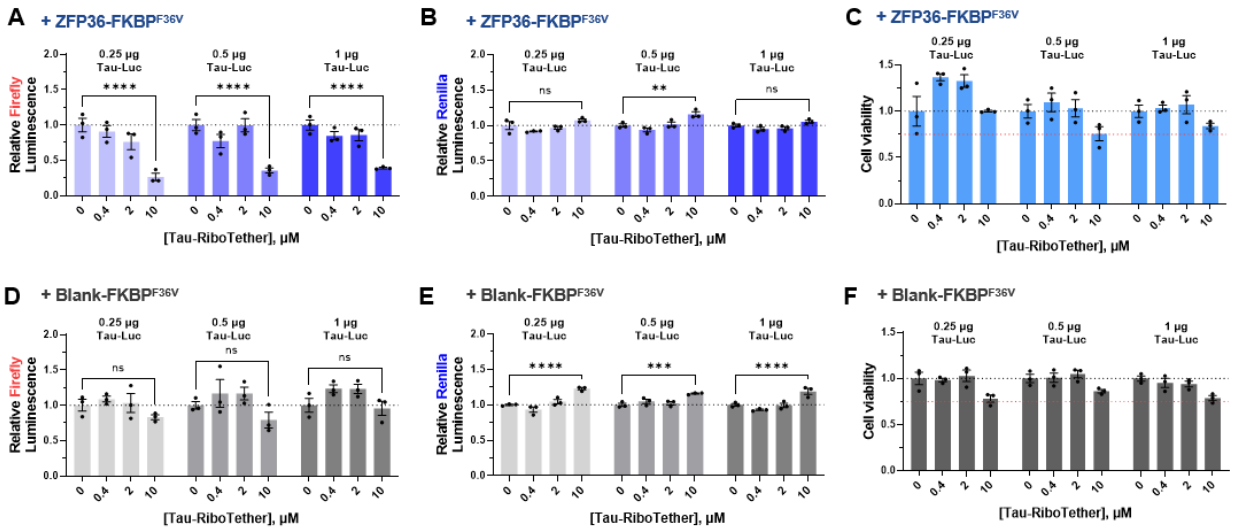


**Figure S5. Relative firefly and Renilla luminescence signal following Tau-RiboTether treatment. (A)** Effect of **Tau-RiboTether** on relative firefly luminescence in HeLa cells co-transfected with a plasmid encoding ZFP36-FKBP12$^{F36V}$ and varying amounts the *Tau-Luc* plasmid (a previously described mini-gene construct[5, 6]; 0.25 μg, 0.5 μg, or 1 μg), without normalization to Renilla signal. **(B)** Effect of **Tau-RiboTether** on relative Renilla luminescence in HeLa cells co-transfected with a plasmid encoding ZFP36-FKBP12$^{F36V}$ and varying amounts the *Tau-Luc* plasmid (0.25 μg, 0.5 μg, or 1 μg). **(C)** Effect of **Tau-RiboTether** on the viability of HeLa cells transfected with 0.25, 0.5, or 1 μg of *Tau-Luc* plasmid, as well as Renilla plasmid and ZFP36-FKBP12$^{F36V}$ plasmid, followed by a 48 h treatment period. **(D)** Effect of **Tau-RiboTether** on relative firefly luminescence in HeLa cells co-transfected with a plasmid encoding Blank-FKBP12$^{F36V}$ and varying amounts of the *Tau-Luc* plasmid (0.25 μg, 0.5 μg, 1 μg), without normalization to Renilla signal. **(E)** Effect of **Tau-RiboTether** on relative Renilla luminescence in HeLa cells co-transfected with a plasmid encoding Blank-FKBP12$^{F36V}$ and varying amounts of *Tau-Luc* plasmid (0.25 μg, 0.5 μg, 1 μg). **(F)** Effect of **Tau-RiboTether** on the viability of cells expressing Blank-FKBP12$^{F36V}$ after a 48 h treatment period. "0" indicates vehicle-treated cells (0.1% (v/v) DMSO). ns, non-significant; $^{**}p < 0.01$, $^{***}p < 0.001$, $^{****}p < 0.0001$, as determined by two-way ANOVA test. All data are reported as the mean ± SEM (n = 3 biological replicates).

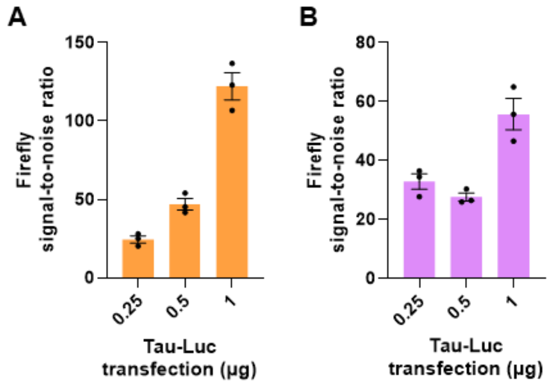


**Figure S6. Signal-to-noise ratio of Firefly luminescence following co-transfection of FKBP12$^{F36V}$ effector protein plasmid and *Tau-Luc* plasmid. (A)** Co-transfection of *ZFP36-FKBP12$^{F36V}$* plasmid and 1 µg of *Tau-Luc* plasmid showed highest signal-to-noise ratio of firefly luminescence, as determined by comparing luminescence to mock-transfected cells. **(B)** Co-transfection of *Blank-FKBP12$^{F36V}$* plasmid and 1 µg of *Tau-Luc* plasmid showed the highest signal-to-noise ratio of Firefly luminescence. All data are reported as the mean ± SEM (n = 3 biological replicates).

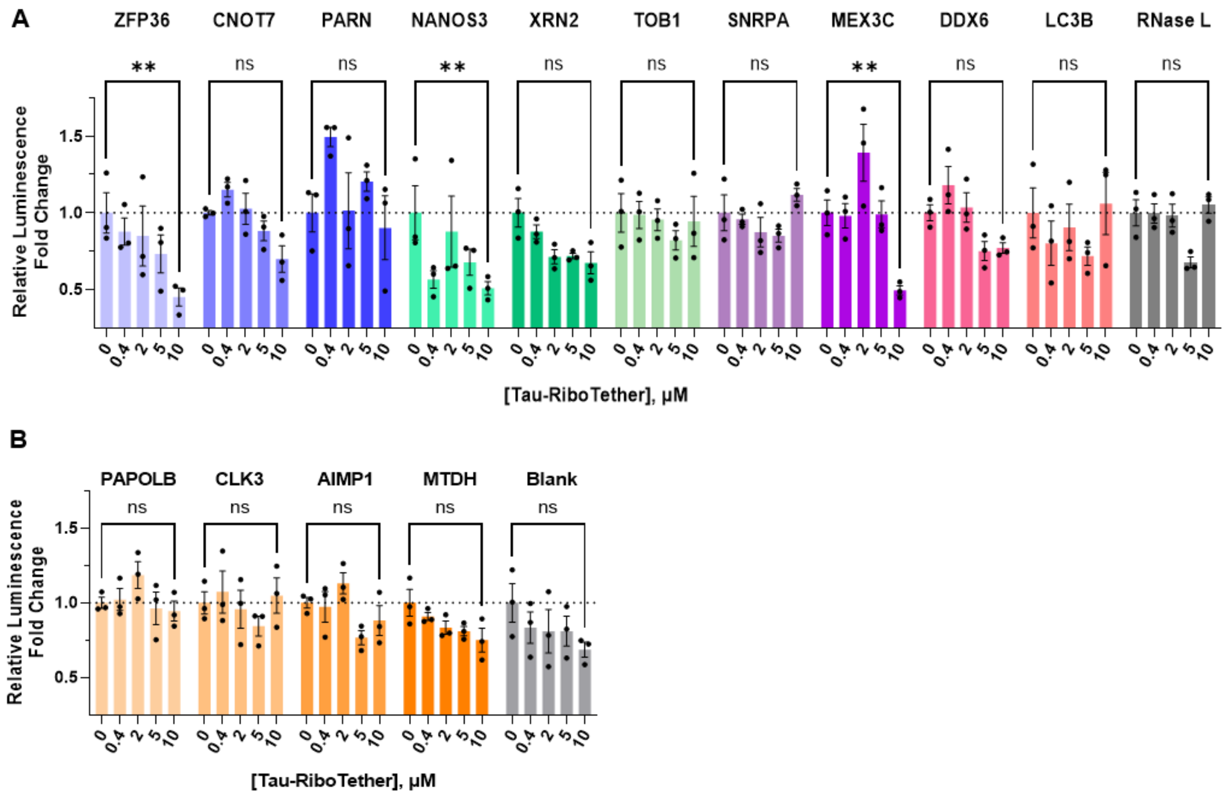


**Figure S7. Dose-response analysis of the FKBP12$^{F36V}$-tagged effectors screen.** Bar graphs correspond to the heatmap shown in **Figure 1D (right)**. Effect of **Tau-RiboTether** on firefly/Renilla fold change following treatment for 48 h, normalized to the corresponding vehicle-treated condition (0.1% (v/v) DMSO). "0" indicates vehicle-treated cells (0.1% (v/v) DMSO). ns, non-significant; $**p < 0.01$, $***p < 0.001$, as determined by two-way ANOVA test. All data are reported as the mean ± SEM (n = 3 biological replicates).

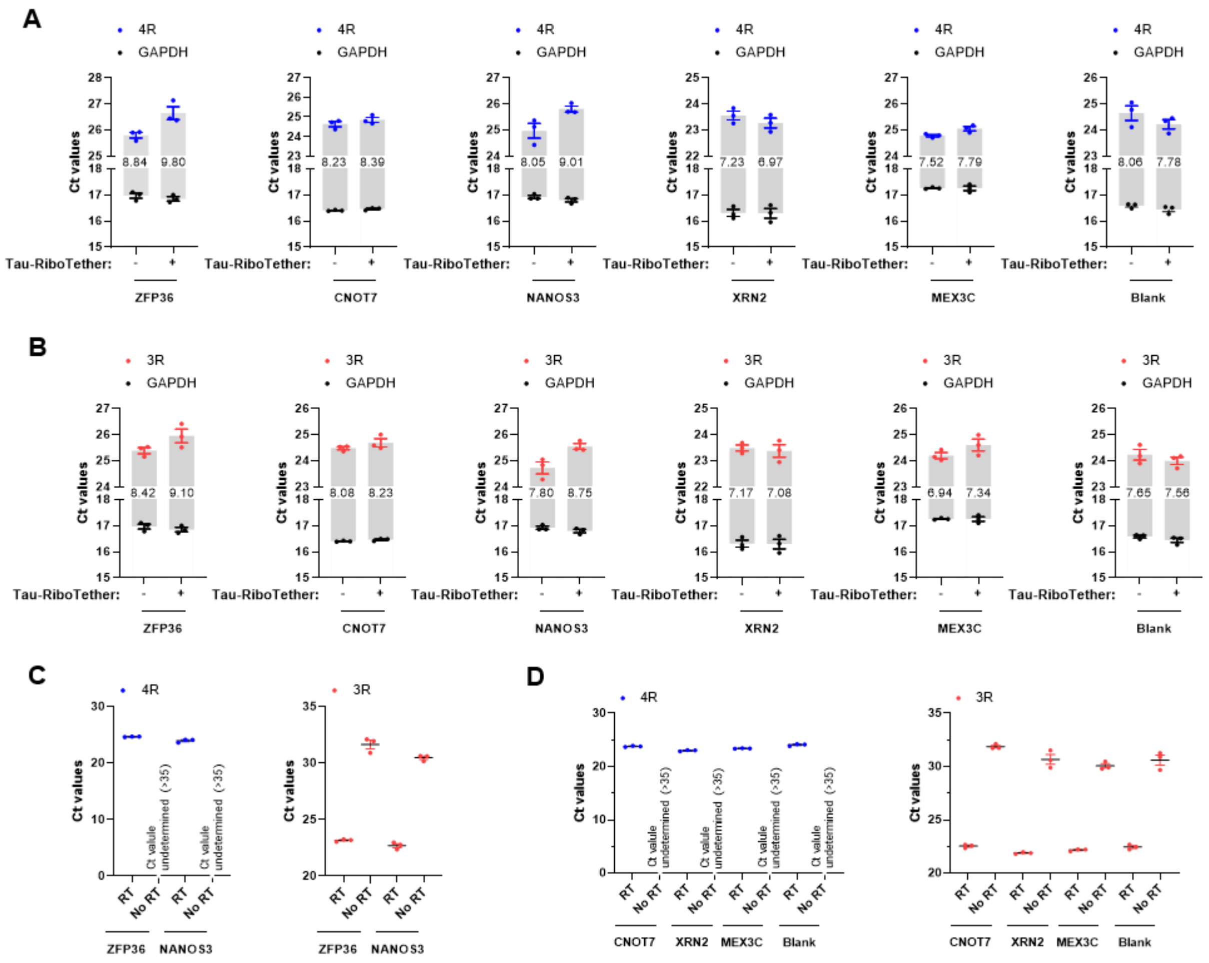


**Figure S8. Raw $C_t$ values for RT-qPCR results presented in Figure 1D. (A)** Raw $C_t$ values for *4R Tau* mRNA levels upon treatment with 10 µM of **Tau-RiboTether** for 48 h. Plots are annotated with $\Delta C_t$ (*4R Tau* – *GAPDH*) values. **(B)** Raw $C_t$ values for *3R Tau* mRNA levels upon treatment with 10 µM of **Tau-RiboTether** for 48 h. Plots are annotated with $\Delta C_t$ (*3R Tau* – *GAPDH*) values. **(C)** $C_t$ values for *4R* and *3R Tau-Luc* in HeLa cells expressing ZFP36 or NANOS3 tagged effectors not subjected to the reverse transcription step (no RT) afforded high (>30) or undetermined (>35) $C_t$ values, indicating sufficient *Tau-Luc* plasmid DNA removal during RNA extraction. **(D)** For the other four tagged effectors, extracted cellular total RNA without reverse transcription (no RT) also showed quite high (>30) or undetermined (>35) $C_t$ values, indicating sufficient *Tau-Luc* plasmid DNA removal during RNA extraction. "-" indicates vehicle-treated cells (0.1% (v/v) DMSO). All data are reported as the mean ± SEM (n = 3 biological replicates).

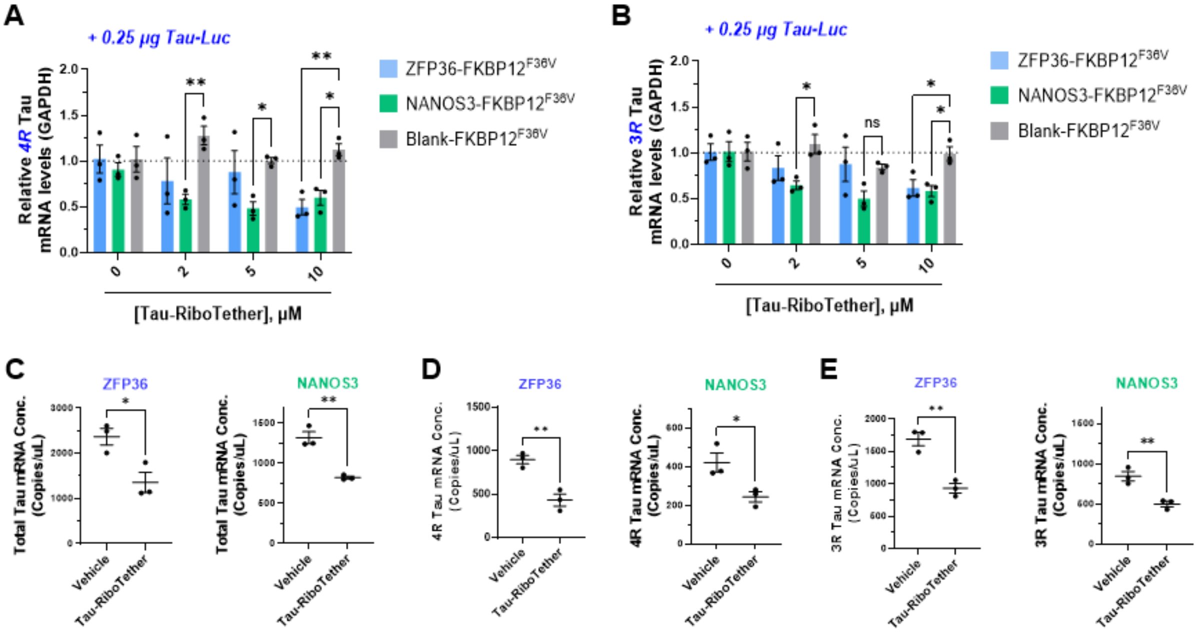


**Figure S9. Tau-RiboTether reduced *Tau-Luc* mRNA levels via recruitment of ZFP36-FKBP12$^{F36V}$ and NANOS3-FKBP12$^{F36V}$ fusion proteins. (A)** Abundance of *4R Tau* mRNA upon **Tau-RiboTether** treatment (0, 2, 5, 10 µM) for 48 h with co-transfection of plasmids encoding different FKBP12$^{F36V}$ effector proteins and 0.25 µg of *Tau-Luc* plasmid. **(B)** Abundance of *3R Tau* mRNA upon treatment of HeLa cells expressing FKBP12$^{F36V}$ effector protein with **Tau-RiboTether** (0, 2, 5, 10 µM) for 48 h. **(C)** Droplet digital (dd)PCR analysis of absolute total *Tau* mRNA concentration upon treatment with 10 µM of **Tau-RiboTether**. **(D)** ddPCR analysis of absolute *4R Tau* mRNA concentration upon treatment with 10 µM **Tau-RiboTether** treatment. **(E)** ddPCR analysis of absolute *3R Tau* mRNA concentration upon treatment with 10 µM **Tau-RiboTether** treatment. Vehicle is 0.1% (v/v) DMSO. ns, non-significant; *$p < 0.05$, **$p < 0.01$, as determined by two-way ANOVA (panels A-B) or two-tailed Student's t-test (panels C-E). All data are reported as the mean ± SEM (n = 3 biological replicates).

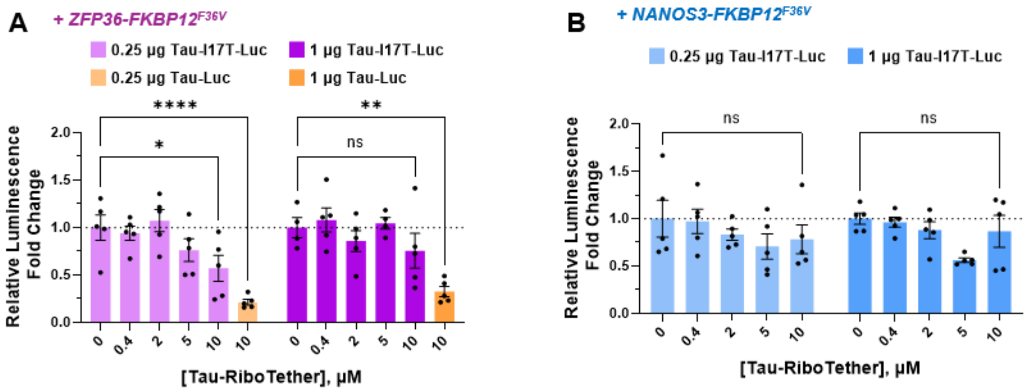


**Figure S10. Effect of Tau-RiboTether on Tau-I17T-Luc (binding site of Tau binder ablated) luminescence. (A)** Effect of **Tau-RiboTether** (10 µM) on luminescence generated from *Tau-I17T-Luc* in HeLa cells expressing ZFP36-FKBP12$^{F36V}$ . **(B)** Effect of **Tau-RiboTether** (10 µM) on luminescence generated from *Tau-I17T-Luc* in HeLa cells expressing NANOS3-FKBP12$^{F36V}$. "0" indicates vehicle-treated cells (0.1% (v/v) DMSO). ns, non-significant; *p < 0.05, ***p < 0.001, ****p < 0.0001, as determined by two-way ANOVA test. All data are reported as the mean ± SEM (n = 5 biological replicates).

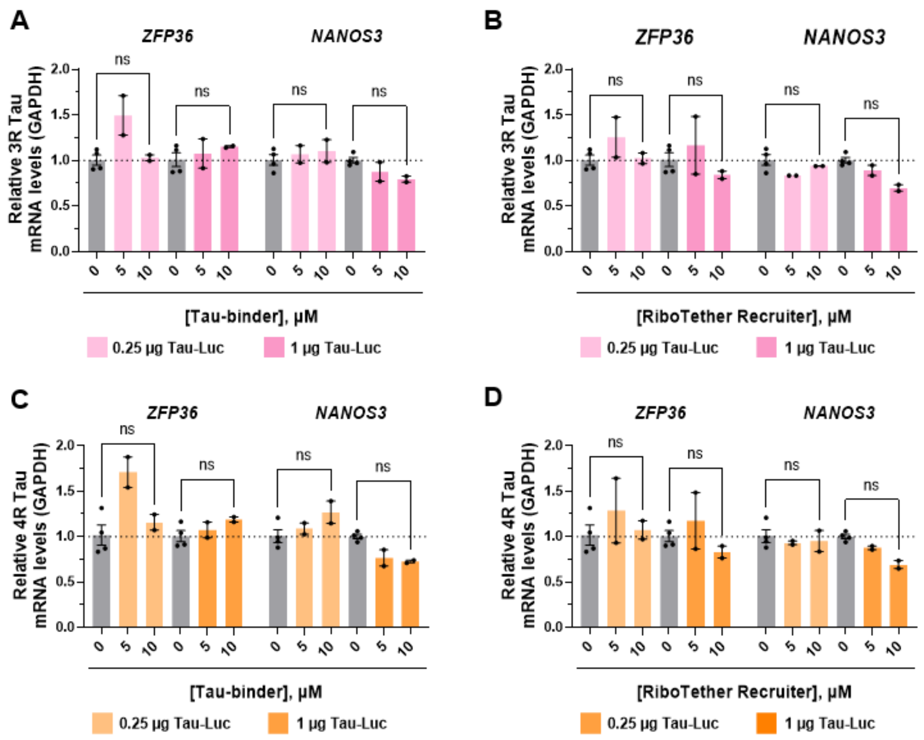


**Figure S11. Effect of Tau-binder and RiboTether Recruiter on *Tau-Luc* transcript abundance in HeLa cells expressing ZFP36-FKBP$^{F36V}$ or NANOS3-FKBP$^{F36V}$. (A)** Abundance of *3R Tau* mRNA upon **Tau-binder** treatment (48 h) in HeLa cells expressing Tau-Luc and ZFP36-FKBP12$^{F36V}$ or NANOS3-FKBP12$^{F36V}$, as determined by RT-qPCR. **(B)** Abundance of *3R Tau* mRNA upon **RiboTether Recruiter** treatment (48 h) in HeLa cells expressing Tau-Luc and ZFP36-FKBP12$^{F36V}$ or NANOS3-FKBP12$^{F36V}$, as determined by RT-qPCR. **(C)** Abundance of *4R Tau* mRNA upon **Tau-binder** treatment (48 h) in HeLa cells expressing Tau-Luc and ZFP36-FKBP12$^{F36V}$ or NANOS3-FKBP12$^{F36V}$, as determined by RT-qPCR. **(B)** Abundance of *4R Tau* mRNA upon **RiboTether Recruiter** treatment (48 h) in HeLa cells expressing Tau-Luc and ZFP36-FKBP12$^{F36V}$ or NANOS3-FKBP12$^{F36V}$, as determined by RT-qPCR. ns, non-significant, as determined by two-way ANOVA test. All data are reported as the mean ± SEM (n = 4 biological replicates for vehicle treated groups (0.1% (v/v) DMSO), n = 2 biological replicates for compound treated groups).

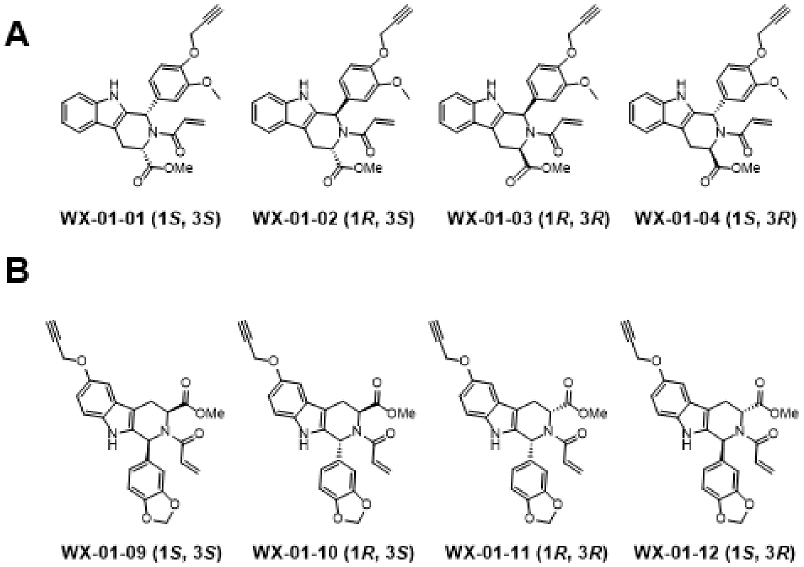


**Figure S12. Structures for stereoisomers of WX-01-02 (A) and WX-01-10 (B)[7].**

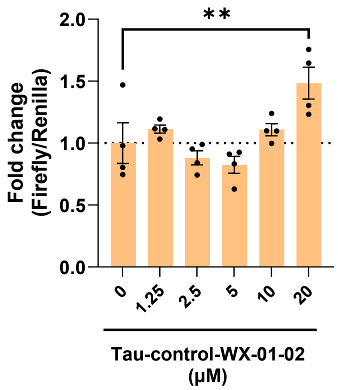


**Figure S13. Concentration-dependent effects of Tau-control-WX-01-02 on Tau-Luc reporter output**. HeLa cells transfected with the *Tau* mini-gene reporter were treated with the indicated compounds for 48 h. Firefly luminescence was normalized to Renilla luminescence and expressed relative to the corresponding vehicle-treated condition (0.2% (v/v) DMSO). Data are reported as the mean ± SEM from n = 4 biological replicates, with individual measurements shown. ns, not significant; **$p < 0.001$, as determined by two-way ANOVA with multiple-comparisons correction.

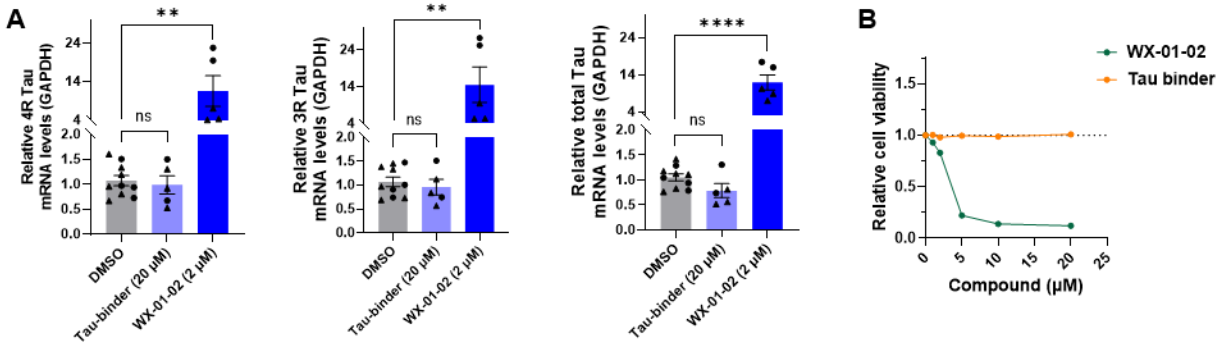


**Figure S14. Effect of Tau-binder and WX-01-02 on *Tau-Luc* mRNA levels and cell viability after 48 h treatment. (A)** Abundance of *4R*, *3R*, and total *Tau* mRNA with compound treatment for 48 h in HeLa cells expressing the *Tau* mini-gene. All data are reported as the mean ± SEM (● and ▲ denote data points collected from two independent experiments; n = 10 biological replicates for vehicle treated groups (0.2% (v/v) DMSO), n = 5 biological replicates for compound treated groups). **(B)** Relative cell viability upon compound treatment for 48 h in HeLa cells expressing the *Tau* mini-gene. **WX-01-02** showed evident cell toxicity at concentrations > 2 µM. ns, non-significant; **p < 0.01, ****p < 0.0001, as determined by two-way ANOVA test. All data are reported as the mean ± SEM (n = 3 biological replicates).

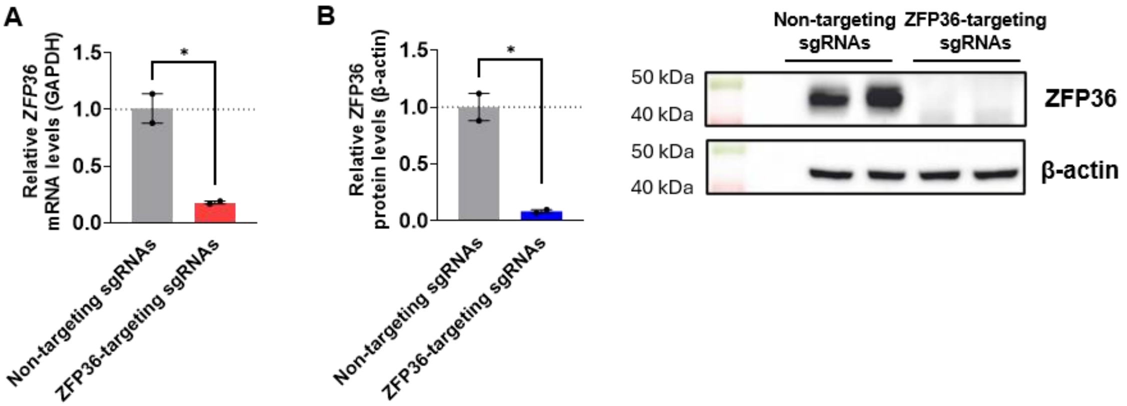


**Figure S15. Validation of ZFP36 CRISPR knockout and analysis of the knock out efficiency in HeLa cells. (A)** Relative abundance of *ZFP36* mRNA upon delivery of non-targeting sgRNAs or *ZFP36*-targeting sgRNAs via electroporation of HeLa cells. **(B)** Relative abundance of ZFP36 protein upon delivery of non-targeting sgRNAs or *ZFP36*-targeting sgRNAs via electroporation of HeLa cells. *$p < 0.05$, as determined by two-way ANOVA test. All data are reported as the mean ± SEM (n = 2 biological replicates).

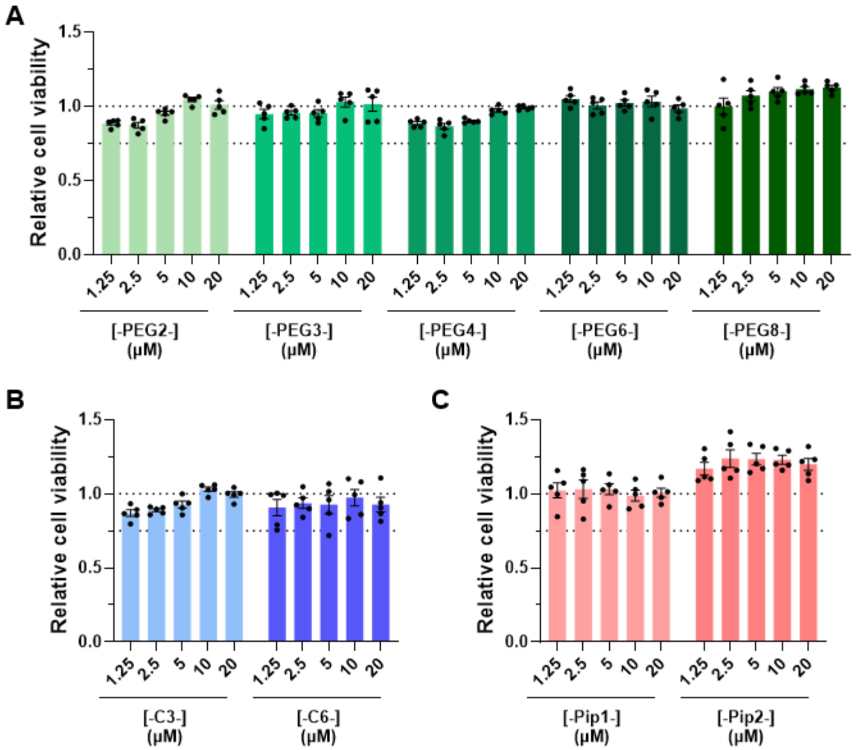


**Figure S16. Relative cell viability upon Tau-[linker]-WX-01-02 compound treatment for 48 h in HeLa cells expressing the *Tau* mini-gene.** All data are reported as the mean ± SEM (n = 5 biological replicates).

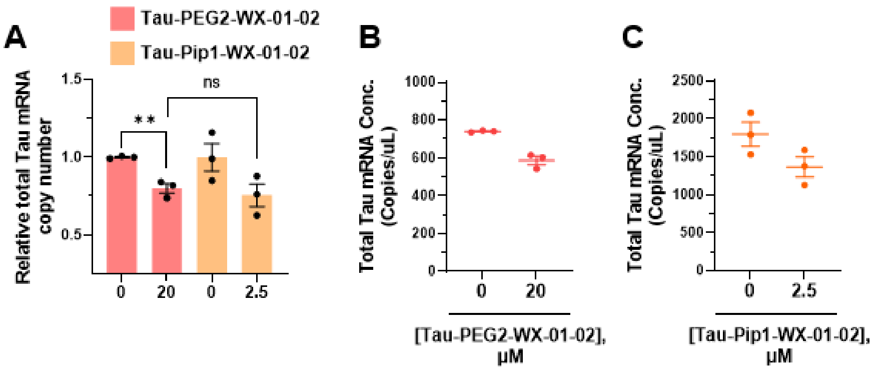


**Figure S17. Relative quantification of total *Tau-Luc* transcript abundance in HeLa cells by ddPCR following treatment with Tau-PEG2-WX-01-02 (20 μM) or Tau-Pip1-WX-01-02 (2.5 μM) for 48 h. (A)** Total *Tau-Luc* transcript copy number was normalized to the corresponding vehicle-treated condition (0.2% (v/v) DMSO). The same amount of input was used for each sample (400 ng of total RNA per RT reaction; 6 ng cDNA per ddPCR reaction). **(B)** Changes of *Tau-Luc* transcript copy number by **Tau-PEG2-WX-01-02 (20 μM)** without normalization to vehicle-treated condition. **(C)** Changes of *Tau-Luc* transcript copy number by **Tau-Pip1-WX-01-02 (2.5 μM)** without normalization to vehicle-treated condition. Data are reported as the mean ± SEM (n = 3 biological replicates), with individual measurements shown. ns, not significant; $^{**}p < 0.01$, as determined by two-tailed Student's t-test.

#### MATERIALS & METHODS

**Plasmid construction.** FKBP12$^{F36V}$ library plasmids encoding FKBP12$^{F36V}$ fused to the C-terminus of the effector protein candidates were constructed by GenScript via restriction enzyme cloning. In brief, a parent pCMV6-Entry plasmid was digested by restriction enzymes AsiSI and MluI to afford the linearized vector. Another DNA fragment containing the sequence encoding a specific effector protein, its C-terminal FKBP12$^{F36V}$ tag, and flanking AsiSI/MluI recognition sequence was PCR amplified, followed by the same restriction digestion to generate overhangs complementary to those of the linearized vector.

FKBP12$^{F36V}$-NLuc plasmid encoding FKBP12$^{F36V}$ fused to the N-terminus of NanoLuciferase (NLuc) were constructed by NEBuilder HiFi assembly. Briefly, the parent pCMV6-Entry vector was linearized by restriction digestion (AsiSI and MluI), followed by assembling FKBP12$^{F36V}$ and NLuc CDS sequence via NEBuilder HiFi DNA Assembly Cloning Kit (NEB, Cat# E5520). The amino acid linker used between the FKBP12$^{F36V}$ and NLuc fragments was AIAGGGGS.

ZFP36-NLuc plasmid encoding ZFP36 fused to the N-terminus of NanoLuciferase (NLuc) were constructed by NEBuilder HiFi assembly. Briefly, the parent pCMV6-Entry vector was linearized by restriction digestion (AsiSI and MluI), followed by assembling ZFP36 and NLuc CDS sequence via NEBuilder HiFi DNA Assembly Cloning Kit (NEB, Cat# E5520). The amino acid linker used between the FKBP12$^{F36V}$ and NLuc fragments was AIAGGGGS. ZFP36-NLuc-C67A_C249A_C250A_C253A mutant plasmid was constructed via Q5 Site-Directed Mutagenesis Kit (NEB, Cat# E0554S).

Full plasmid sequencing files for all plasmids were deposited on Mendeley (doi: 10.17632/pkc7y8d8zy.1).

**Cell culture.** HeLa cells were cultured in DMEM medium (Corning, Cat# 15017CV) supplemented with glutaGRO (Corning, Cat# 25015CI), sodium pyruvate (Corning, Cat# 25000CI) and 10% (v/v) fetal bovine serum (FBS, Gibco, Cat# 26140079) at 37 °C with 5% $CO_2$. All cells were used before passage 20 and tested to be free of mycoplasma contamination (PromoKine, Cat# PK-CA91-1024) prior to cellular experiments.

**Luciferase assay.** HeLa cells were seeded in 6-well plates such that they reached ~80% confluency the next day (~$3\times10^5$ cells/well). Transfection of *Tau-Luc* reporter plasmid, Renilla reporter plasmid, and FKBP12$^{F36V}$-related plasmid were transfected by using JetPrime (Polyplus, Cat# 101000015) according to manufacturer's instructions. In brief, for each well in 6-well plates, Tau-NLuc plasmid (0.25-1 µg), Renilla plasmid (0.2 µg), and a related FKBP$^{F36V}$ plasmid (optimized for different constructs, i.e. 0.4 µg for ZFP36-, CNOT7-, RNase L, and PAPOLB-FKBP$^{F36V}$ to avoid cell toxicity caused by higher expression levels; 1.6 µg for PARN- and CLK3-FKBP$^{F36V}$; and 0.8 µg for other RBP-FKBP$^{F36V}$) were diluted in 200 µL of jetPRIME buffer first, followed by 10 s vortex and then spun down. For mini-gene test without overexpression of FKBP12$^{F36V}$-fused effectors, Tau-NLuc plasmid (0.25 µg), Renilla plasmid (0.2 µg), and a carrier pladmis pcDNA3.1 (0.4 µg) were co-transfected for each well in 6-well plates. For mini-gene test without overexpression of FKBP12$^{F36V}$-fused effectors but with overexpression of ZFP36, Tau-NLuc plasmid (0.25 µg), Renilla plasmid (0.2 µg), and pCMV6-ZFP36 plasmid (0.4-1.6 µg, Origene, Cat# RC202049) were co-transfected for each well in 6-well plates. The jetPRIME transfection reagent was added following 1:2 ratio (i.e., 2 µL transfection reagent per µg of plasmid DNA), followed by 1 s vortex and then spun down. After 10 min incubation at room temperature, the prepared ~200 µL mixture was added into the well after replacing the old cell culture medium with 2 mL new cell culture medium.

The culturing medium containing the transfection cocktail was removed 5 h post-transfection, and the cells were trypsinized followed by seeding into 96-well plates to reach ~40% confluency after 16 h (i.e., resuspend cells trypsinized from each well from the 6-well plates into 12 mL cell culture medium, then add 100 µL this cell suspension into each well in 96-well plates). Compound or DMSO (0.1% (v/v)) was added to each well by first diluting 1000x compound stock solution or DMSO into cell culture medium, and then replacing the 100 µL old cell culture medium with 100 µL new compound/DMSO-containing cell culture medium in each well, followed by incubated for another 48 h. After the treatment period, cell viability in each well was first measured by using CellTiter-Fluor Kit (Promega, Cat# G6081) according to manufacturer's instructions, followed by measuring luciferase activities by using the Dual-Glo Luciferase assay Kit (Promega, Cat# E2940) according to manufacturer's instructions.

**Measurement of *Tau-Luc* mRNA abundance by RT-qPCR.** HeLa cells were seeded, transfected, and treated with DMSO or compound as described in **Luciferase assay**. After 48 h treatment, total cellular RNA was extracted by using Quick-RNA Miniprep Kit (Zymo, Cat# R1054) per the manufacturer's instructions including the on-column DNase I digestion. Reverse transcription (RT) was performed for each biological replicate using qScript cDNA synthesis kit (QuantaBio), with an input of 400 ng total RNA in a total volume of 20 µL. Quantitative PCR (qPCR) was completed using 2 µL of cDNA, 300 nM of forward and reverse gene-specific primers (**Table S2**), and Power SYBR Green Master Mix (Applied Biosystems) in a total volume of 33 µL. Next, three aliquots of 10 µL was aliquoted into 384-well plates, affording three technical replicates per biological sample. A QuantStudio 5 Real-Time PCR system was used to measure $C_t$ values and melting curves. Relative expression level of a specific transcript was calculated by the ΔΔCt method[8] by comparison to *GAPDH*.

**Validation of fusion protein expression by Western blotting.** HeLa cells were seeded into 6-well plates such that the reached ~50% confluency after overnight incubation. The next day, FKBP-related plasmid was transfected by using JetPrime (Polyplus, Cat# 101000015) according to manufacturer's instructions. The plasmid transfection amount was optimized for different constructs, i.e. 0.2 µg for ZFP36-, CNOT7-, RNase L, and PAPOLB-FKBP$^{F36V}$ to avoid cell toxicity caused by higher expression levels; 0.8 µg for PARN- and CLK3-FKBP$^{F36V}$; and 0.4 µg for other RBP-FKBP$^{F36V}$. The cell culturing medium containing the transfection cocktail was replaced 5 h post-transfection, and the cells were incubated for another 48 h.

After the incubation period, the cells were washed twice with 1× DPBS, scraped into 1 mL of 1× DPBS, and pelleted by centrifµged in 1.6 mL tubes for 2 min at 4 °C and 2500 rpm. Cell pellets were then resuspended in 50 µL of Mammalian Protein Extraction Reagent (M-PER, Thermo Scientific) to which 1x Mammalian Protease Inhibitor Cocktail (RPI, Cat# P50700-1, 100x stock) was added, and the cells were lysed by placing the resuspsended pellets on ice for 15 min. After centrifµgation at at 4 °C and 14000 × g for 15 min, total cellular protein was harvested in the supernatant and quantified by using Micro BCA Protein Assay Kit (Thermo Scientific, Cat# 23235). Approximately 20 µg total protein per biological replicate was separated by sodium dodecyl sulfate-polyacrylamide gel electrophoresis (SDS-PAGE; 10% (w/v)) using 1× Running buffer (25 mM Tris, pH 8.3, 190 mM glycine, and 0.1% (w/v) SDS). Proteins were then electroblotted to a polyvinylidene fluoride (PVDF) membrane (0.45 µm, Cytiva) in 1× Transfer buffer (25 mM Tris, pH 8.3, 190 mM glycine, and 20% (v/v) MeOH).

For immunoblotting, the membrane was washed first with 1× TBST (20 mM Tris-HCl, pH 7.6, 150 mM NaCl, and 0.1% (v/v) Tween-20), followed by blocking with 5% (w/v) non-fat dry milk in 1× TBST for 1 h at room temperature. Anti-FLAG primary antibody (Cell Signaling Technology, Cat# 2368 in Figure S2; Cell Signaling Technology, Cat# 14793 in Figure S1) was diluted 1000-fold in 1× TBST containing 5% (w/v) non-fat dry milk and incubated with the blocked membrane for 16 h at 4 °C. After washing three times in 1× TBST, the membrane was incubated with anti-rabbit HRP-linked secondary antibody (Cell Signaling Technology, Cat# 7074, 1:2000 dilution in 1× TBST containing 5% (w/v) non-fat dry milk) for 2 h at room temperature. After washing three times in 1× TBST, imaging of the membrane was performed by incubation with SuperSignal West Pico Chemiluminescent Substrate (Pierce) per the manufacturer's procedure.

**Gel-based ABPP validation of ZFP36 engagement for covalent ligands.** HeLa cells were seeded in T175 flasks to reach ~70% confluency the next day (~$4.8 \times 10^6$ cells/flask). The next day, the cells were transfected with ZFP36-NLuc plasmid (28.8 µg per flask) using JetPrime (57.6 µL, the transfection mixture was prepared in a total volume of 1 mL) for 6 h after refreshing the cell culturing medium. Following transfection, the cells were trypsinized and re-seeded into two T175 flasks and incubated for another 42 h. At the end point of transfection, **WX-01-10** (20 µM; 0.2% (v/v)), **WX-01-02** (20 µM; 0.2% (v/v) DMSO), or DMSO (0.2% (v/v)) were added by first diluting each compound or DMSO stock into new cell culturing medium and then replacing the old cell culturing medium with the new one containing compounds or DMSO, followed by incubation for 3 h in an incubator.

After compound incubation, the cells were scraped into ice-cold PBS in a 1.5 mL tube, and pelleted at 2000 rpm for 2 min at 4 $^{o}$C. The pellets were washed twice with ice-cold PBS. For each biological replicates, the washed cell pellet was resuspended in 500 µL of NP-40 lysis buffer (25 mM Tris-HCl, pH 7.4, 150 mM NaCl, 10% (v/v) glycerol, and 1% (v/v) NP-40) supplemented with 1x Mammalian Protease Inhibitor Cocktail (RPI, Cat# P50700-1), followed by sonication on ice to lyse the cells (on 1 s, off 1 s, total 8 s, at 10% output, 2 rounds for each sample). Each sample was then rotated at 4 $^{o}$C for 10 min, followed by centrifµgation at 16000 × g for 5 min at 4 $^{o}$C to remove cellular debris. Total cellular proteins were harvested in the supernatant and quantified using Micro BCA Protein Assay Kit (Thermo Scientific, Cat# 23235), affording concentrations of 1-2 mg/mL.

Pull-down of FLAG-tagged ZFP36-NLuc protein was performed by using Pierce™ anti-DYKDDDDK magnetic agarose beads (Thermo Scientific, Cat# 36797). In brief, 20-25 µL of magnetic beads were placed into a 1.5 mL microcentrifµge tube, followed by addition of 200 µL of ice-cold IP wash buffer (25 mM Tris-HCl, pH 7.4, 150 mM NaCl, and 0.2% (v/v) NP-40). The tube was gently vortexed to mix and placed onto a magnetic stand to collect the beads. The supernatant was removed, followed by 2 × 1 mL washes with the same IP wash buffer. Finally, the cell lysate corresponding to 1.5 mg of total cellular proteins was added to the beads (pre-adjusting cell lysate volume to reach 500 µL for each sample). Samples were then rotated for 2 h at 4 $^{o}$C, followed by washing the beads to remove non-specific binders using the same ice-cold IP washing buffer four times (1 mL per wash).

After the FLAG pull-down procedure, the beads in each tube were resuspended in 30 µL PBS followed by addition of a "click" mixture (3.6 µL). The click mixture was prepared by sequentially adding 45 µL TBTA solution (1.7 mM, 0.9 mg TBTA/mL in 4:1 *t*-BuOH/DMSO solution), 15 µL $CuSO_4$ solution (12.5 mg/mL in $H_2O$), 15 µL TCEP solution (13 mg/mL in PBS, fresh), vortexing, and finally adding 15 µL TAMRA-$N_3$ to give a red clear solution after mixing. Click reactions were performed at room temperature for 1 h on a shaking bed at 800 rpm protected from light. Next, 4× Laemmli buffer was added to the beads to afford a final concentration of 2×. After the samples were boiled for 10 min, they were cooled down on the benchtop, placed onto a magnetic stand, and the supernatant was collected and placed in a new tube.

SDS-PAGE analysis of samples both before and after pull-down/labeling was performed following the same gel-running procedure described in **Validation of fusion protein expression by Western blotting**, except that before membrane transfer, TAMRA labeling events were visualized on gel instead (imaged by Azure Sapphire Biomolecular Imager).

**ZFP36 NanoBRET assay in live cells.** HeLa cells were seeded in 6-well plates such that they reached ~80% confluency the next day (~$3\times10^5$ cells/well). Transfection was then performed by using JetPrime (Polyplus, Cat# 101000015) according to manufacturer's instructions. In brief, for each well in 6-well plates, 400 ng ZFP36-NLuc plasmid was diluted in 200 µL of jetPRIME buffer first, followed by 10 s vortex and then spun down. The jetPRIME transfection reagent was added following 1:2 ratio (i.e., 0.8 µL transfection reagent per µg of plasmid DNA), followed by 1 s vortex and then spun down. After 10 min incubation at room temperature, the prepared ~200 µL mixture was added into the well after replacing the old cell culture medium with 2 mL new cell culture medium.

The transfected cells were detached with trypsin after 4~6 h transfection, resuspended in Opti-MEM I Reduced Serum Medium without phenol red (Gibco, Cat# 11058021), and reseeded into white 96-well, clear bottom plates (Corning, Cat# 29444-010) to afford ~70% confluency the next day (i.e., resuspend cells trypsinized from each well from the 6-well plates into 7 mL cell culture medium, then add 100 µL this cell suspension into each well in 96-well plates). Meanwhile, 20× stock solutions of **WX-01-02 Tracer** or **WX-01-10 Tracer** were freshly prepared in probe dilution buffer (12.5 mM HEPES, pH 7.5, and 31.25% (w/v) PEG-400) by diluting 1 mM tracer stock (prepared in 100% DMSO) according to a previously described procedure[9]. A 5 µL aliquot of the tracer (0.2% (v/v) DMSO final concentration) was added to each well, followed by 2 h incubation at 37 $^{o}$C. Intracellular TE Nano-Glo Substrate/Inhibitor was then added at 1× final concentration per manufacturer's instructions (Promega, Cat# N2160). Filtered luminescence was measured by the GloMaxDiscover luminometer equipped with 450 nm donor BP filter and 600 nm acceptor LP filter with 0.6 s integration time. Milli-BRET units (mBu) were calculated by multiplying the raw BRET signal values by a factor of 1000 [9]. Apparent probe affinity value ($EC_{50}$) was calculated by plotting ΔmBu (after subtracting background mBu, i.e., cells treated with 0.2% (v/v) DMSO only) as a function of compound concentration and fitting the resulting curve to a sigmoidal dose-response (stimulation) equation available in GraphPad Prism:

$$Y = Bottom + \frac{(Top - Bottom) \cdot X}{EC_{50} + X}$$

where Y is ΔmBu, Bottom is the background mBu level, Top is the saturated ΔmBu level, X is tracer concentration, and the $EC_{50}$ is the apparent probe affinity value.

**$FKBP12^{F36V}$ NanoBRET assay in live cells.** HeLa cells were seeded in 6-well plates such that they reached ~80% confluency the next day (~$3x10^5$ cells/well). Transfection was then performed by using JetPrime (Polyplus, Cat# 101000015) according to manufacturer's instructions. In brief, for each well in 6-well plates, 800 ng $FKBP12^{F36V}$-NLuc plasmid was diluted in 200 µL of jetPRIME buffer first, followed by 10 s vortex and then spun down. The jetPRIME transfection reagent was added following 1:2 ratio (i.e., 1.6 µL transfection reagent per µg of plasmid DNA), followed by 1 s vortex and then spun down. After 10 min incubation at room temperature, the prepared ~200 µL mixture was added into the well after replacing the old cell culture medium with 2 mL new cell culture medium.

The transfected cells were detached with trypsin after 4~6 h transfection, resuspended in Opti-MEM I Reduced Serum Medium without phenol red (Gibco, Cat# 11058021), and reseeded into white 96-well, clear bottom plates (Corning, Cat# 29444-010) to afford ~70% confluency the next day (i.e., resuspend cells trypsinized from each well from the 6-well plates into 7 mL cell culture medium, then add 100 µL this cell suspension into each well in 96-well plates). Meanwhile, 20× stock solutions of **o-AP1867 Tracer** were freshly prepared in probe dilution buffer (12.5 mM HEPES, pH 7.5, and 31.25% (w/v) PEG-400) by diluting 1 mM tracer stock (prepared in 100% DMSO) according to a previously described procedure. A 5 µL aliquot of the tracer (0.2% (v/v) DMSO final concentration) was added to each well, followed by 2 h incubation at 37 °C. When competing **o-AP1867 Tracer** in live cells, the competitor compound was serially diluted in Opti-MEM I Reduced Serum Medium without phenol red to afford their 10× stock solution (10 µL per well, final 0.4% DMSO v/v) and then co-incubated with 8 nM **o-AP1867 Tracer** for 2 h at 37 °C. Intracellular TE Nano-Glo Substrate/Inhibitor was then added at 1× final concentration per manufacturer's instructions (Promega, Cat# N2160). Filtered luminescence was measured by the GloMaxDiscover luminometer equipped with 450 nm donor BP filter and 600 nm acceptor LP filter with 0.5 s integration time. Milli-BRET units (mBu) were calculated by multiplying the raw BRET signal values by a factor of 1000 *9*. Apparent probe affinity value ($EC_{50}$) was calculated by plotting ΔmBu (after subtracting background mBu, i.e., cells treated with 0.2% (v/v) DMSO only) as a function of compound concentration and fitting the resulting curve to a sigmoidal dose-response (stimulation) equation available in GraphPad Prism:

$$Y = Bottom + \frac{(Top - Bottom) \cdot X}{EC_{50} + X}$$

where Y is ΔmBu, Bottom is the background mBu level, Top is the saturated ΔmBu level, X is tracer concentration, and the EC*50* is the apparent probe affinity value.

The $IC_{50}$ of a competing molecule was by using the sigmoidal dose-response (inhibition) equation available in GraphPad Prism:

$$Y = Bottom + \frac{Top - Bottom}{\frac{X}{IC_{50}} + 1}$$

where Y is ΔmBu, Bottom is the background mBu level, Top is the saturated ΔmBu level, X is competitor compound concentration, and the $IC_{50}$ is the half-maximal inhibitory concentration value.

**Construction of control and ZFP36 CRISPR knockout cells.** Delivery of sgRNA and Cas9 ribonucleoprotein (RNP) complex was performed by using SG Cell Line 4D-Nucleofector X Kit L (Lonza, Cat# V4XC-3024) with program designed for HeLa cell line. Generally, two ZFP36-targeting or non-targeting sgRNAs were fully resuspended to afford 100 µM stock solutions. Next, 1.1 µL of one sgRNA, 1.1 µL of the other sgRNA, and 3 µL of Alt-R S.p were gently mixed following by addition of HiFi Cas9 Nuclease V3 protein solution (10 µg/µL, IDT, Cat# 421953095). The samples were then incubated for 20 min at room temperature. HeLa cells were trypsinized, resuspended, and washed twice with PBS. For each electroporation sample, 0.6 million cells were resuspended in 100 µL nucleofector solution (18 µL supplement to 82 µL Nucleofection according to the kit manual) and immediately added into the prepared 5.2 µL RNP complex solution. After mixing by gently pipetting up and down, 100 µL was transferred to the electroporation module. After gently tapping to remove any air bubbles, the celles were electroporated and 500 µL of pre-warmed culture (37 $^{o}$C) medium was added immediately to the cuvette. The electroporated cells were gently transferred into 6-well plates containing 2 mL of pre-warmed medium. Repeated aspiration of the sample was avoided. The cells were grown for another 1 week following stand cell culture procedure; knockout efficiency was validated by qPCR and Western blots as described above.

**Proteomics data reanalysis**

Raw data files were downloaded from the PRIDE repository under the dataset identifier PXD042541. Database searching was performed using Proteome Discoverer 3.3 with the Sequest HT search engine. Cysteine residues were searched with dynamic modifications corresponding to carbamidomethylation (+57.021 Da) or IA-DTB labeling (+455.274 Da). Protein N-termini and lysine residues were searched with a static TMT modification (+229.163 Da). MS3-based quantification was performed using the default settings.

**Synthesis and Characterization of compounds**

**Abbreviations:**

Boc: *tert*-Butoxycarbonyl
BRET: Bioluminescence resonance energy transfer
*t*-BuOH: Tertiary butanol
DAD: Diode array detector
DCM: Dichloromethane
DIPEA: *N*,*N*-Diisopropylethylamine
DMSO: Dimethyl sulfoxide
DMF: *N*,*N*-Dimethylformamide
ESI: Electrospray ionization
$Et_3N$: Triethylamine
FA: Formic acid
HPLC: High-performance liquid chromatography
LCMS: Liquid chromatography mass spectrometry
MeOH: Methanol
NHS: *N*-Hydroxysuccinimide
NMR: Nuclear magnetic resonance
OBD: Optimum bed density
PEG: Polyethylene glycol
PyBOP: Benzotriazol-1-yloxytripyrrolidinophosphonium hexafluorophosphate
TFA: Trifluoroacetic acid
THPTA: Tris(3-hydroxypropyltriazolylmethyl)amine
TLC: Thin-layer chromatography

**General Synthetic Methods**. Chemicals were purchased from Sigma Aldrich, Ambeed, TCI, and Tocris Bioscience. A 400 MHz UltraShieldTM and a 600 MHz UltraShieldTM NMR spectrometer (Bruker) were used to collect $^{1}H$ NMR and $^{13}C$ NMR spectra, respectively. Chemical shifts were reported in ppm and coupling constants were reported in Hz.

Thin layer chromatography (TLC) was performed on Merck $60F_{254}$ precoated silica gel plates, and small molecules or intermediates thereof were visualized by fluorescence quenching under UV light and by staining with phosphomolybdic acid. Reaction progress was also monitored by LC/MS on Agilent 1260 Infinity II system equipped with1260 Infinity II binary pump, 1260 Infinity II DAD, 1260 Infinity II Multisampler, and 6130 Single Quadruple LCMS (ESI). An Agilent SB-C18 column (1.8 μm OBD, 2.1 × 50 mm) was employed with a linear gradient of MeOH (Solvent B) containing 0.1% (v/v) formic acid (FA) in water containing 0.1% (v/v) FA (Solvent A) at a flow rate of 0.350 mL/min. A mode of linear gradient of B in A + B, 0 to 95%, was performed over 9 min.

Preparative HPLC was performed with a Binary HPLC pump (Waters 1525), a dual absorbance detection system (Waters 2487), and a Sunfire C18 OBD S-14 column (Waters, 5 µm, 19 ×150 mm). Purity of final products was assessed by analytical HPLC using a Symmetry C18 column (Waters, 5 µm, 4.6 × 150 mm) with flow rate set as 1 mL/min and a linear gradient (0-100% MeOH in water added with 0.1% (v/v) TFA over 60 min); absorbances were monitored at 220 and 254 nm.

High-resolution mass spectra of compounds were acquired by using positive and negative ESI mode on an Orbitrap Exploris 120 (Thermo Fisher Scientific) that was coupled to a Vanquish HPLC system (Thermo Fisher Scientific).

**WX-01-02** and **WX-01-10** were synthesized according to previously reported procedures.[7, 10]

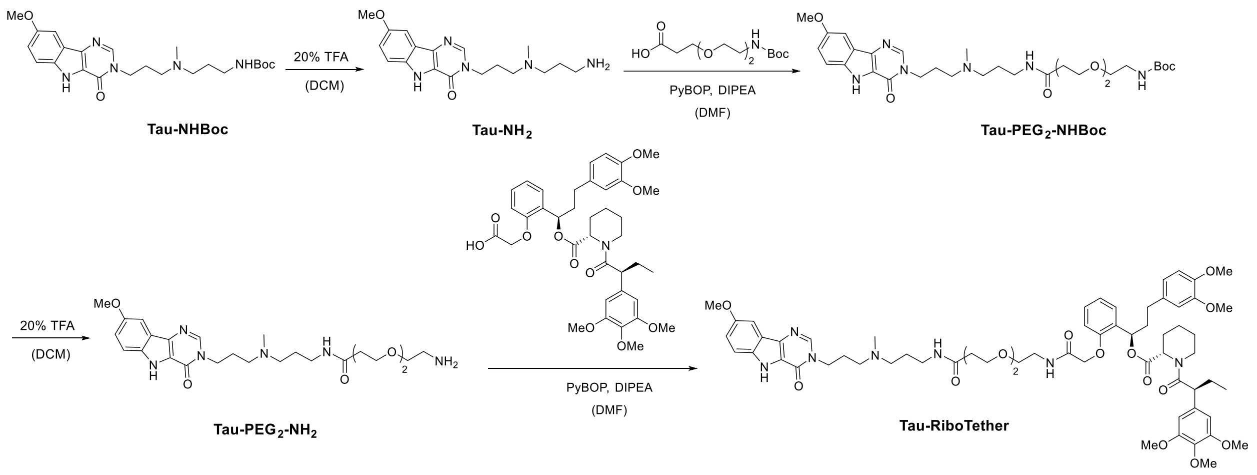


**Scheme 1. Synthesis of Tau-RiboTether.**

**Synthesis of Tau-RiboTether.** **Tau-NHBoc** was synthesized according to previously reported procedure.[11] **Tau-NHBoc** (23 mg) was dissolved in 2 mL of DCM:TFA (4:1) solvent, incubated for 3 h with stirring at room temperature, and then dried overnight to afford **Tau-NH$_2$** (97% yield, 17.3 mg), which was used without further purification. NHBoc-PEG$_2$-Acid (21.0 mg, 75.6 µmol) was dissolved in 1 mL of DMF followed by the addition of PyBOP (78.6 mg, 151.1 µmol) and DIPEA (52.6 µL, 302.2 µmol). The reaction was stirred at room temperature for 10 min followed by the addition of **Tau-NH$_2$** (17.3 mg, 50.4 µmol). The reaction was stirred at room temperature for 16 h and then diluted by adding 1 mL of MeOH containing 0.1% (v/v) TFA. The product **Tau-PEG$_2$-NHBoc** (49% yield, 14.9 mg) was purified by HPLC as described in the **General Synthetic Methods**, with a linear gradient from 0-100% MeOH in water supplemented with 0.1% (v/v) TFA over 60 min.

**Tau-PEG$_2$-NHBoc** (14.9 mg) obtained from HPLC purification was dissolved in 1.5 mL of DCM:TFA (4:1) solvent, incubated for 3 h with stirring at room temperature, and then dried overnight to afford **Tau-PEG$_2$-NH$_2$** (97% yield, 12.0 mg), which was used without further purification. **Ortho-AP1867** (5.5 mg, 7.9 µmol) was dissolved in 1 mL of DMF followed by the addition of PyBOP (12.4 mg, 23.8 µmol) and DIPEA (8.3 µL, 47.6 µmol). The reaction was stirred at room temperature for 10 min followed by the addition of **Tau-PEG$_2$-NH$_2$** (6.0 mg, 11.9 µmol). The reaction was stirred at room temperature for 16 h then diluted by adding 1 mL of MeOH containing 0.1% (v/v) TFA. The product **Tau-RiboTether** (62% yield, 5.8 mg) was purified by HPLC as described in the **General Synthetic Methods**, with a linear gradient from 0-100% methanol in water supplemented with 0.1% (v/v) TFA over 60 min. $^1$H NMR (400 MHz, Methanol-*d*$_4$) δ 8.28 (dd, *J* = 6.9, 2.9 Hz, 1H), 7.58 – 7.45 (m, 2H), 7.26 – 7.13 (m, 2H), 6.91 – 6.70 (m, 5H), 6.63 (d, *J* = 6.0 Hz, 2H), 6.09 (dd, *J* = 8.0, 5.8 Hz, 1H), 5.47 – 5.36 (m, 1H), 4.55 (d, *J* = 15.2 Hz, 1H), 4.43 (dd, *J* = 14.6, 3.8 Hz, 1H), 4.27 (dt, *J* = 11.5, 6.6 Hz, 2H), 4.13 (d, *J* = 13.3 Hz, 1H), 3.87 (s, 3H), 3.78 (d, *J* = 5.4 Hz, 6H), 3.68 (d, *J* = 7.0 Hz, 9H), 3.60 (ddd, *J* = 16.0, 6.2, 3.8 Hz, 2H), 3.50 – 3.38 (m, 9H), 3.24 – 3.06 (m, 4H), 2.89 (d, *J* = 6.9 Hz, 3H), 2.69 – 2.21 (m, 10H), 2.08 – 1.89 (m, 5H), 1.78 – 1.48 (m, 5H), 1.29 (s, 1H), 1.22 – 1.11 (m, 1H), 0.88 (t, *J* = 7.3 Hz, 3H). $^{13}$C NMR (151 MHz, MeOD) δ 173.97, 173.47, 171.11, 169.48, 155.02, 154.09, 153.21, 148.92, 147.38, 143.17, 136.60, 135.56, 134.80, 133.68, 129.02, 128.71, 126.77, 122.06, 121.84, 121.05, 120.42, 119.07, 113.42, 112.23, 111.73, 105.20, 100.20, 69.82, 69.80, 69.61, 68.82, 67.02, 66.67, 59.68, 55.17, 55.14, 55.08, 54.75, 53.76, 53.57, 52.11, 49.57, 48.17, 43.63, 43.07, 38.61, 38.46, 36.27, 36.01, 35.05, 30.74, 27.92, 26.19, 24.97, 24.55, 24.47, 20.51, 11.18. HRMS (ESI) m/z: $[M + 2H]^{2+}$ for $C_{63}H_{85}N_7O_{15}^{2+}$, calculated 589.8046, found 589.8046.

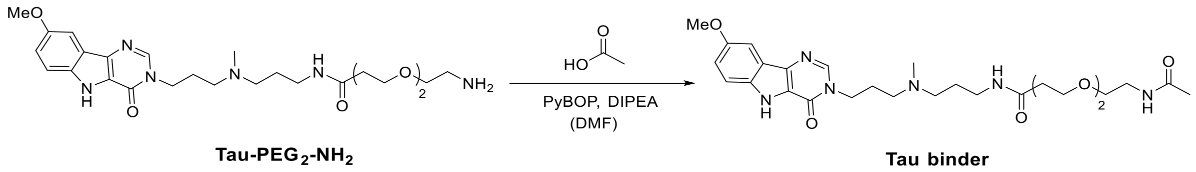


**Scheme 2. Synthesis of Tau-binder.**

**Synthesis of Tau-binder.** Acetic acid (0.5 µL, 9.0 µmol) was dissolved in 0.5 mL of DMF followed by the addition of PyBOP (9.3 mg, 17.9 µmol) and DIPEA (6.2 µL, 35.8 µmol). The reaction was stirred at room temperature for 10 min followed by the addition of **Tau-PEG$_2$-NH$_2$** (3 mg, 6.0 µmol). The reaction was stirred at room temperature for 16 h then diluted by adding 1 mL of MeOH containing 0.1% (v/v) TFA. The product **Tau-binder** (71% yield, 2.3 mg) was purified by HPLC as described in the **General Synthetic Methods**, with a linear gradient from 0-100% MeOH in water supplemented with 0.1% (v/v) TFA over 60 min. $^1$H NMR (400 MHz, Methanol-*d*$_4$) δ 8.30 (s, 1H), 7.55 (d, *J* = 2.3 Hz, 1H),

7.49 (d, $J$ = 9.0 Hz, 1H), 7.18 (dd, $J$ = 9.0, 2.5 Hz, 1H), 4.31 (t, $J$ = 6.9 Hz, 2H), 3.90 (s, 3H), 3.71 – 3.59 (m, 3H), 3.50 – 3.44 (m, 7H), 3.26 – 3.22 (m, 2H), 3.16 – 3.07 (m, 2H), 2.91 (s, 3H), 2.42 (q, $J$ = 6.1, 5.7 Hz, 2H), 2.32 (p, $J$ = 7.2 Hz, 2H), 2.02 – 1.89 (m, 5H), 1.29 (s, 2H). HRMS (ESI) m/z: $[M - H]^-$ for $C_{63}H_{83}N_7O_{15}$, calculated 543.2937, found 543.2926.

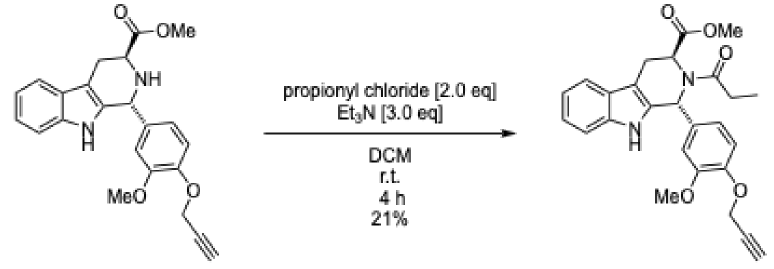


**Scheme 3. Synthesis of control WX-01-02.**

**Synthesis of Methyl (1*R*,3*S*)-1-(3-methoxy-4-(prop-2-yn-1-yloxy)phenyl)-2-propionyl-2,3,4,9-tetrahydro-1*H*-pyrido[3,4-*b*]indole-3-carboxylate (control WX-01-02)**: To a solution of methyl (1*R*,3*S*)-1-(3-methoxy-4-(prop-2-yn-1-yloxy)phenyl)-2,3,4,9-tetrahydro-1*H*-pyrido[3,4-*b*]indole-3-carboxylate[6] (WX-01-02, 9.70 mg, 0.0248 mmol) in DCM (0.20 mL) was added $Et_3N$ (10.7 mL, 0.0768 mmol) and propionyl chloride (4.45 mL, 0.0512 mmol) at room temperature. The reaction mixture was stirred at the same temperature for 4 h and then concentrated under reduced pressure. The resulting residue was purified by HPLC to obtain **control WX-01-02** (2.30 mg, 0.00515 mmol, 21%) as a white solid. $^1H$ NMR (600 MHz, $CDCl_3$): d 7.78-7.69 (br, 1H), 7.53 (d, $J$ = 7.8 Hz, 1H), 7.20-7.14 (m, 1H), 7.12 (t, $J$ = 7.3 Hz, 1H), 7.05-6.93 (brs, 1H), 6.93-6.83 (br, 2H), 6.20-5.90 (brs, 1H), 5.30-5.00 (br, 1H), 4.85-4.60 (br, 2H), 3.83 (s, 3H), 3.64 (s, 3H), 3.59-3.47 (br, 1H), 3.46-3.13 (br, 1H), 2.56-2.40 (m, 2H), 2.36-2.14 (br, 1H), 1.15-1.00 (br, 3H); HRMS (ESI), *m/z* calcd for $C_{26}H_{27}N_2O_5$ $[M+H]^+$ 447.1914, found 447.1912.

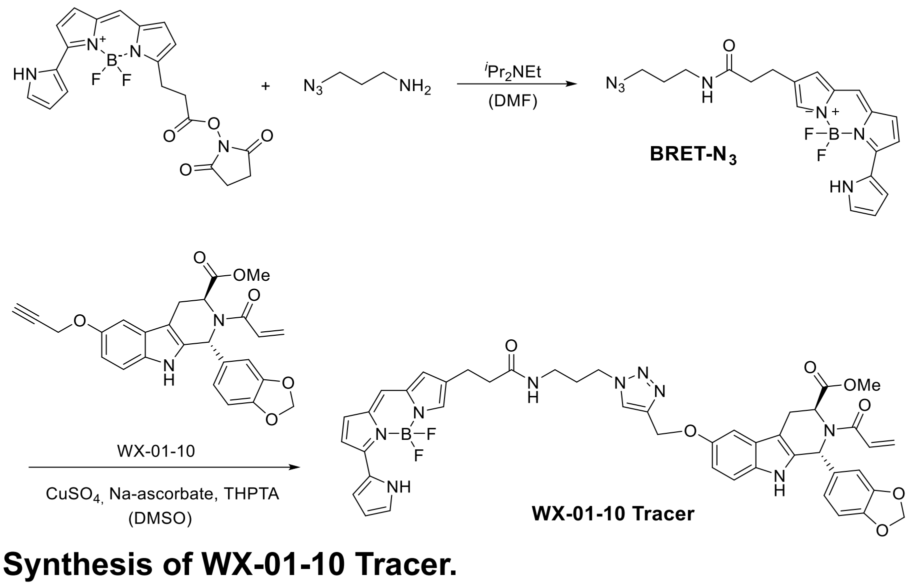

**Scheme 4. Synthesis of WX-01-10 Tracer.**

**Synthesis of WX-01-10 Tracer.** A portion of 3-azidopropan-1-amine (3.0 mg, 30.0 μmol) was dissolved in 0.2 mL DMF followed by the addition of BRET-dye-NHS (15.3 mg, 36.0 μmol) and DIPEA (16 μL, 89.9 μmol). The reaction was stirred at room temperature overnight. The product **BRET-N$_3$** (23% yield, 2.8 mg) was purified by HPLC as described in the **General Synthetic Methods**, with a linear gradient from 0-100% MeOH in water supplemented with 0.1% (v/v) TFA over 60 min. Next, to a solution of **WX-01-10** (2.5 mg) in *t*-BuOH (205 μL) and $H_2O$ (60 μL) were added a mixture of $CuSO_4{\cdot}5H_2O$ (327 μg in 32.7 μL $H_2O$), THPTA (0.95 mg in 20 μL $H_2O$), sodium ascorbate (972 μg in 97.2 μL $H_2O$), BRET-N$_3$ (2.7 mg, in 28 μL *t*-BuOH, 28 μL $H_2O$, and 28 μL DMSO). The mixture was prepared sequentially following the order above. The reaction was stirred at 37 °C for 5 h, protected from light. The product **WX-01-10 Tracer** (19% yield, 0.9 mg) was purified by HPLC as described in the **General Synthetic Methods**, with a linear gradient from 0-100% MeOH in water containing with 0.1% (v/v) TFA over 80 min, followed by a 20 min elution with 100% MeOH. $^1$H NMR (400 MHz, Methanol-*d*$_4$) δ 10.77 – 10.62 (m, 1H), 8.01 – 7.90 (m, 1H), 7.20 – 7.14 (m, 3H), 7.12 (d, *J* = 4.6 Hz, 1H), 7.08 (s, 1H), 7.05 (s, 1H), 6.97 (d, *J* = 4.6 Hz, 1H), 6.92 – 6.72 (m, 6H), 6.35 – 6.29 (m, 2H), 6.25 – 6.05 (m, 2H), 5.94 – 5.81 (m, 2H), 5.73 – 5.64 (m, 1H), 5.13 (s, 2H), 5.00 – 4.95 (m, 1H), 4.32 (t, *J* = 6.8 Hz, 2H), 3.65 – 3.56 (m, 3H), 3.29 – 3.11 (m, 6H), 2.63 (t, *J* = 7.5 Hz, 2H), 2.06 (p, *J* = 8.0, 7.1 Hz, 2H). HRMS (ESI) m/z: $[M + H]^+$ for $C_{45}H_{43}BF_2N_9O_7^+$, calculated 870.3341, found 870.3348.

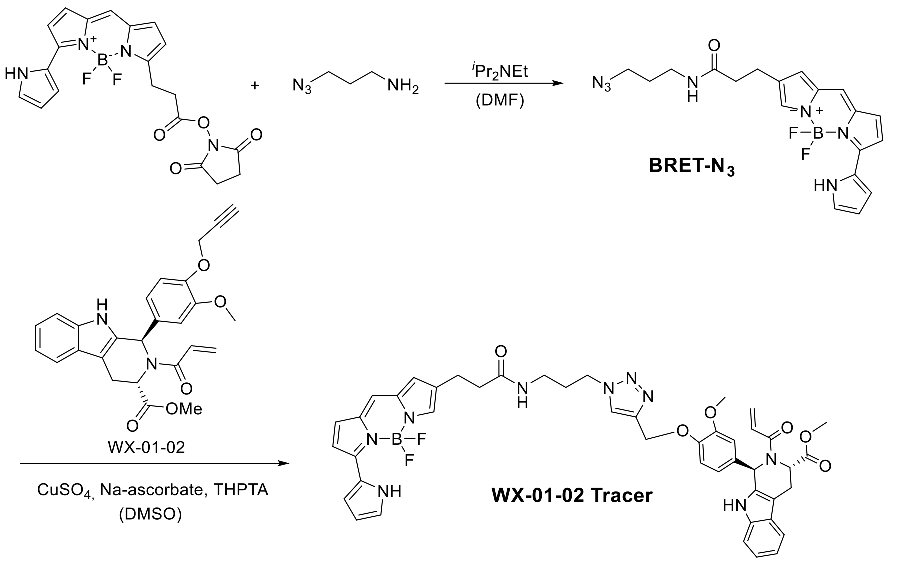


**Scheme 5. Synthesis of WX-01-02 Tracer.**

**Synthesis of WX-01-02 Tracer.** To a solution of **WX-01-02** (3.0 mg) in DMSO (125 µL) were added mixture of $CuSO_4{\cdot}5H_2O$ (410 µg in 20.5 µL $H_2O$), THPTA (3.6 mg in 36 µL $H_2O$), sodium ascorbate (1.2 mg in 60 µL $H_2O$), and **BRET-N$_3$** (3.4 mg). The mixture was prepared sequentially following the order above. The reaction was stirred at 37 °C, overnight protected from light. The product **WX-01-02 Tracer** (31% yield, 1.8 mg) was purified by HPLC as described in the **General Synthetic Methods**, with a linear gradient from 0-100% MeOH in water supplemented with 0.1% (v/v) TFA over 80 min, followed by 20 min methanol elution. $^1H$ NMR (400 MHz, Methanol-$d_4$) δ 7.92 – 7.82 (m, 1H), 7.45 (d, $J$ = 7.7 Hz, 1H), 7.34 – 6.96 (m, 10H), 6.95 – 6.74 (m, 4H), 6.34 – 6.28 (m, 2H), 6.26 – 6.14 (m, 1H), 5.69 (dd, $J$ = 10.6, 1.8 Hz, 1H), 5.11 – 5.06 (m, 2H), 4.97 (s, 1H), 4.30 (td, $J$ = 6.8, 1.5 Hz, 2H), 3.79 (s, 3H), 3.68 – 3.54 (m, 3H), 3.22 (dt, $J$ = 38.9, 7.6 Hz, 6H), 2.62 (t, $J$ = 7.6 Hz, 2H), 2.02 (dt, $J$ = 12.3, 5.5 Hz, 2H). HRMS (ESI) m/z: $[M + H]^+$ for $C_{45}H_{45}BF_2N_9O_6^+$, calculated 856.3548, found 856.3554.

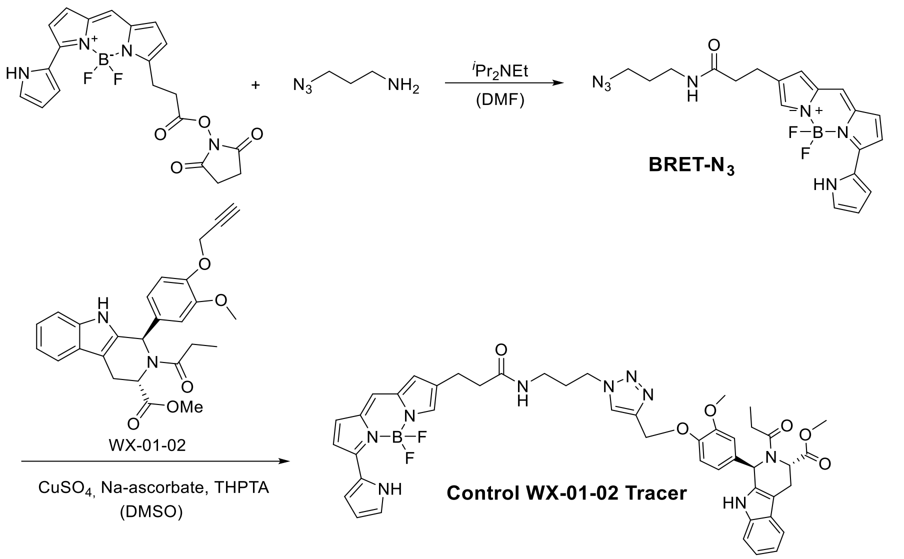


**Scheme 6. Synthesis of Control WX-01-02 Tracer.**

**Synthesis of WX-01-02 Tracer.** To a solution of **control WX-01-02** (2.5 mg) in DMSO (125 µL) were added mixture of $CuSO_4{\cdot}5H_2O$ (336 µg in 16.8 µL $H_2O$), THPTA (2.9 mg in 29 µL $H_2O$), sodium ascorbate (1.0 mg in 50 µL $H_2O$), and **BRET-N$_3$** (2.8 mg). The mixture was prepared sequentially following the order above. The reaction was stirred at 37 °C, overnight protected from light. The product **Control WX-01-02 Tracer** (21% yield, 1 mg) was purified by HPLC as described in the **General Synthetic Methods**, with a linear gradient from 0-100% MeOH in water supplemented with 0.1% (v/v) TFA over 80 min, followed by 20 min methanol elution. 1H NMR (400 MHz, Methanol-d4) δ 10.76 – 10.66 (m, 1H), 7.91 – 7.79 (m, 1H), 7.44 (d, J = 7.4 Hz, 1H), 7.35 – 6.73 (m, 14H), 6.31 (dd, J = 7.5, 2.9 Hz, 2H), 6.21 – 5.98 (m, 1H), 5.06 (dd, J = 20.7, 3.3 Hz, 3H), 4.33 – 4.25 (m, 2H), 3.78 (s, 3H), 3.59 (d, J = 20.6 Hz, 3H), 3.26 (d, J = 7.7 Hz, 2H), 3.16 (dd, J = 10.3, 6.2 Hz, 4H), 2.62 (t, J = 7.6 Hz, 2H), 2.55 (s, 1H), 2.38 – 2.27 (m, 1H), 2.07 – 1.95 (m, 2H), 1.10 – 0.96 (m, 3H). HRMS (ESI) m/z: $[M + H]^+$ for $C_{45}H_{45}BF_2N_9O_6^+$, calculated 856.3548, found 856.3554.

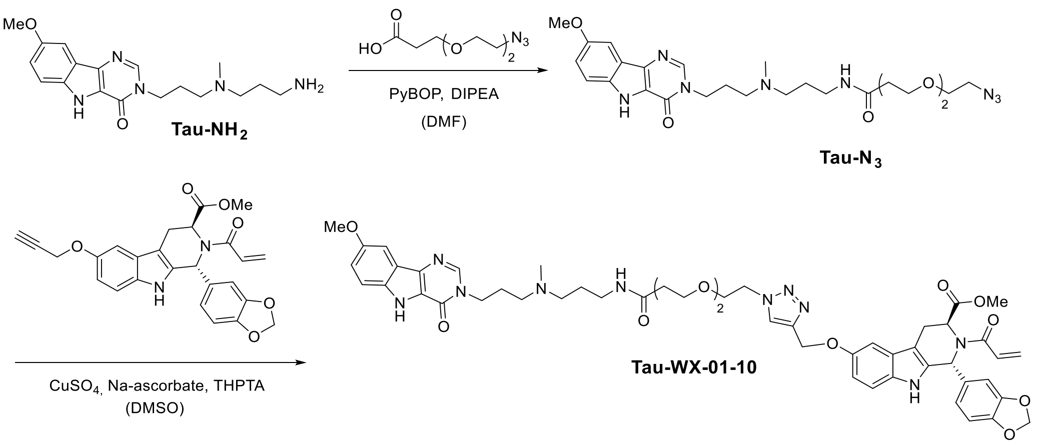


**Scheme 7. Synthesis of Tau-WX-01-10**.

**Synthesis of Tau-WX-01-10**. To solution of HOOC-PEG$_2$-N$_3$ (0.9 mL, 16.0 mg, 79.0 μmol) was added DIPEA (55 μL, 315.9 μmol) and PyBOP (82.2 mg, 157.9 μmol), and the reaction was stirred at room temperature for 10 min. Then, **Tau-NH$_2$** (20.0 mg, 52.6μmol) was added, and the reaction was incubated for another 3 h at room temperature. After 3h, LC-MS showed incomplete conversion of both starting materials. Thus, another 55 μL of DIPEA was added followed by overnight reaction at room temperature. The product **Tau-N$_3$** (61% yield, 16.9 mg) was purified by HPLC as described in the **General Synthetic Methods**, with a linear gradient from 0-100% MeOH in water supplemented with 0.1% (v/v) TFA over 60 min.

Next, to a solution of **WX-01-10** (2.5 mg) in DMSO (125 μL) were added mixture of THPTA (1.9 mg in 40 μL $H_2O$), $CuSO_4·5H_2O$ (218 μg in 20.8 μL $H_2O$), sodium ascorbate (648 μg in 64.8 μL $H_2O$), and **Tau-N$_3$** (3.5 mg). The mixture was prepared sequentially following the order above. The reaction was stirred at 37°C overnight. The product **Tau-WX-01-10** (46% yield, 2.5 mg) was purified by HPLC as described in the **General Synthetic Methods**, with a linear gradient from 0-100% methanol in water supplemented with 0.1% (v/v) TFA over 80 min. $^1$H NMR (400 MHz, Methanol-*d*$_4$) δ 8.20 (s, 1H), 8.05 (s, 1H), 7.55 – 7.41 (m, 2H), 7.23 – 7.04 (m, 3H), 6.94 – 6.70 (m, 5H), 6.29 – 6.05 (m, 2H), 5.98 – 5.79 (m, 2H), 5.70 (dd, *J* = 10.6, 1.8 Hz, 1H), 5.15 (s, 2H), 4.93 (s, 1H), 4.57 – 4.52 (m, 2H), 4.17 (t, *J* = 6.6 Hz, 2H), 3.90 – 3.81 (m, 5H), 3.64 – 3.45 (m, 8H), 3.36 (d, *J* = 5.7 Hz, 5H), 3.21 – 3.03 (m, 6H), 2.84 – 2.76 (m, 3H), 2.33 (q, *J* = 5.5 Hz, 2H), 2.25 – 2.17 (m, 2H). HRMS (ESI) m/z: $[M + H]^+$ for $C_{51}H_{59}N_{10}O_{11}^+$, calculated 987.4359, found 987.4356.

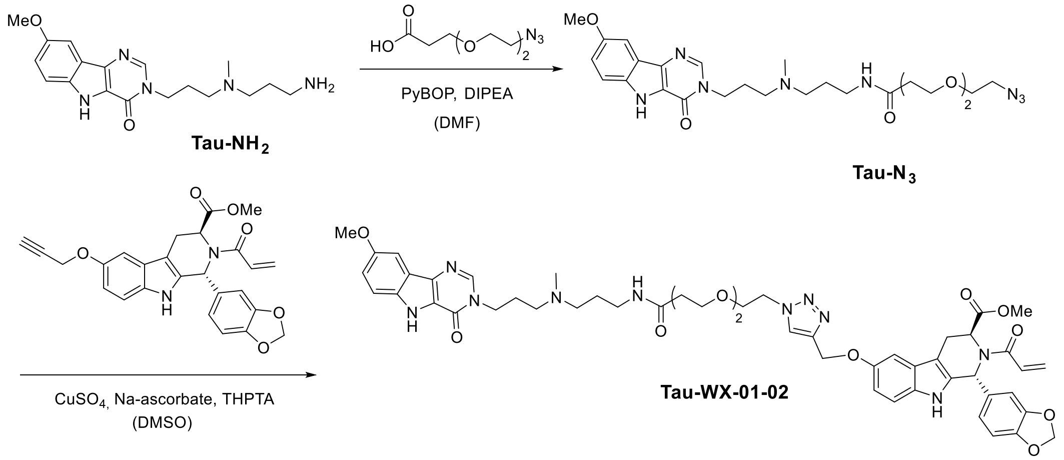


**Scheme 8. Synthesis of Tau-WX-01-02**.

**Synthesis of Tau-WX-01-02**. To a solution of **WX-01-02** (2.5 mg) in DMSO (125 µL) were added mixture of THPTA (3.0 mg in 30 µL $H_2O$), $CuSO_4{\cdot}5H_2O$ (338 µg in 16.9 µL $H_2O$), sodium ascorbate (1 mg in 50 µL $H_2O$), and **Tau-N$_3$** (3.6 mg). The mixture was prepared sequentially following the order above. The reaction was stirred at 37 °C overnight. The product **Tau-WX-01-02** (16% yield, 0.9 mg) was purified by HPLC as described in the **General Synthetic Methods**, with a linear gradient from 0-100% MeOH in water containing with 0.1% (v/v) TFA over 80 min. 1H NMR (400 MHz, Methanol-*d*4) δ 8.16 (s, 1H), 8.02 (s, 1H), 7.55 – 7.50 (m, 1H), 7.44 (dd, *J* = 20.1, 8.4 Hz, 2H), 7.32 – 7.24 (m, 1H), 7.24 – 7.14 (m, 2H), 7.10 – 6.89 (m, 5H), 6.83 – 6.66 (m, 1H), 6.28 – 6.10 (m, 2H), 5.73 – 5.66 (m, 1H), 5.10 (s, 3H), 4.58 – 4.44 (m, 2H), 4.15 (t, *J* = 7.0 Hz, 2H), 3.86 (s, 3H), 3.80 (d, *J* = 7.7 Hz, 5H), 3.67 – 3.42 (m, 10H), 3.35 (s, 3H), 3.18 – 3.01 (m, 4H), 2.80 – 2.68 (m, 3H), 2.37 – 2.26 (m, 2H), 2.24 – 2.11 (m, 2H), 1.89 – 1.70 (m, 2H). HRMS (ESI) m/z: $[M + H]^+$ for $C_{51}H_{61}N_{10}O_{10}^+$, calculated 973.4567, found 973.4564.

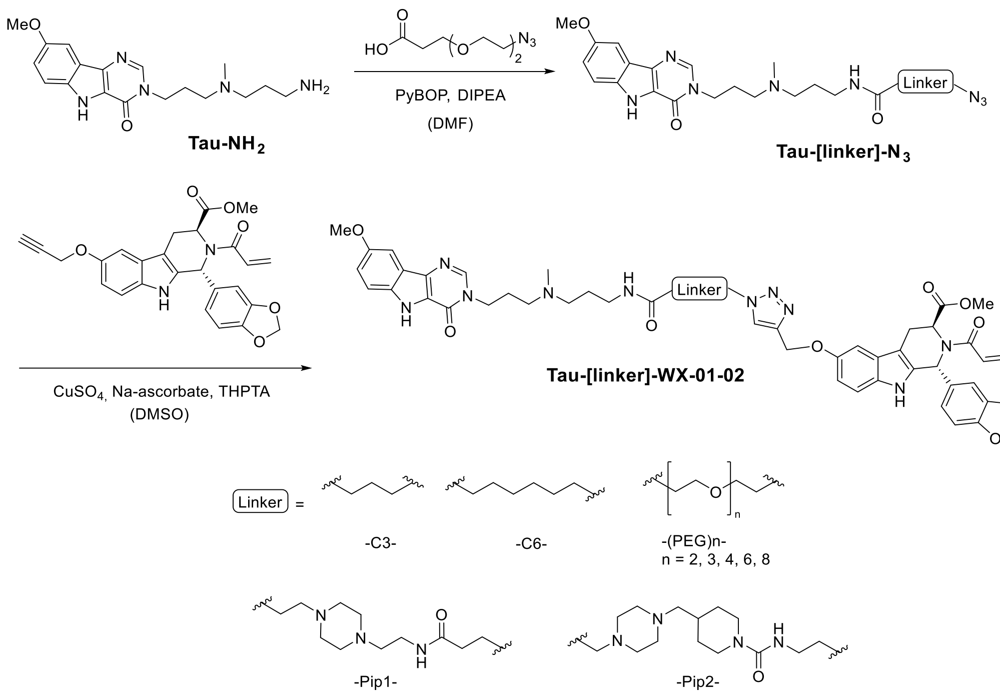


**Scheme 9. Synthesis of Tau-[linker]-WX-01-02**.

**Syntheses of Tau-[linker]-WX-01-02**. The syntheses of **Tau-[linker]-N**$_3$ was similarly performed as described in **Scheme 6**. In brief, to a solution of HOOC-C3/C6/Pip1/Pip2/PEG$_3$/PEG$_4$/PEG$_6$/PEG$_8$-N$_3$ (0.9 mL; 79.0 µmol) was added DIPEA (110 µL, 631.8 µmol) and PyBOP (82.2 mg, 157.9 µmol). The reaction was stirred at room temperature for 10 min, following by addition of **Tau-NH**$_2$ (20.0 mg, 52.6µmol), after which the reaction was stirred for another 3 h. The product **Tau-[linker]-N**$_3$ was purified by HPLC as described in the **General Synthetic Methods**, with a linear gradient from 0-100% MeOH in water supplemented with 0.1% (v/v) TFA over 60 min.

Next, to a solution of **WX-01-02** (2.5 mg, 5.6 µmol) in DMSO (125 µL) was added a mixture of THPTA (3.0 mg in 30 µL $H_2O$, 6.8 µmol), $CuSO_4 \cdot 5H_2O$ (338 µg in 16.9 µL $H_2O$, 1.4 µmol), sodium ascorbate (1 mg in 50 µL $H_2O$, 5.1 µmol), and **Tau-[linker]-N**$_3$ (6.8 µmol). The mixture was prepared sequentially following the order above, and stirred overnight at 37 °C. The products **Tau-[linker]-WX-01-02** were purified by HPLC as described in the **General Synthetic Methods**, with a linear gradient from 0-100% MeOH in water supplemented with 0.1% (v/v) TFA over 80 min. **Tau-C3-WX-01-02:** HRMS (ESI) m/z: [M + H]$^+$ for $C_{48}H_{55}N_{10}O_8^+$, calculated 899.4199, found 899.4194. **Tau-C6-WX-01-02:** $^1$H NMR (400 MHz, Methanol-*d*$_4$) δ 8.23 (s, 1H), 8.00 – 7.84 (m, 1H), 7.58 – 7.40 (m, 3H), 7.36 – 7.21 (m, 1H), 7.16 (dd, *J* = 9.0, 2.3 Hz, 1H), 7.12 – 6.88 (m, 5H), 6.85 – 6.72 (m, 1H), 6.33 – 6.11 (m, 2H), 5.75 – 5.66 (m, 1H), 5.19 – 4.95 (m, 3H), 4.26 (dq, *J* = 19.5, 6.6 Hz, 4H), 3.87 (s, 3H), 3.82 (s, 3H), 3.70 – 3.58 (m, 3H), 3.58 – 3.34 (m, 3H), 3.31 – 3.02 (m, 7H), 2.88 (s, 3H), 2.32 – 2.19 (m, 2H), 2.13 (t, *J* = 7.3 Hz, 2H), 1.99 – 1.87 (m, 2H),

1.79 (q, $J$ = 7.0 Hz, 2H), 1.51 (p, $J$ = 7.3 Hz, 2H), 1.19 (ddd, $J$ = 22.6, 12.0, 5.1 Hz, 4H). HRMS (ESI) m/z: $[M + H]^+$ for $C_{51}H_{61}N_{10}O_8^+$, calculated 941.4668, found 941.4663. **Tau-Pip1-WX-01-02:** $^1$H NMR (600 MHz, Methanol-$d_4$) δ 8.24 (s, 1H), 8.04 – 7.95 (m, 1H), 7.53 (d, $J$ = 2.3 Hz, 1H), 7.46 (dd, $J$ = 25.0, 8.4 Hz, 2H), 7.32 – 7.20 (m, 1H), 7.17 (dd, $J$ = 9.0, 2.2 Hz, 1H), 7.10 – 6.74 (m, 6H), 6.31 – 6.10 (m, 2H), 5.70 (d, $J$ = 10.5 Hz, 1H), 5.12 (s, 3H), 4.43 (t, $J$ = 6.3 Hz, 2H), 4.27 – 4.21 (m, 2H), 3.87 (s, 3H), 3.81 (s, 3H), 3.63 (s, 3H), 3.39 – 3.36 (m, 3H), 3.22 – 3.17 (m, 3H), 3.11 – 3.06 (m, 3H), 2.90 – 2.76 (m, 7H), 2.66 (s, 9H), 2.57 (t, $J$ = 6.5 Hz, 2H), 2.29 – 2.13 (m, 7H), 1.95 – 1.87 (m, 2H). HRMS (ESI) m/z: $[M + H]^+$ for $C_{57}H_{72}N_{13}O_9^+$, calculated 1082.5570, found 1082.5566. **Tau-Pip2-WX-01-02:** $^1$H NMR (600 MHz, Methanol-$d_4$) δ 8.29 (s, 1H), 8.03 – 7.93 (m, 1H), 7.57 – 7.51 (m, 1H), 7.47 (dd, $J$ = 28.3, 8.6 Hz, 2H), 7.26 (dd, $J$ = 33.5, 9.6 Hz, 1H), 7.17 (dd, $J$ = 9.0, 2.4 Hz, 1H), 7.01 (ddd, $J$ = 46.1, 38.2, 7.9 Hz, 5H), 6.78 (dd, $J$ = 15.4, 11.1 Hz, 1H), 6.33 – 6.04 (m, 2H), 5.70 (d, $J$ = 10.7 Hz, 1H), 5.19 – 5.10 (m, 3H), 4.51 – 4.41 (m, 4H), 4.30 – 4.24 (m, 2H), 3.88 (s, 3H), 3.82 (s, 3H), 3.63 (s, 3H), 3.48 – 3.41 (m, 3H), 3.26 – 3.10 (m, 9H), 2.92 (d, $J$ = 15.0 Hz, 7H), 2.30 (s, 3H), 2.18 – 2.10 (m, 2H), 2.06 – 1.95 (m, 3H), 1.73 (d, $J$ = 11.2 Hz, 2H), 1.14 (dd, $J$ = 30.4, 14.1 Hz, 2H). HRMS (ESI) m/z: $[M + H]^+$ for $C_{59}H_{74}N_{14}O_9^+$, calculated 1122.5759, found 1122.5885. **Tau-PEG$_3$-WX-01-02:** HRMS (ESI) m/z: $[M + H]^+$ for $C_{53}H_{65}N_{10}O_{11}^+$, calculated 1017.4829, found 1017.4828. **Tau-PEG$_4$-WX-01-02:** $^1$H NMR (600 MHz, Methanol-$d_4$) δ 8.20 (d, $J$ = 3.9 Hz, 1H), 8.01 (d, $J$ = 12.1 Hz, 1H), 7.52 (s, 1H), 7.45 (dd, $J$ = 26.8, 8.5 Hz, 2H), 7.32 – 7.24 (m, 1H), 7.20 – 7.12 (m, 1H), 7.09 – 6.90 (m, 5H), 6.83 – 6.71 (m, 1H), 6.30 – 6.09 (m, 2H), 5.69 (d, $J$ = 10.7 Hz, 1H), 5.10 (s, 3H), 4.51 (s, 2H), 4.19 (s, 3H), 3.86 (s, 3H), 3.80 (s, 5H), 3.59 – 3.44 (m, 10H), 3.42 (s, 5H), 3.37 (dd, $J$ = 12.9, 8.4 Hz, 6H), 3.17 (dd, $J$ = 18.4, 8.1 Hz, 4H), 2.80 (s, 3H), 2.36 – 2.28 (m, 2H), 2.26 – 2.17 (m, 2H), 1.89 – 1.76 (m, 2H). HRMS (ESI) m/z: $[M + H]^+$ for $C_{55}H_{69}N_{10}O_{12}^+$, calculated 1061.5091, found 1061.5083. **Tau-PEG$_6$-WX-01-02:** HRMS (ESI) m/z: $[M + H]^+$ for $C_{59}H_{77}N_{10}O_{14}^+$, calculated 1149.5615, found 1149.5613. **Tau-PEG$_8$-WX-01-02:** HRMS (ESI) m/z: $[M + H]^+$ for $C_{63}H_{85}N_{10}O_{16}^+$, calculated 1237.6140, found 1237.6140.

#### COMPOUND CHARACTERIZATION
#### $^1$H NMR, $^{13}$C NMR, Analytical HPLC, and HR-MS Spectra

##### Tau-RiboTether

$^1$H NMR

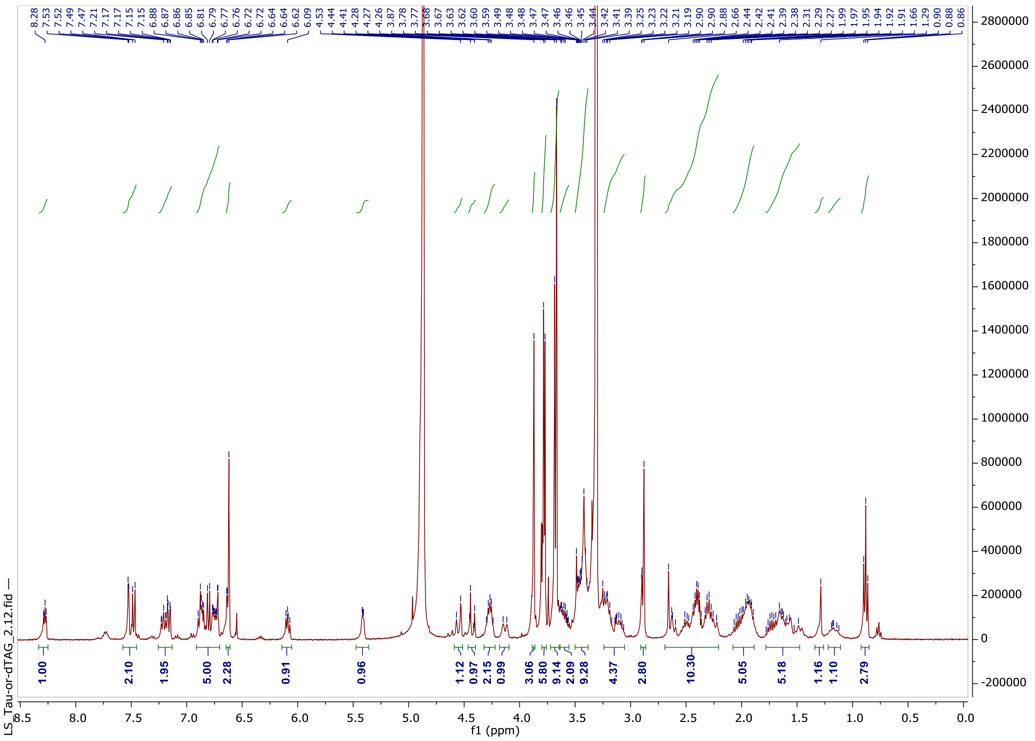

$^{13}$C NMR

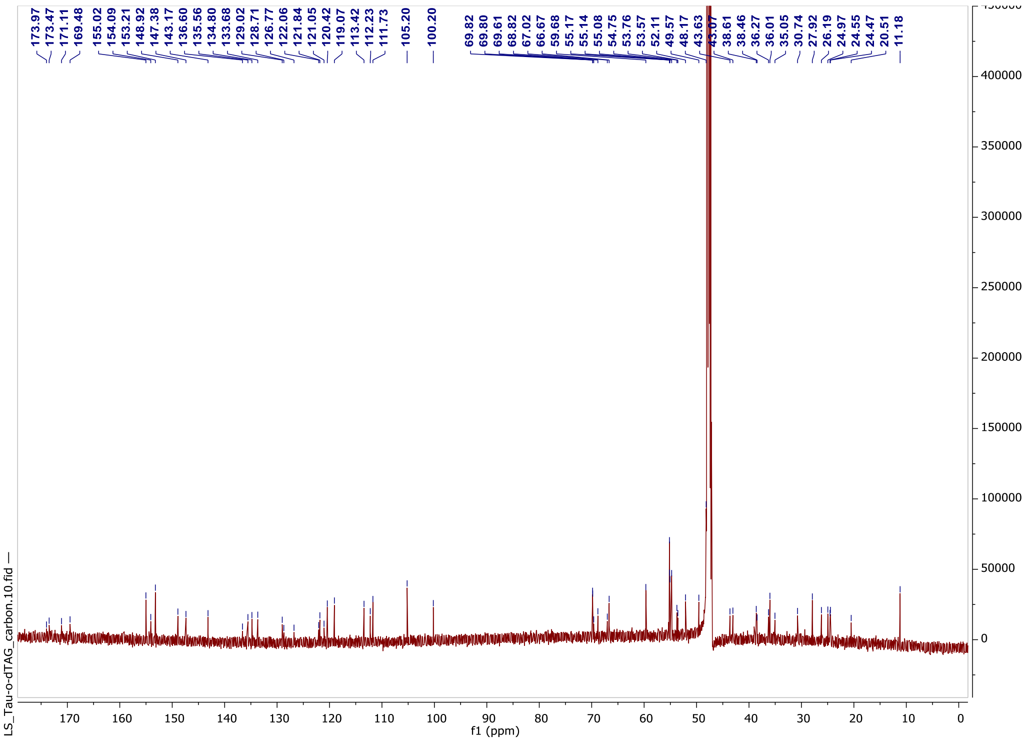

Analytical HPLC

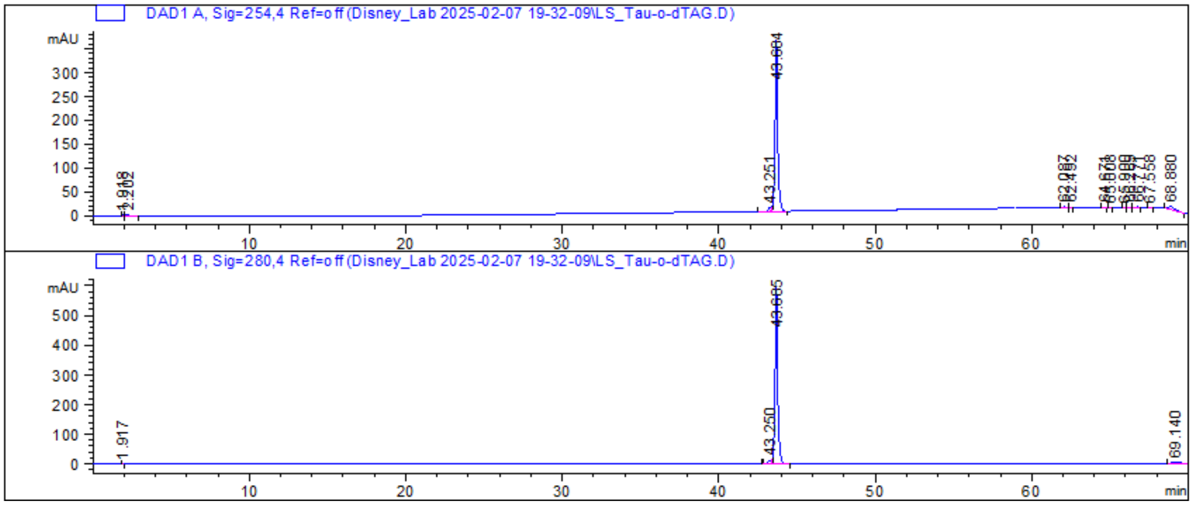


HRMS

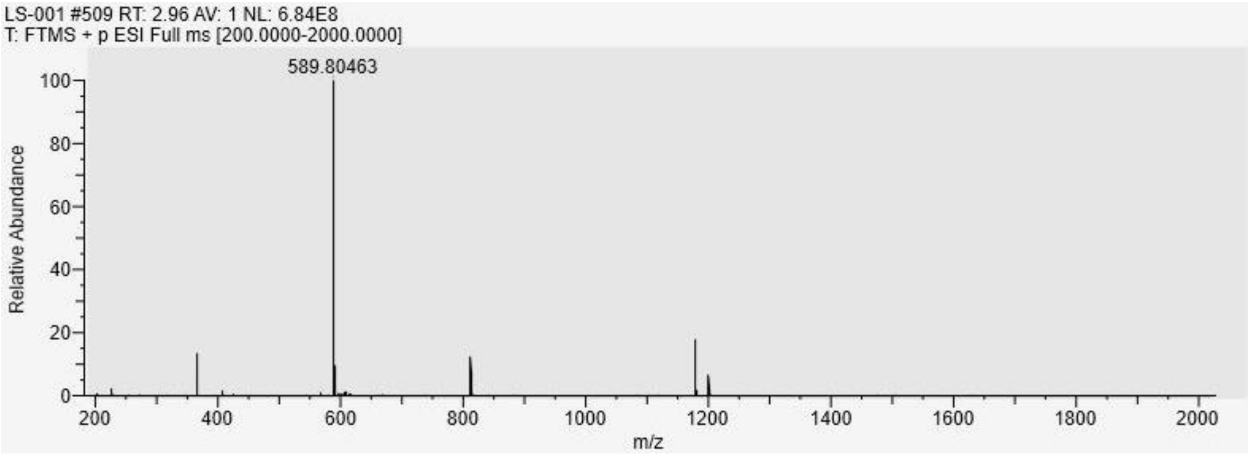

#### Tau-binder

##### $^1$H NMR

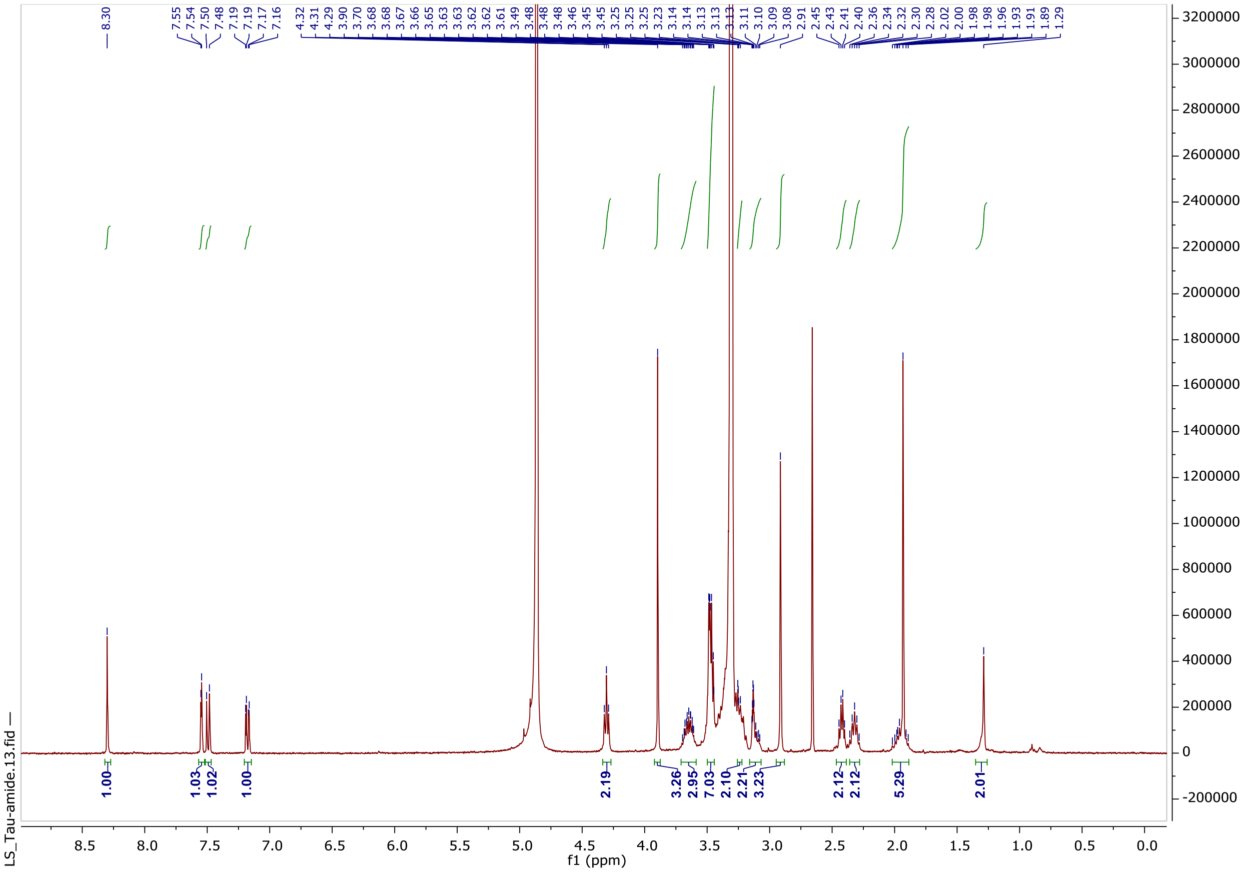

##### Analytical HPLC

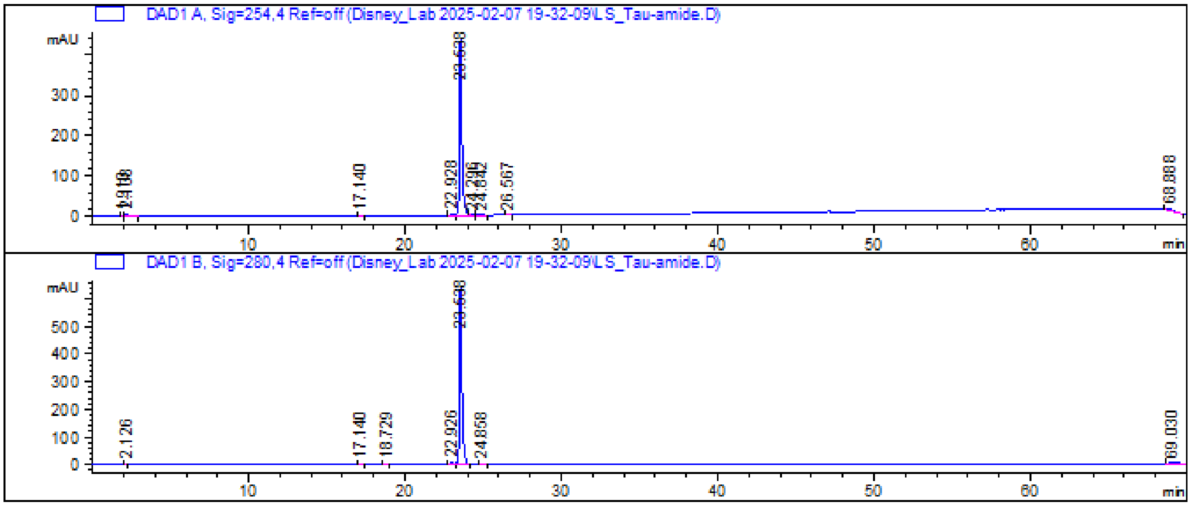

##### HRMS

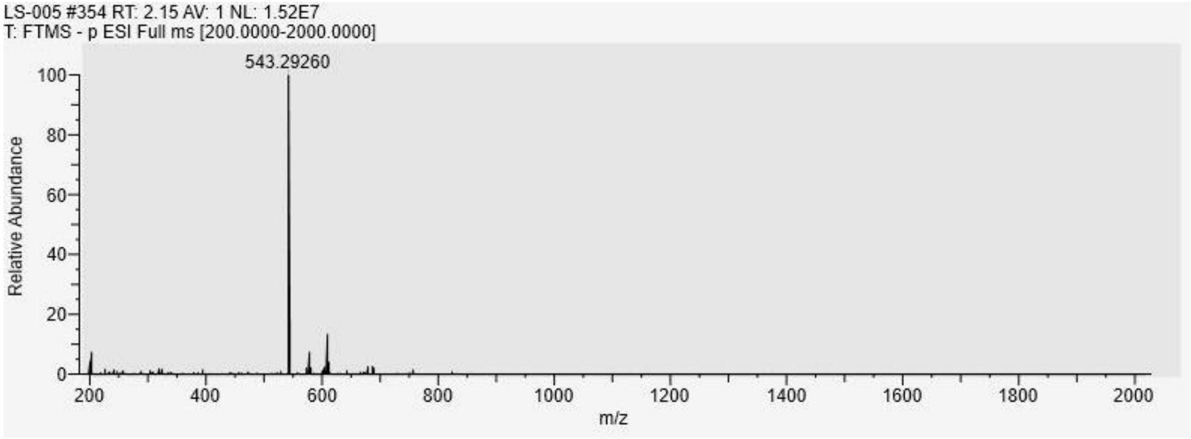
LS-005 #354 RT: 2.15 AV: 1 NL: 1.52E7
T: FTMS - p ESI Full ms [200.0000-2000.0000]
543.29260
Relative Abundance
100
80
60
40
20
0
200
400
600
800
1000
1200
1400
1600
1800
2000
m/z

### Control WX-01-02

$^1$H NMR

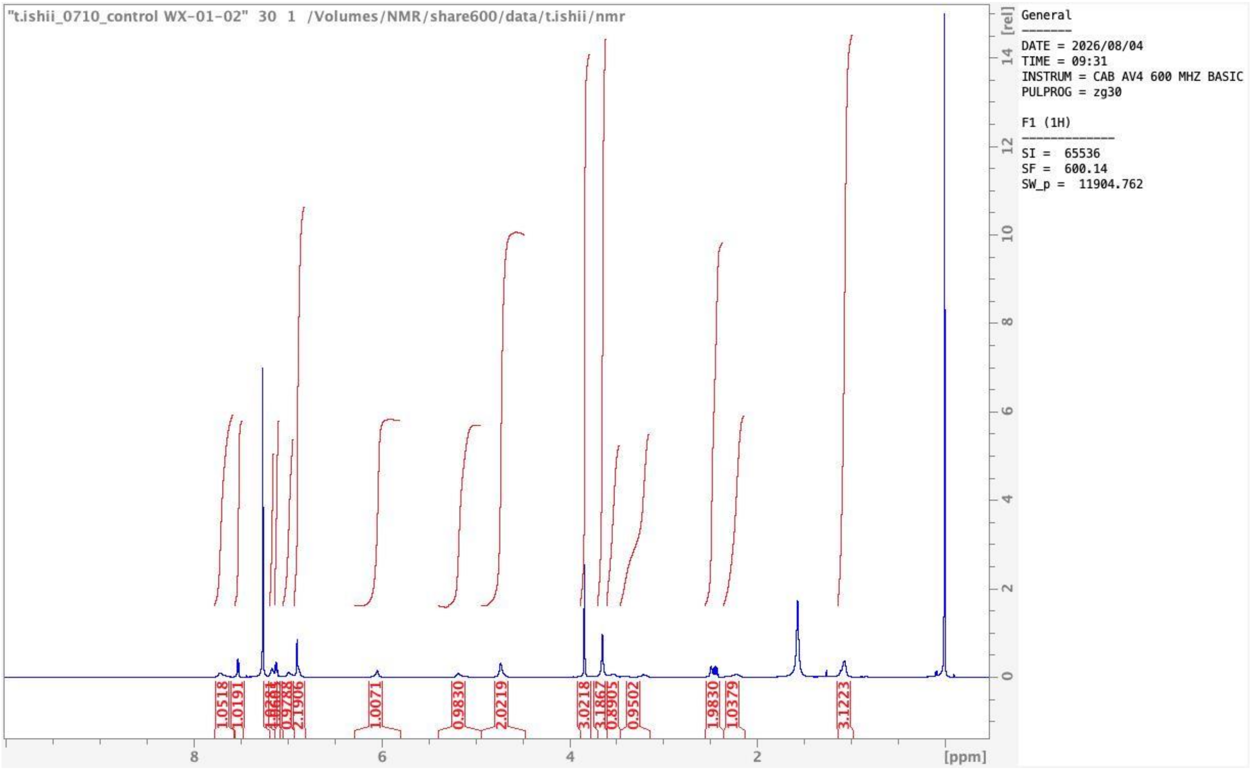


#### Analytical HPLC

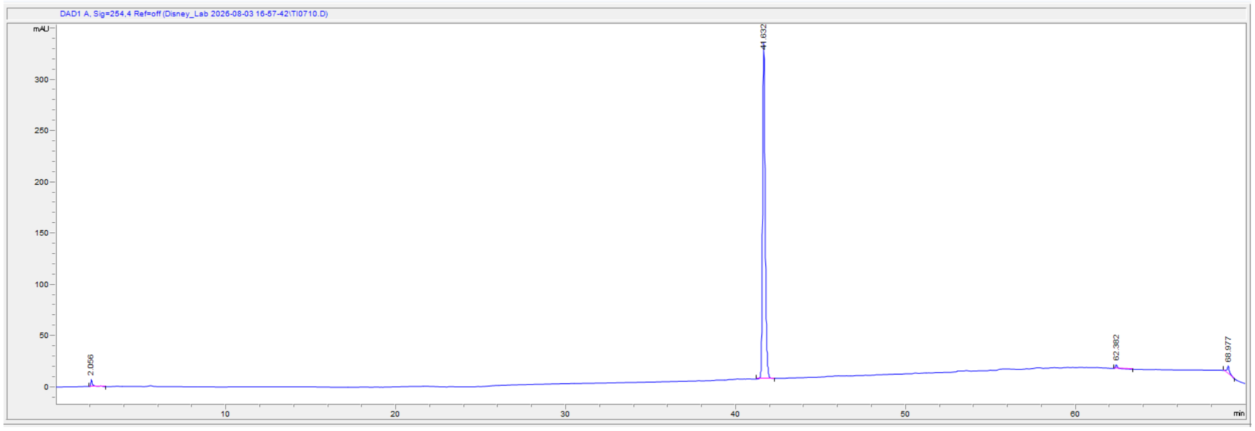

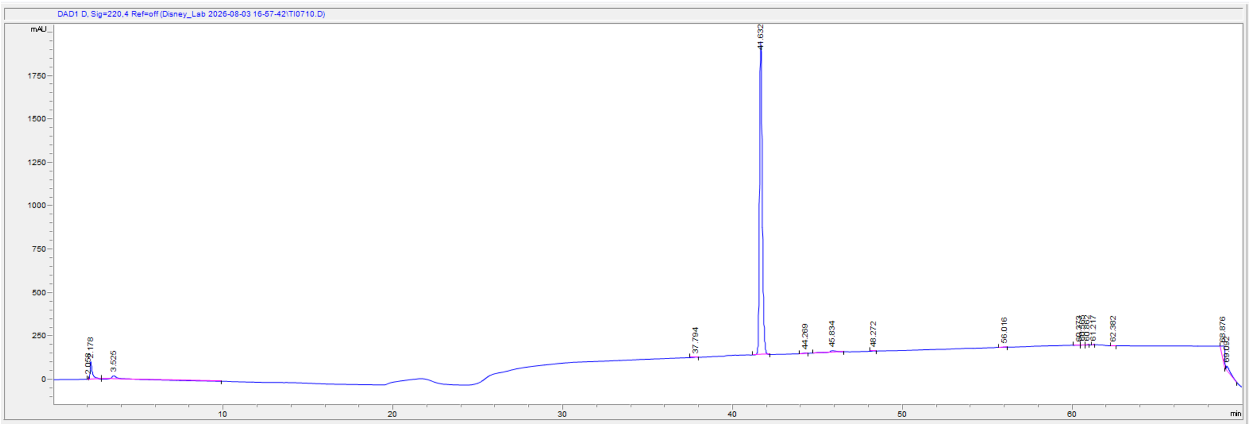
DAD1 D, Sig=220,4 Ref=off (Disney_Lab 2026-08-03 16-57-42\TI0710.D)
mAU
41.632
min

#### HRMS

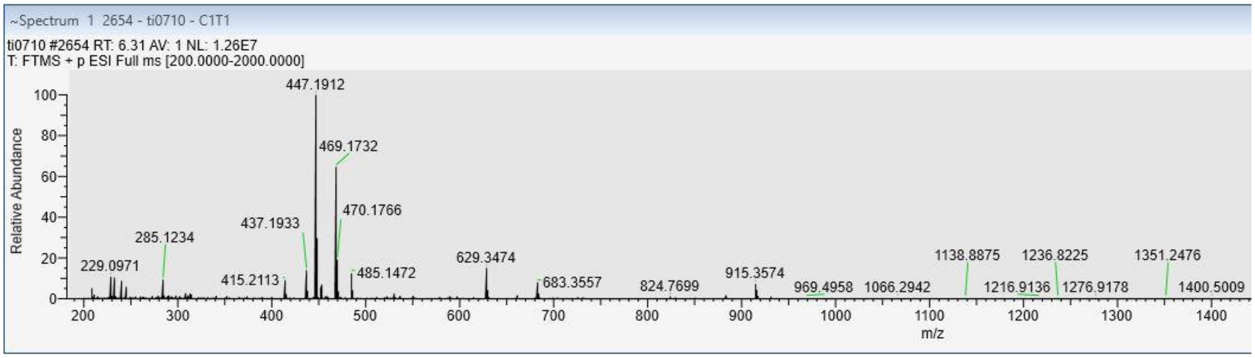
~Spectrum 1 2654 - ti0710 - C1T1
ti0710 #2654 RT: 6.31 AV: 1 NL: 1.26E7
T: FTMS + p ESI Full ms [200.0000-2000.0000]
Relative Abundance
447.1912
469.1732
470.1766
437.1933
285.1234
229.0971
415.2113
485.1472
629.3474
683.3557
824.7699
915.3574
969.4958
1066.2942
1138.8875
1216.9136
1236.8225
1276.9178
1351.2476
1400.5009
m/z

#### WX-01-10 Tracer

##### $^{1}H$ NMR

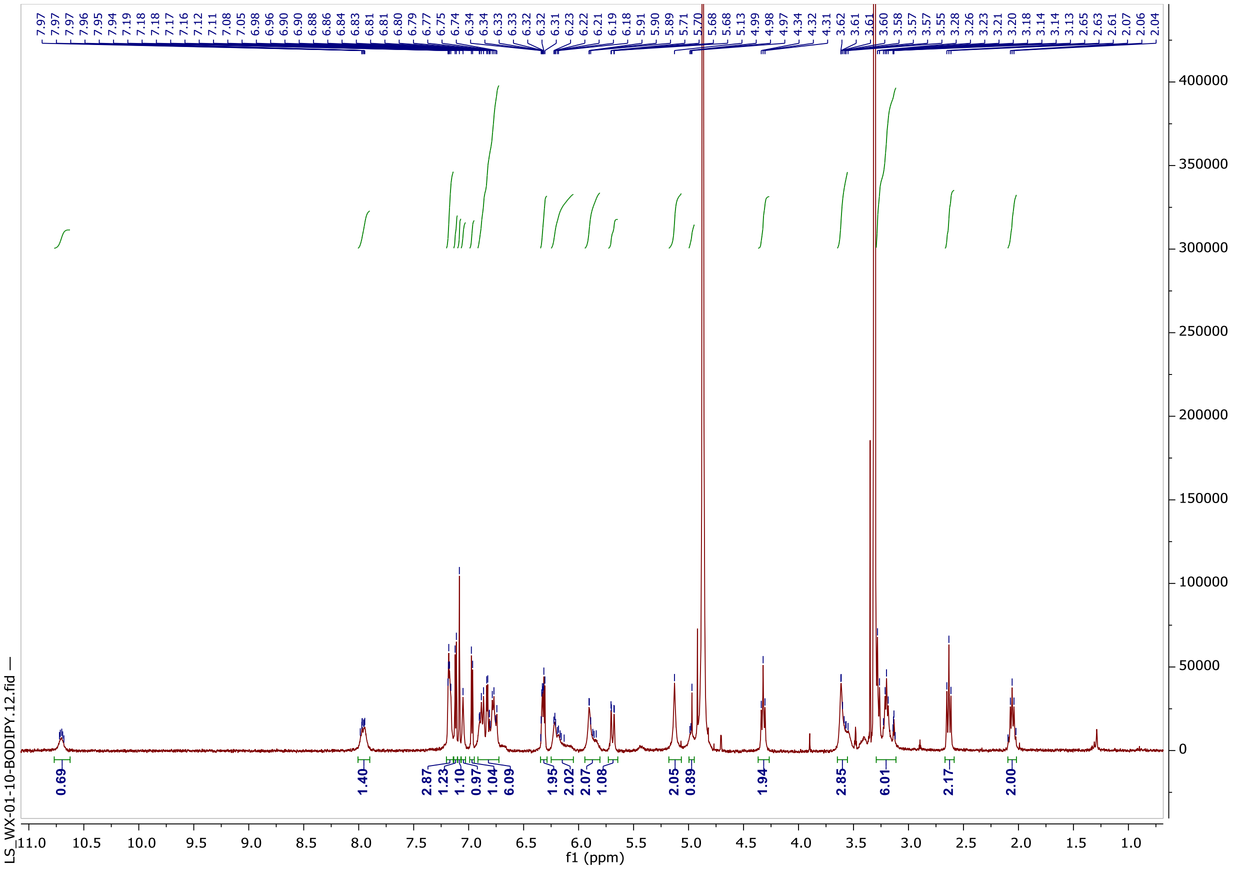

##### Analytical HPLC

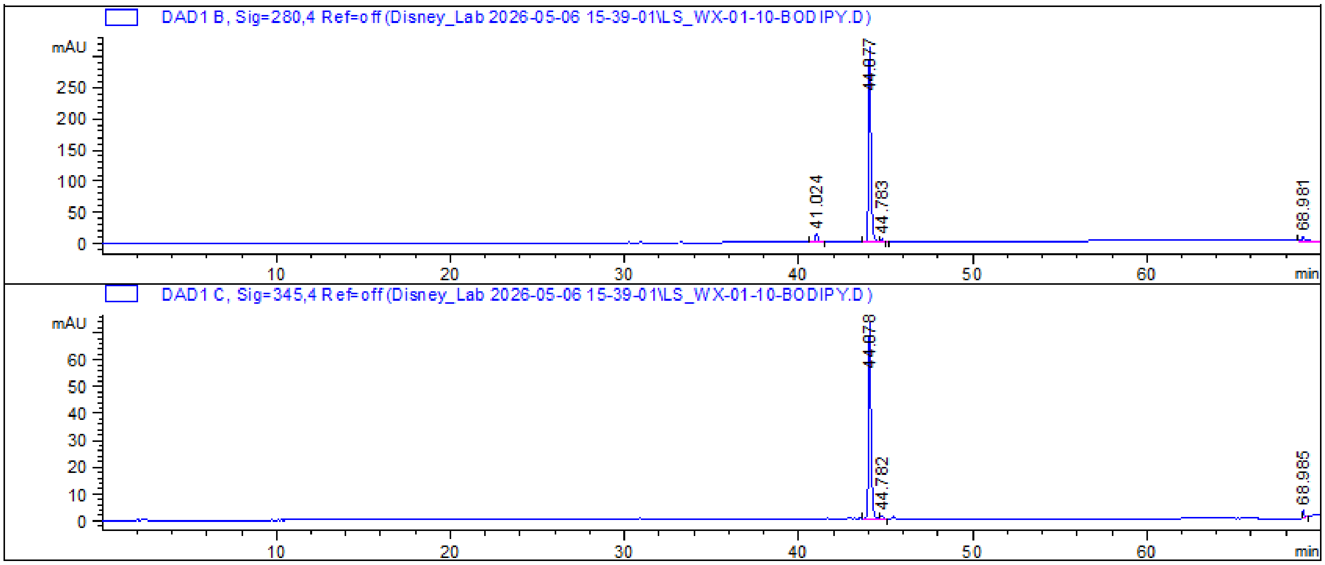

HRMS

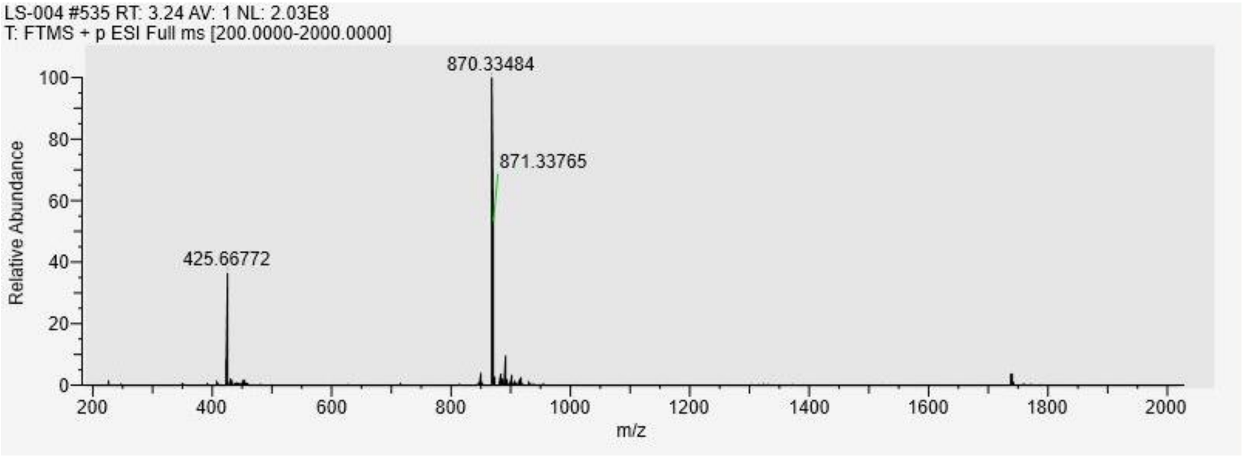

LS-004 #535 RT: 3.24 AV: 1 NL: 2.03E8
T: FTMS + p ESI Full ms [200.0000-2000.0000]
870.33484
871.33765
425.66772
Relative Abundance
m/z

#### WX-01-02 Tracer

##### $^1$H NMR

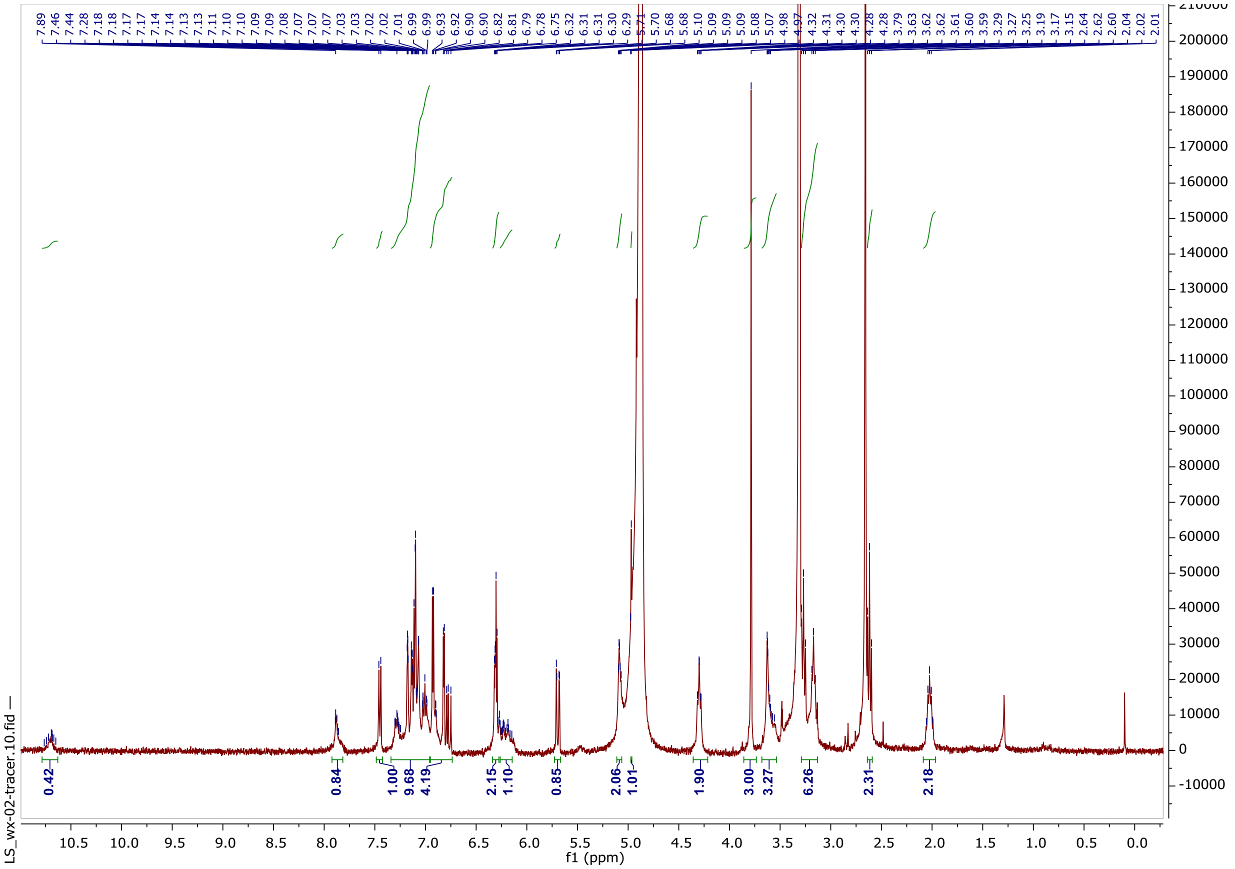


##### Analytical HPLC

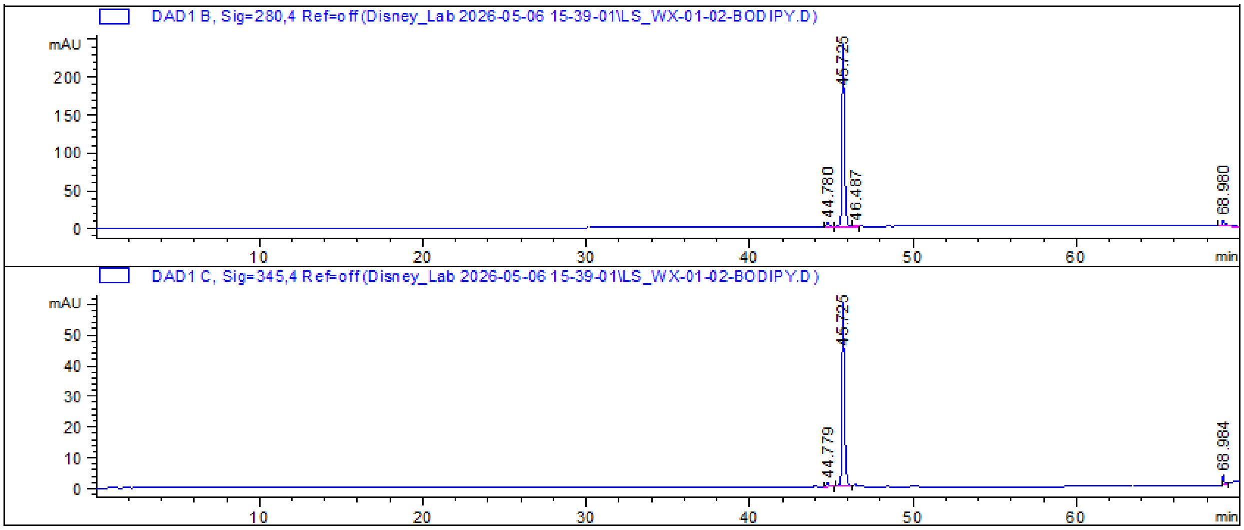

HRMS

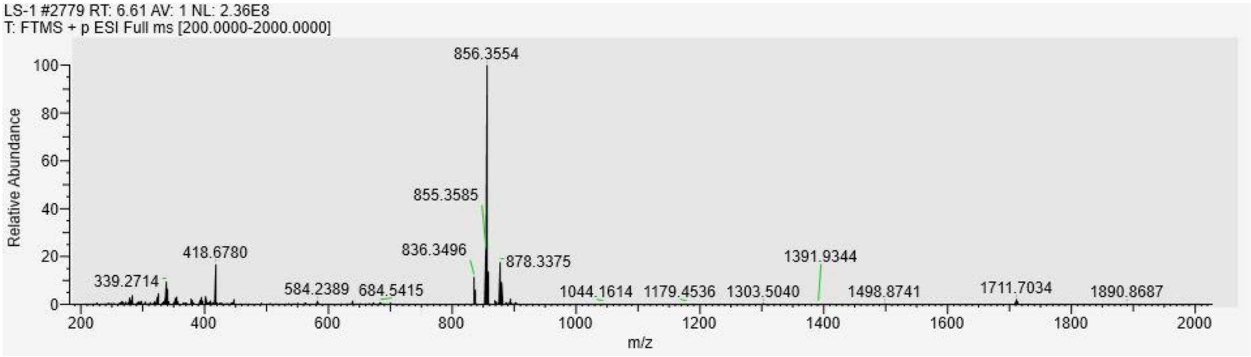
LS-1 #2779 RT: 6.61 AV: 1 NL: 2.36E8
T: FTMS + p ESI Full ms [200.0000-2000.0000]
856.3554
855.3585
836.3496
878.3375
418.6780
339.2714
584.2389
684.5415
1044.1614
1179.4536
1303.5040
1391.9344
1498.8741
1711.7034
1890.8687
Relative Abundance
m/z

#### Control WX-01-02 Tracer

##### $^{1}$H NMR

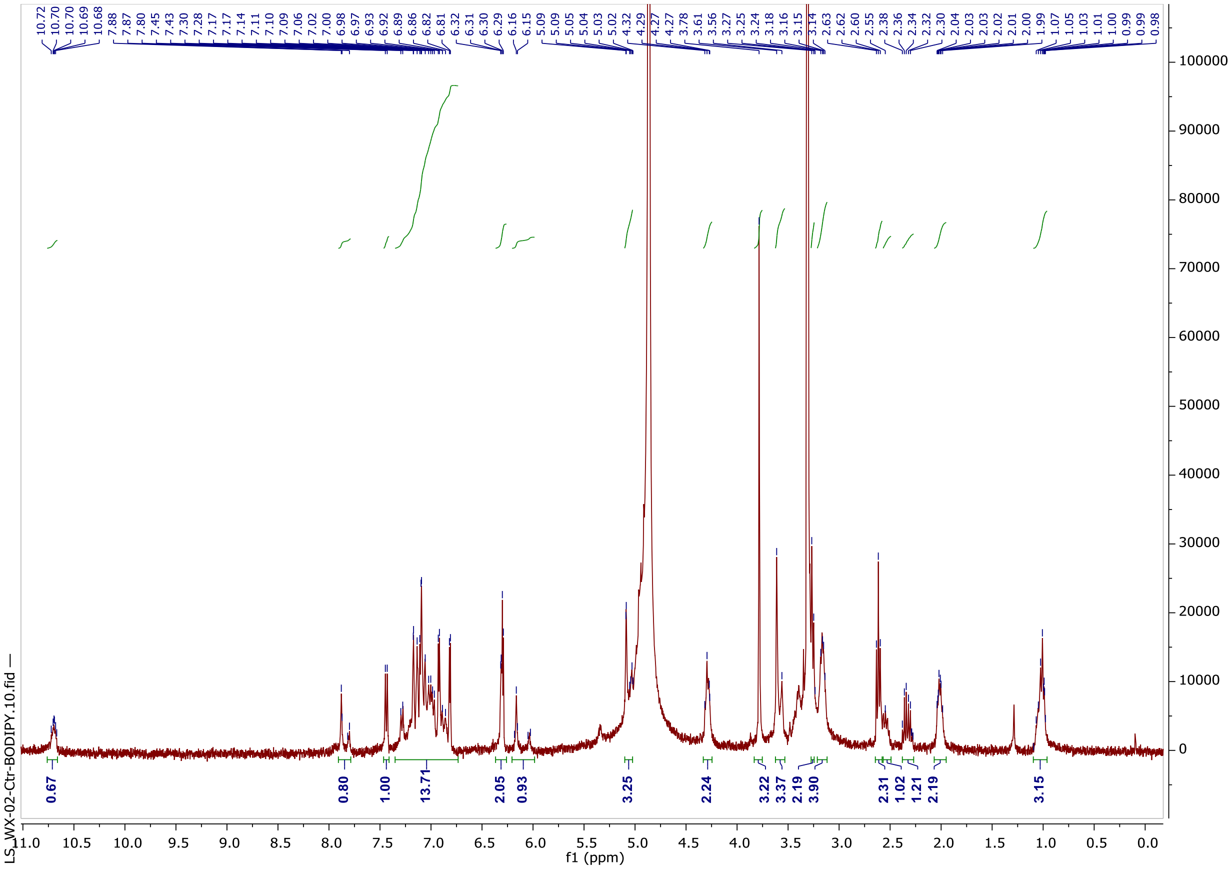

##### Analytical HPLC

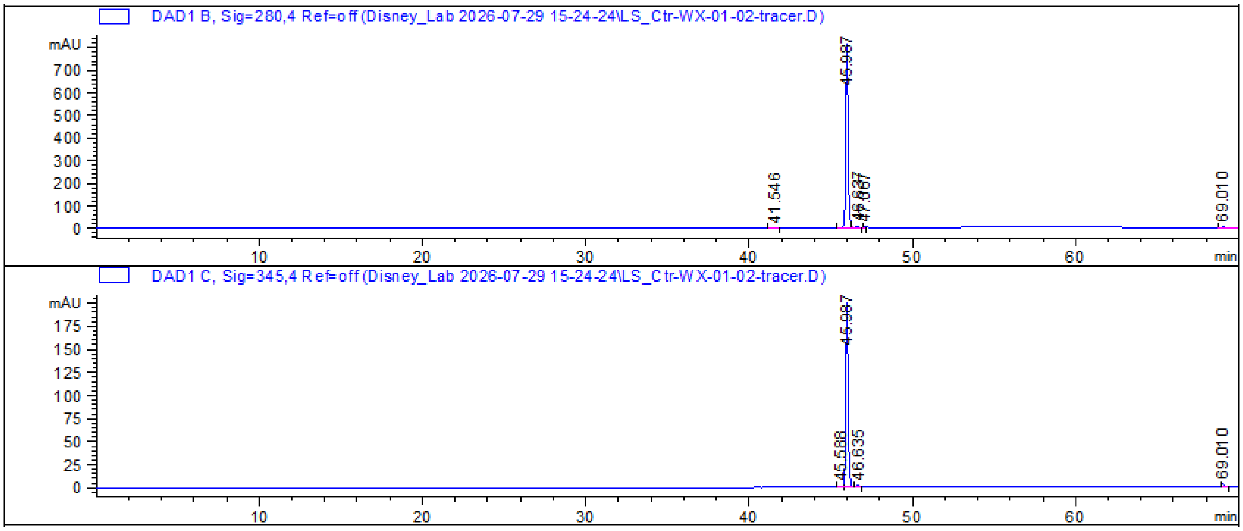

HRMS

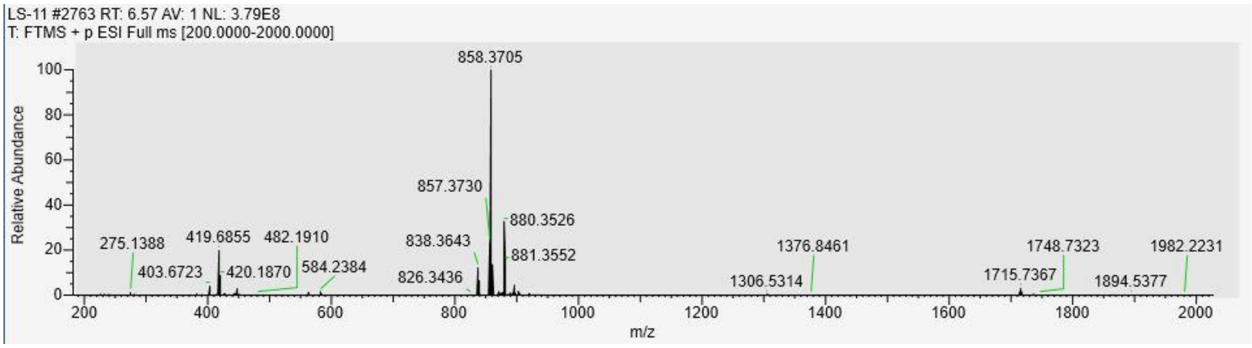
LS-11 #2763 RT: 6.57 AV: 1 NL: 3.79E8
T: FTMS + p ESI Full ms [200.0000-2000.0000]
858.3705
857.3730
880.3526
881.3552
838.3643
826.3436
275.1388
419.6855
482.1910
403.6723
420.1870
584.2384
1376.8461
1306.5314
1748.7323
1715.7367
1982.2231
1894.5377
Relative Abundance
m/z

#### Tau-WX-01-02

##### $^1$H NMR

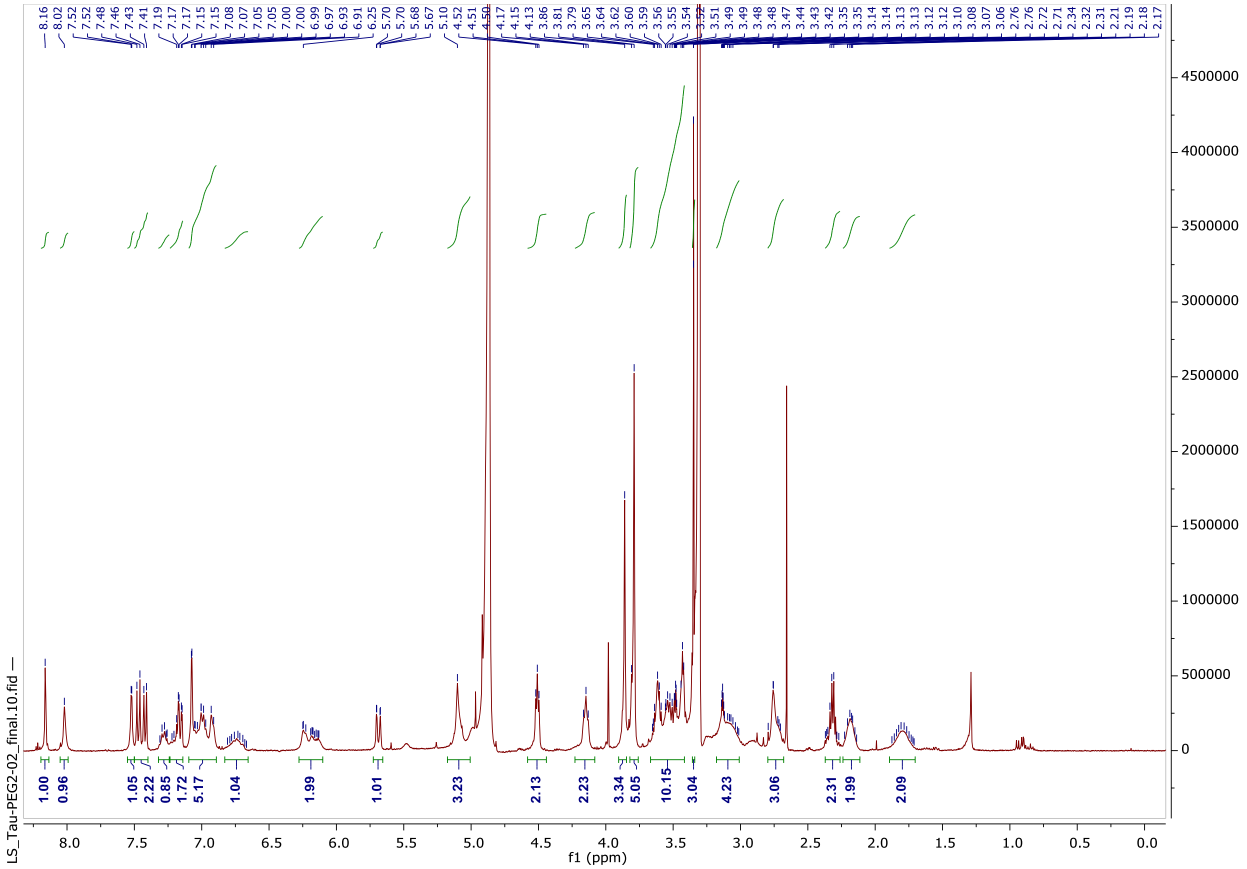

##### Analytical HPLC

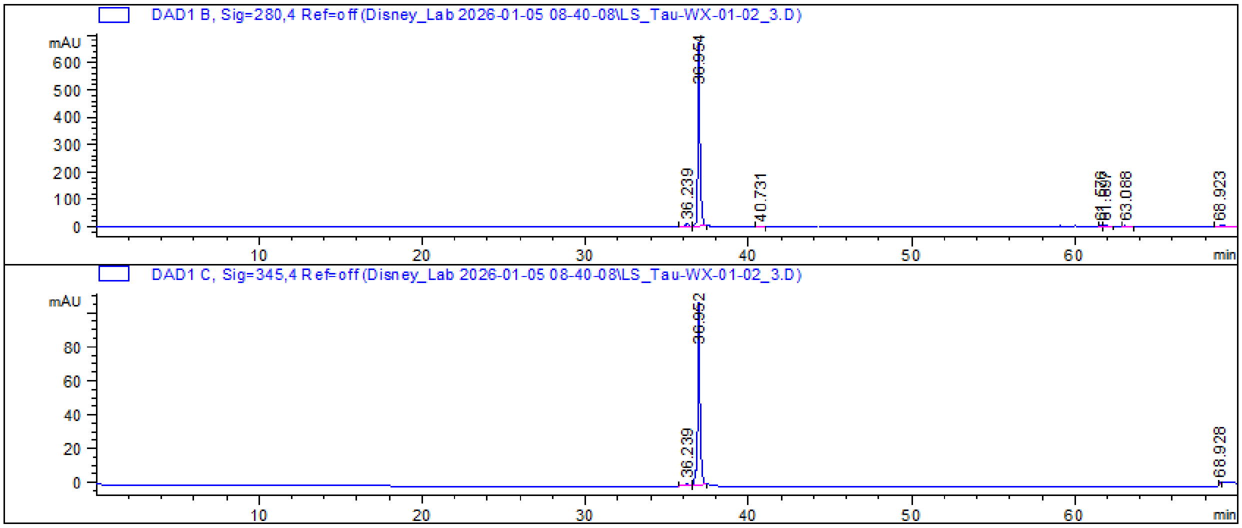

HRMS

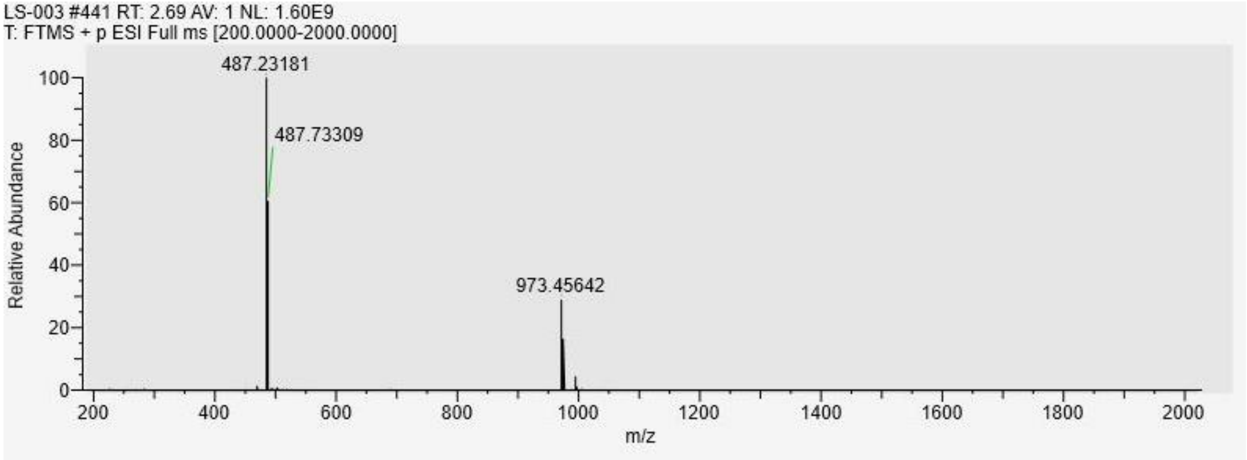
LS-003 #441 RT: 2.69 AV: 1 NL: 1.60E9
T: FTMS + p ESI Full ms [200.0000-2000.0000]
487.23181
487.73309
973.45642
Relative Abundance
m/z

#### Tau-WX-01-10

##### $^{1}H$ NMR

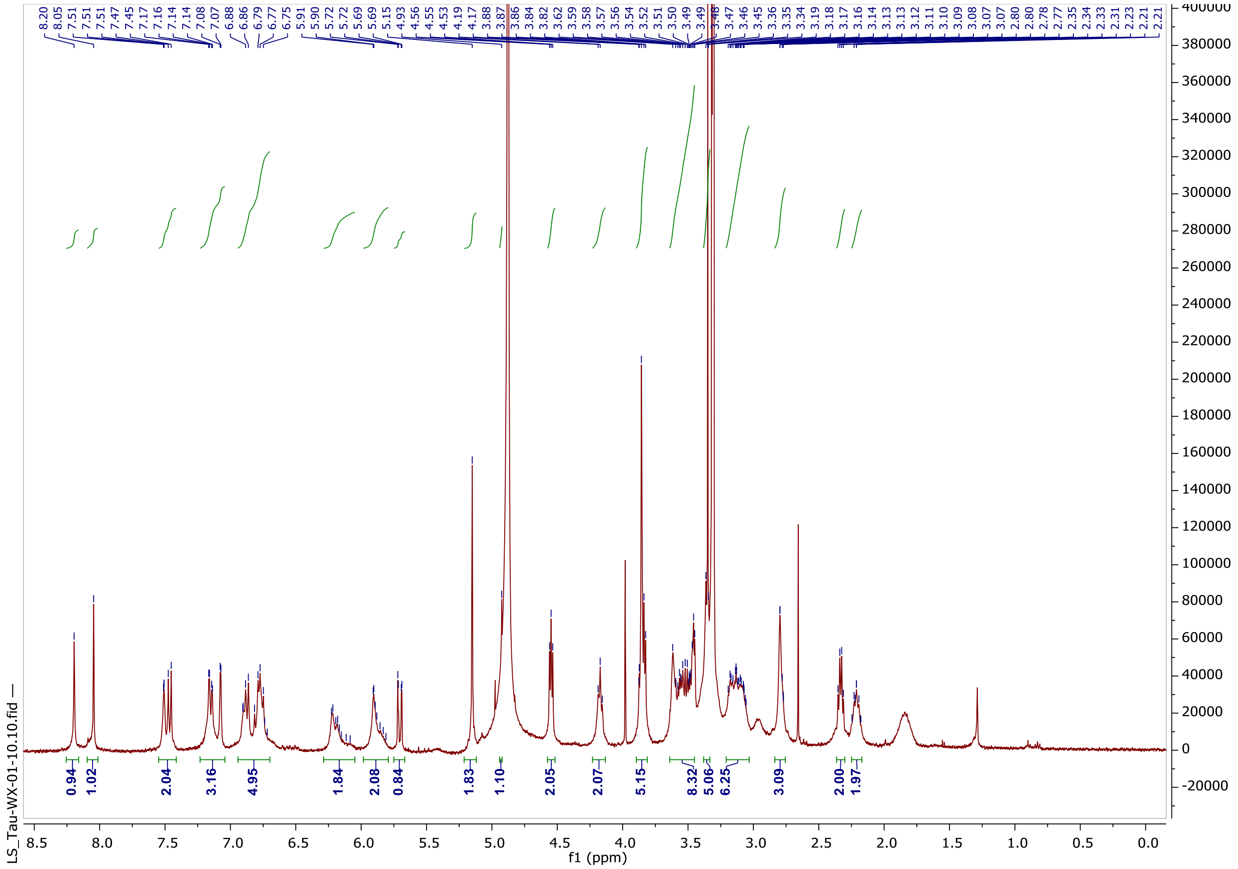

##### Analytical HPLC

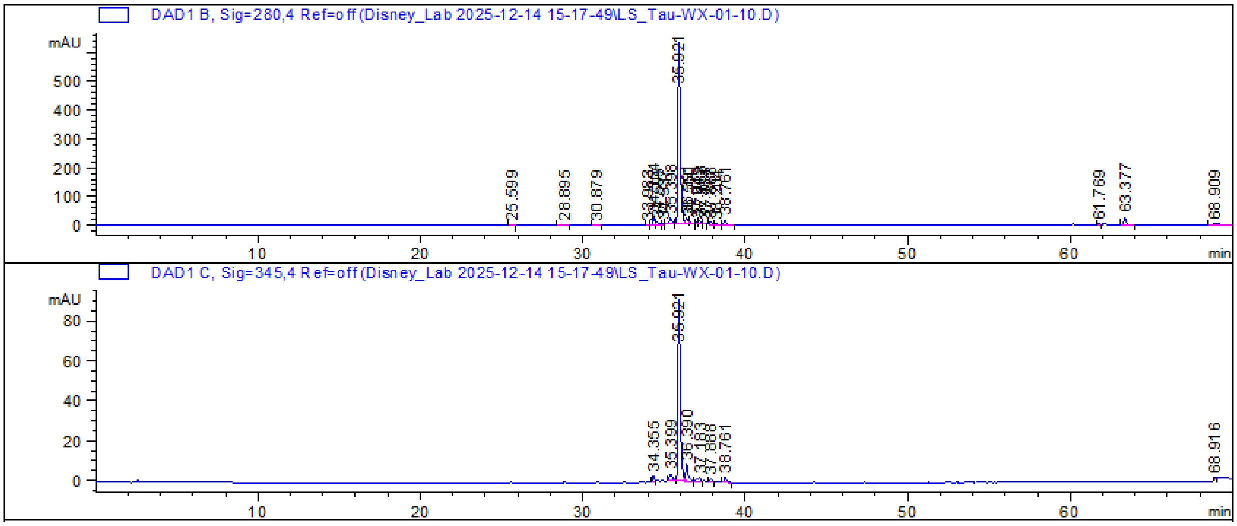

HRMS

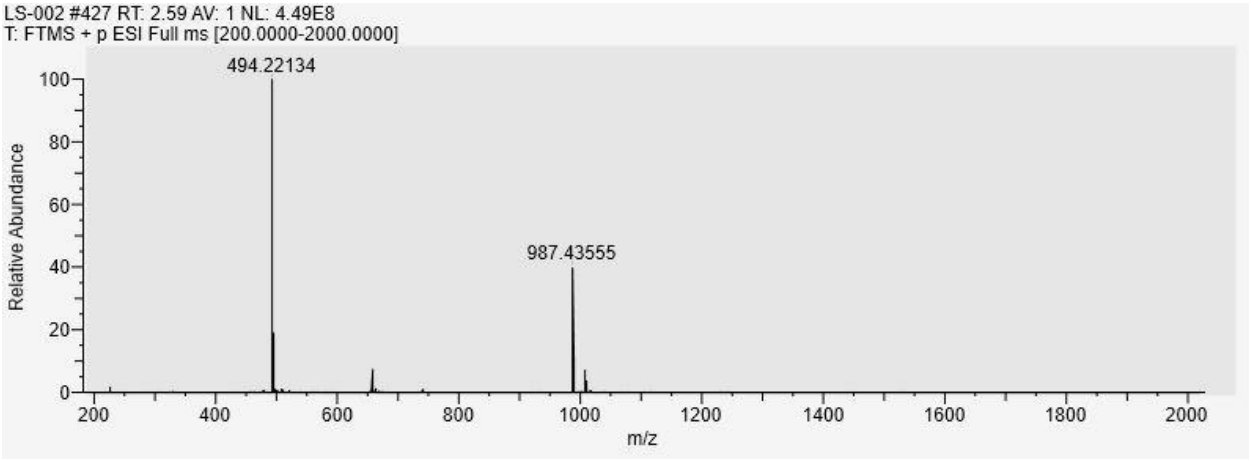
LS-002 #427 RT: 2.59 AV: 1 NL: 4.49E8
T: FTMS + p ESI Full ms [200.0000-2000.0000]
494.22134
987.43555
Relative Abundance
m/z
0
20
40
60
80
100
200
400
600
800
1000
1200
1400
1600
1800
2000

#### Tau-C3-WX-01-02

Analytical HPLC

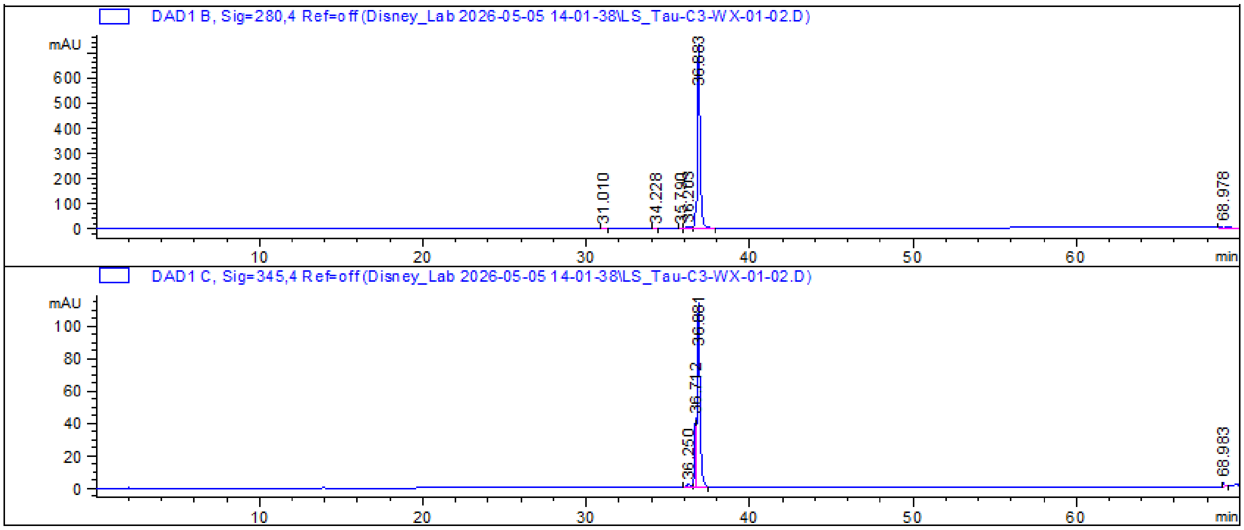


HRMS

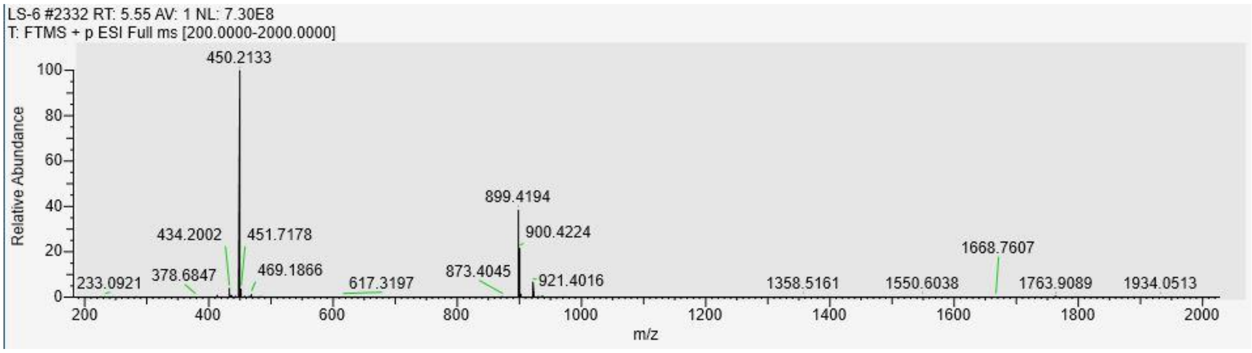

#### Tau-C6-WX-01-02

$^1$H NMR

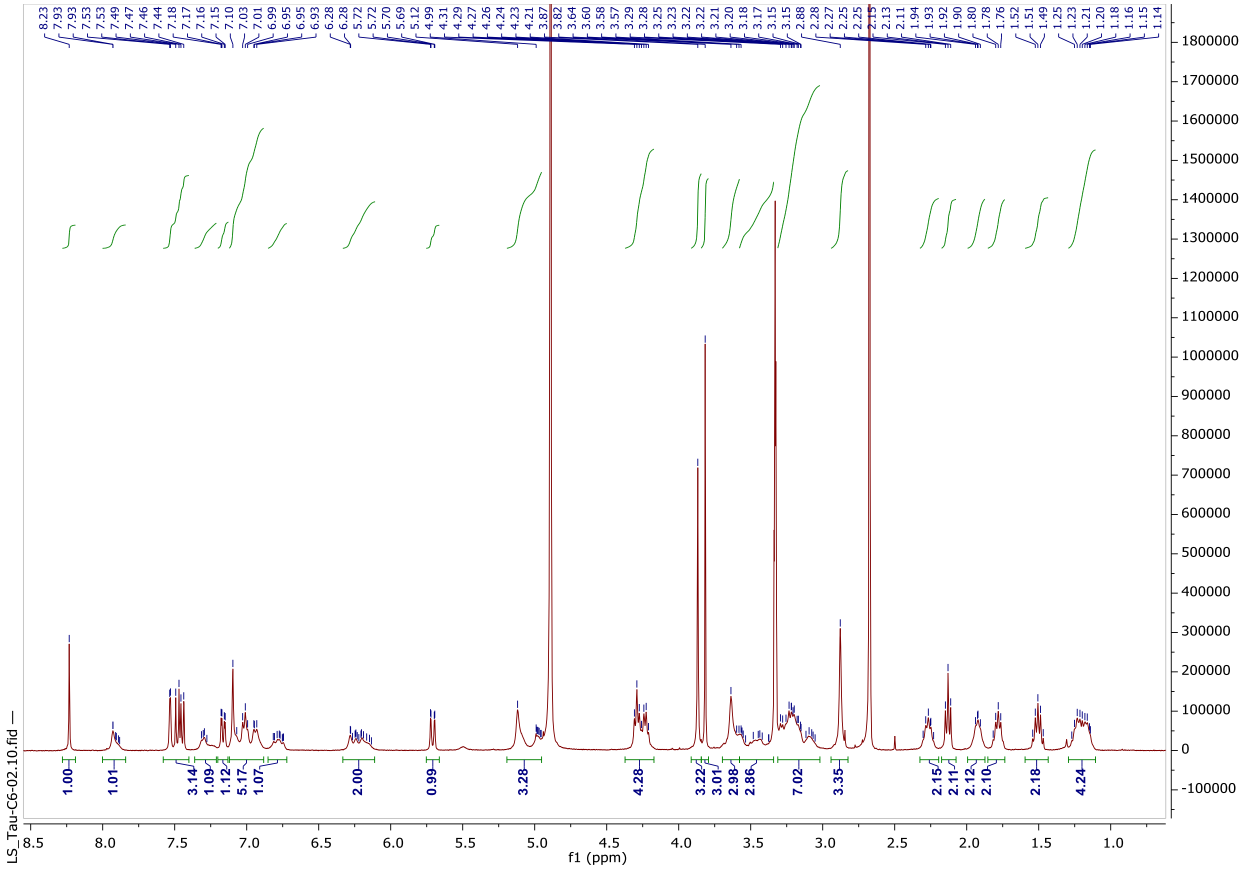

Analytical HPLC

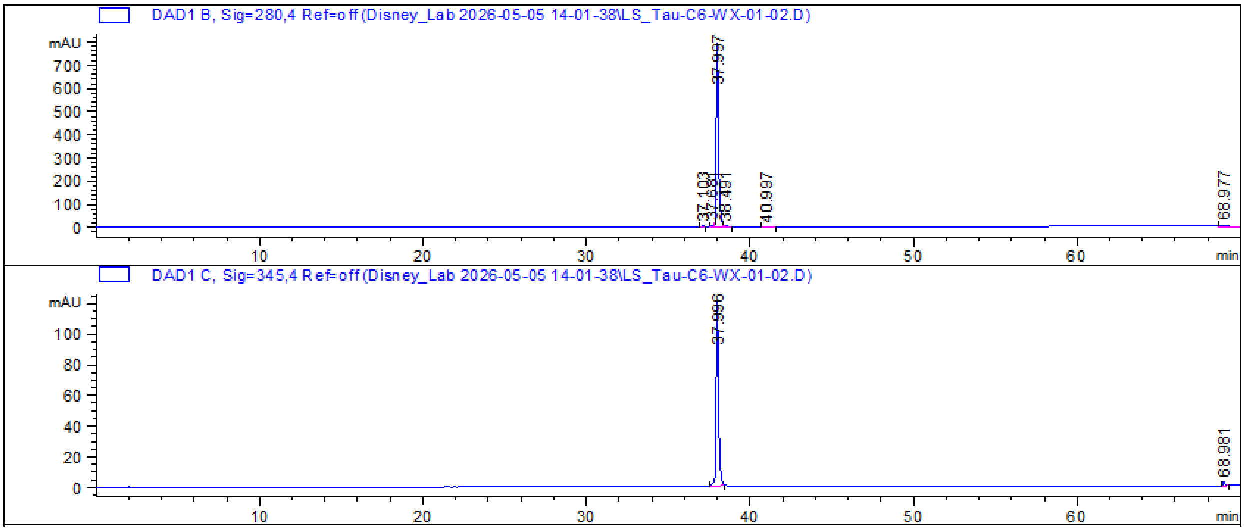

HRMS

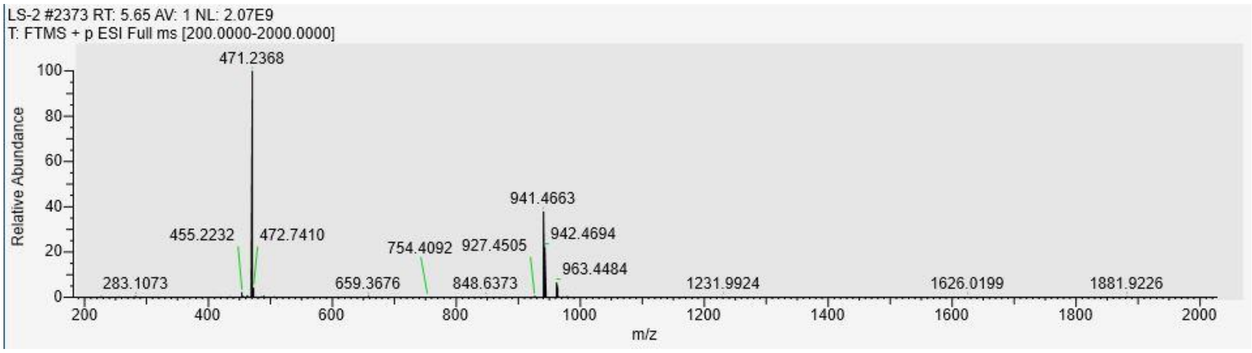
LS-2 #2373 RT: 5.65 AV: 1 NL: 2.07E9
T: FTMS + p ESI Full ms [200.0000-2000.0000]
Relative Abundance
471.2368
455.2232
472.7410
754.4092
927.4505
941.4663
942.4694
963.4484
283.1073
659.3676
848.6373
1231.9924
1626.0199
1881.9226
m/z

#### Tau-PEG3-WX-01-02

Analytical HPLC

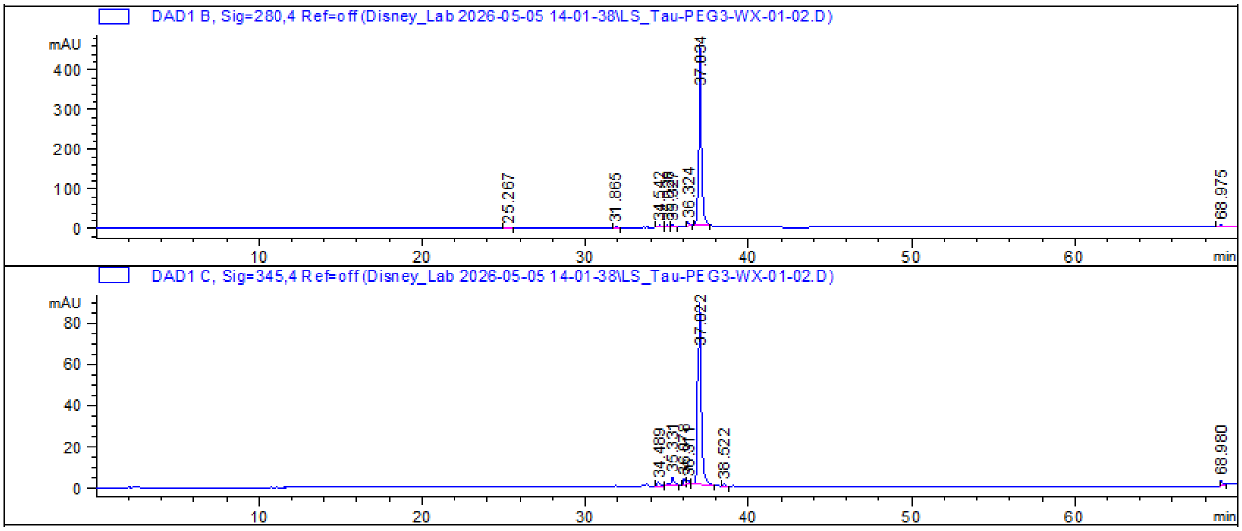


HRMS

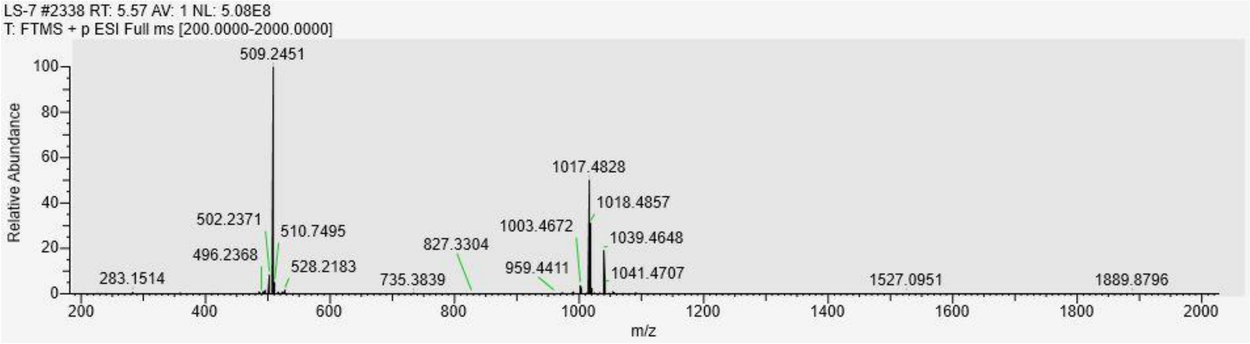

#### Tau-PEG4-WX-01-02

##### $^1$H NMR

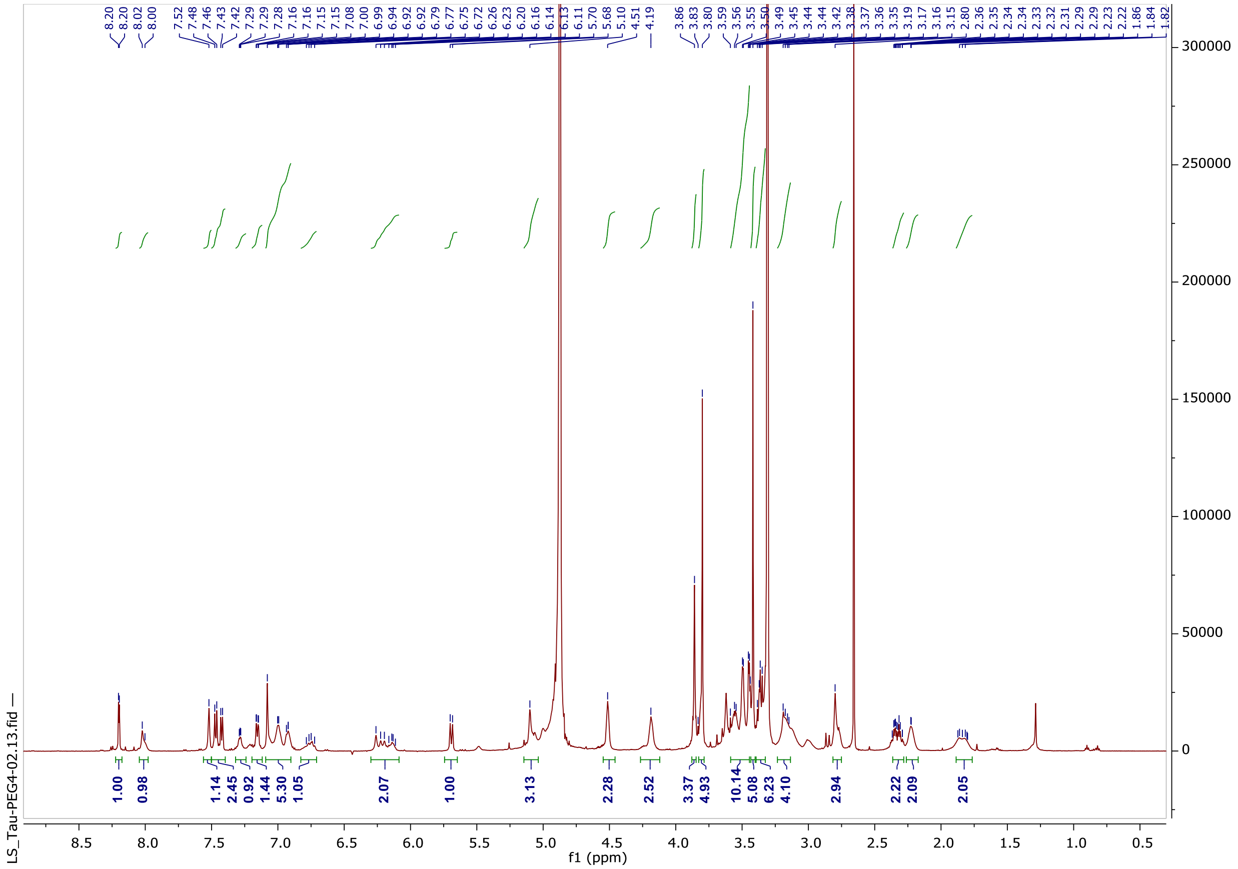

##### Analytical HPLC

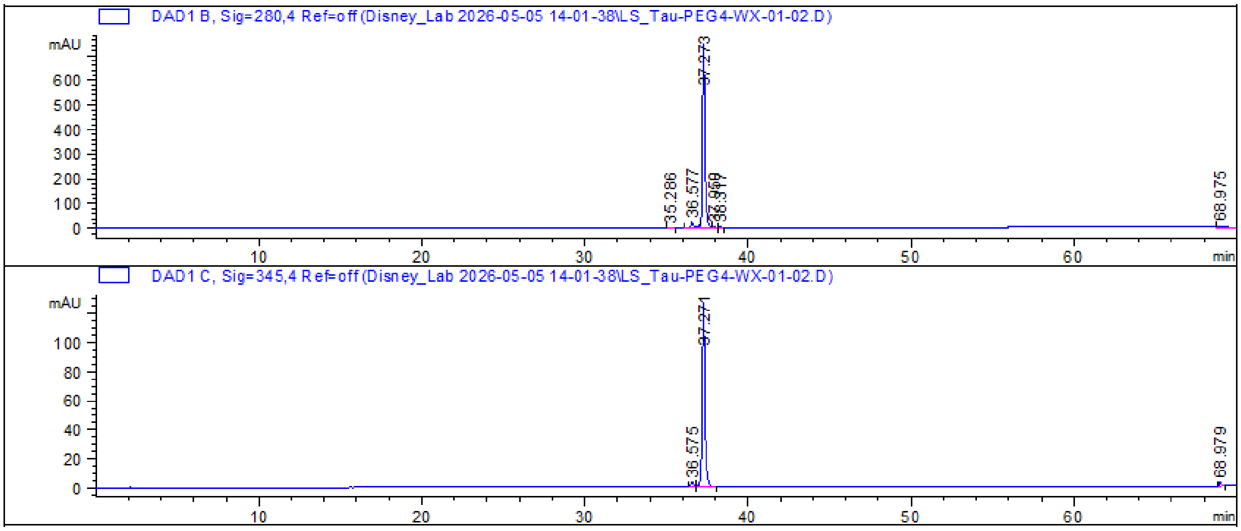


##### HRMS

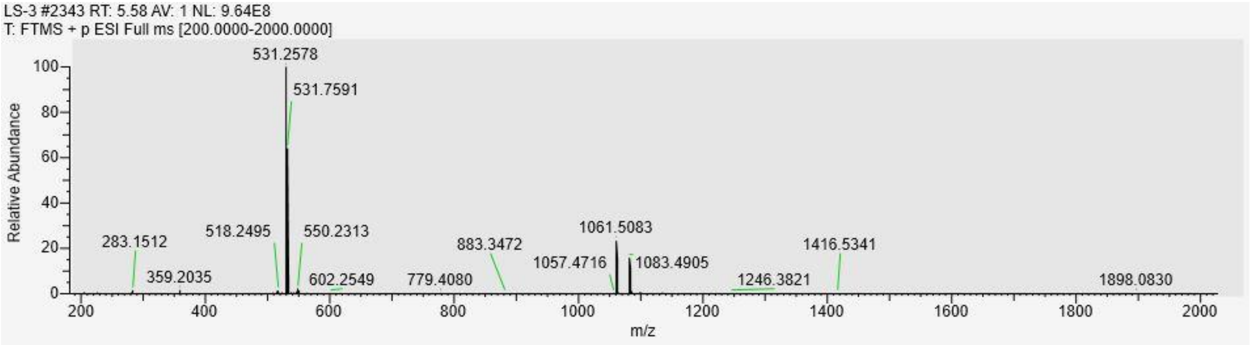

LS-3 #2343 RT: 5.58 AV: 1 NL: 9.64E8
T: FTMS + p ESI Full ms [200.0000-2000.0000]
531.2578
531.7591
283.1512
359.2035
518.2495
550.2313
602.2549
779.4080
883.3472
1057.4716
1061.5083
1083.4905
1246.3821
1416.5341
1898.0830
Relative Abundance
m/z

#### Tau-PEG6-WX-01-02

##### Analytical HPLC

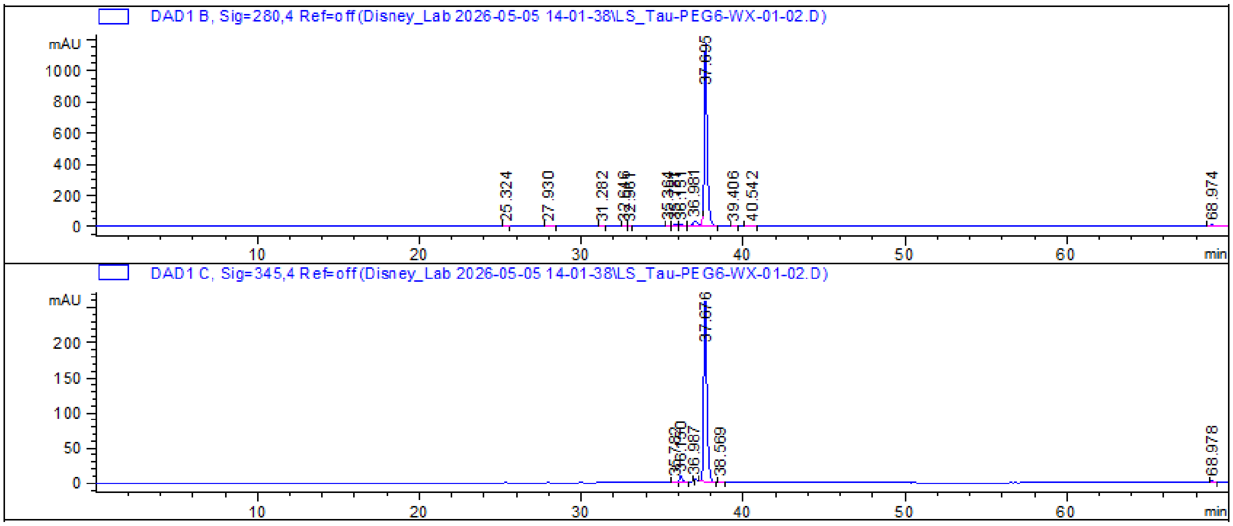


##### HRMS

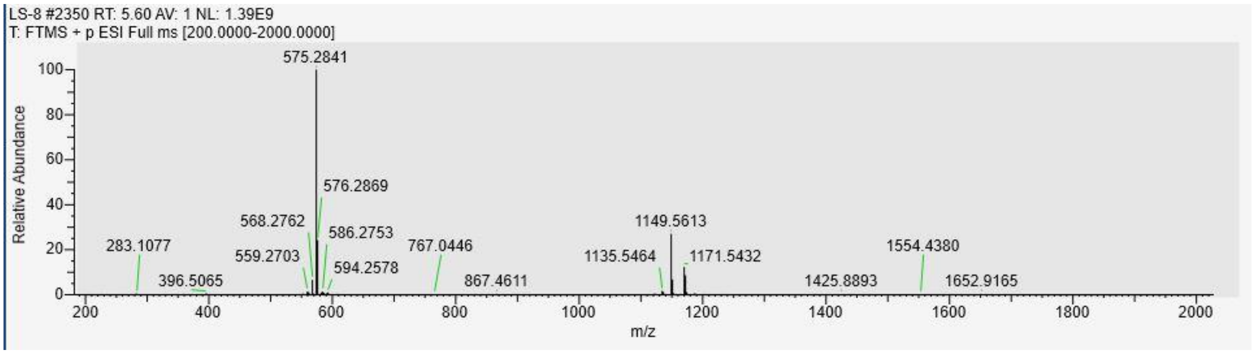

**Tau-PEG8-WX-01-02**

Analytical HPLC

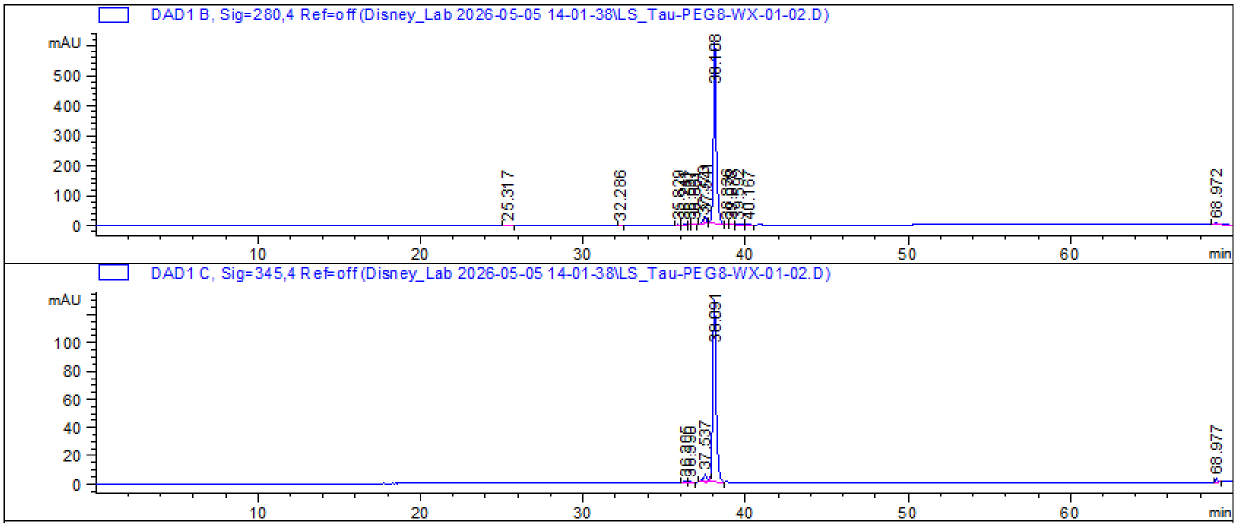


HRMS

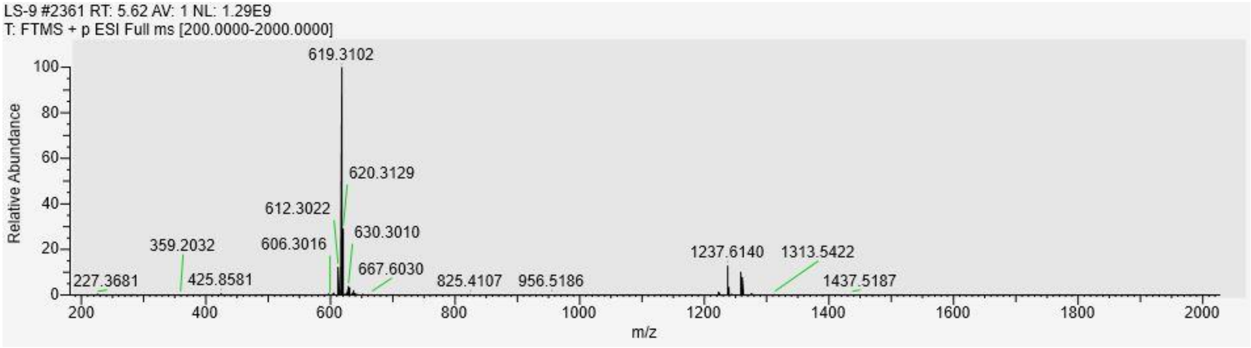

#### Tau-Pip1-WX-01-02

$^{1}$H NMR

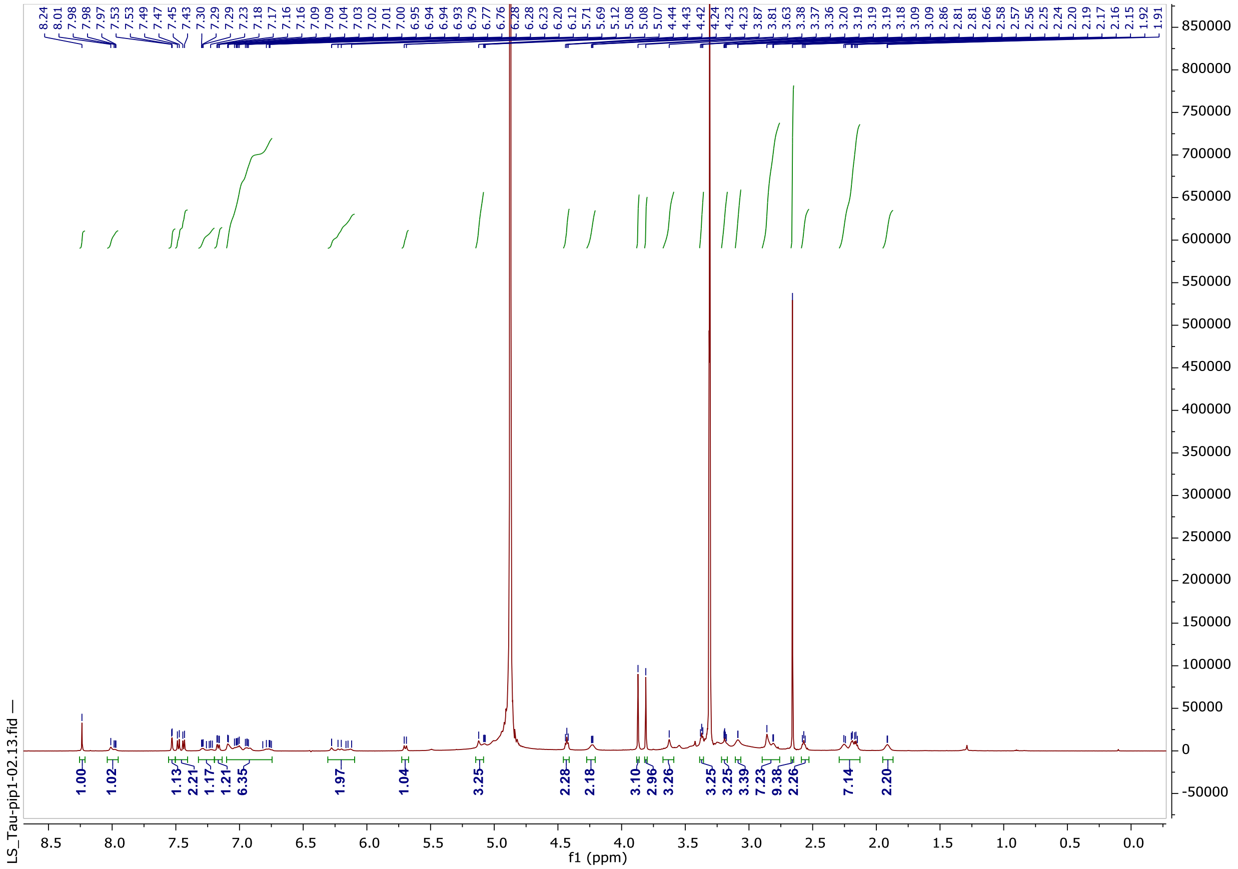

Analytical HPLC

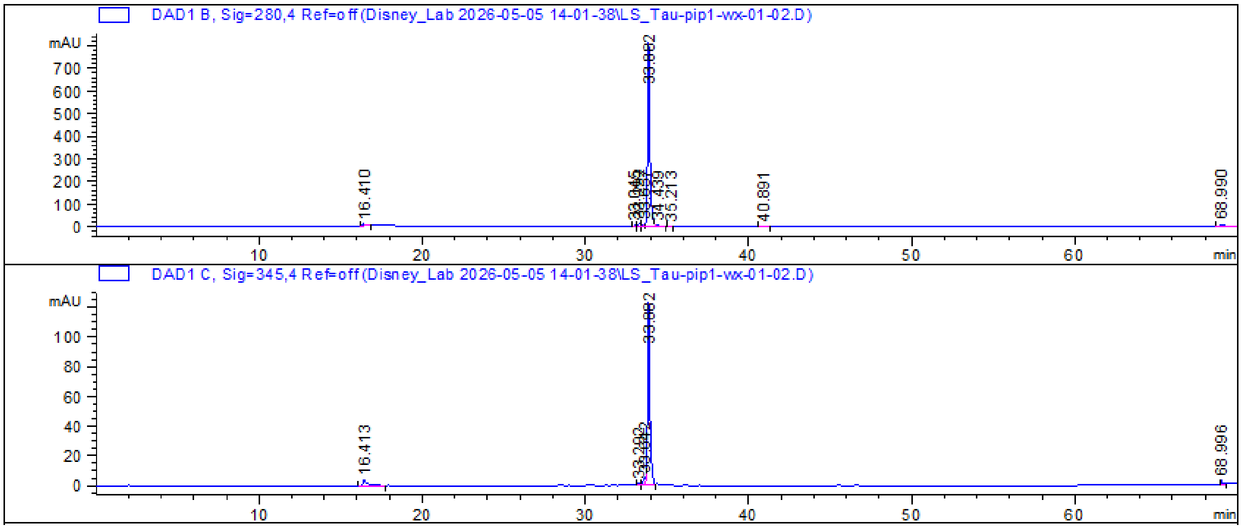

HRMS

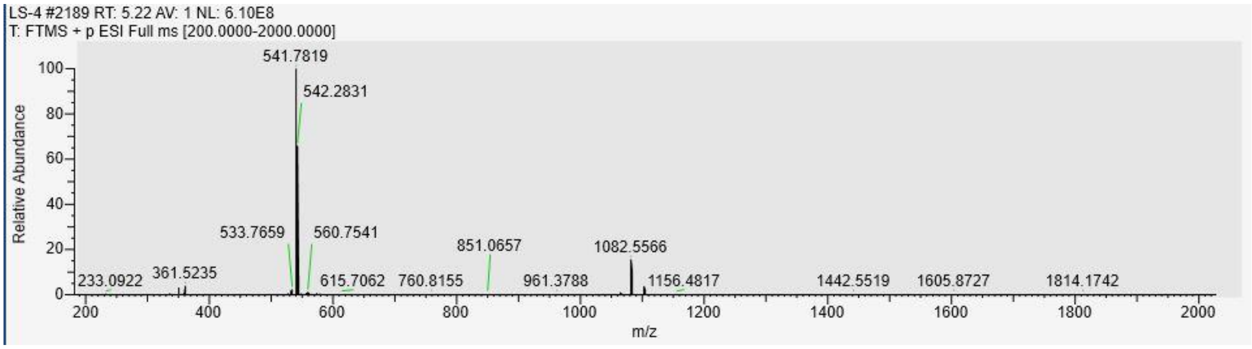
LS-4 #2189 RT: 5.22 AV: 1 NL: 6.10E8
T: FTMS + p ESI Full ms [200.0000-2000.0000]
541.7819
542.2831
533.7659
560.7541
851.0657
1082.5566
233.0922
361.5235
615.7062
760.8155
961.3788
1156.4817
1442.5519
1605.8727
1814.1742
Relative Abundance
m/z

#### Tau-Pip2-WX-01-02

$^1$H NMR

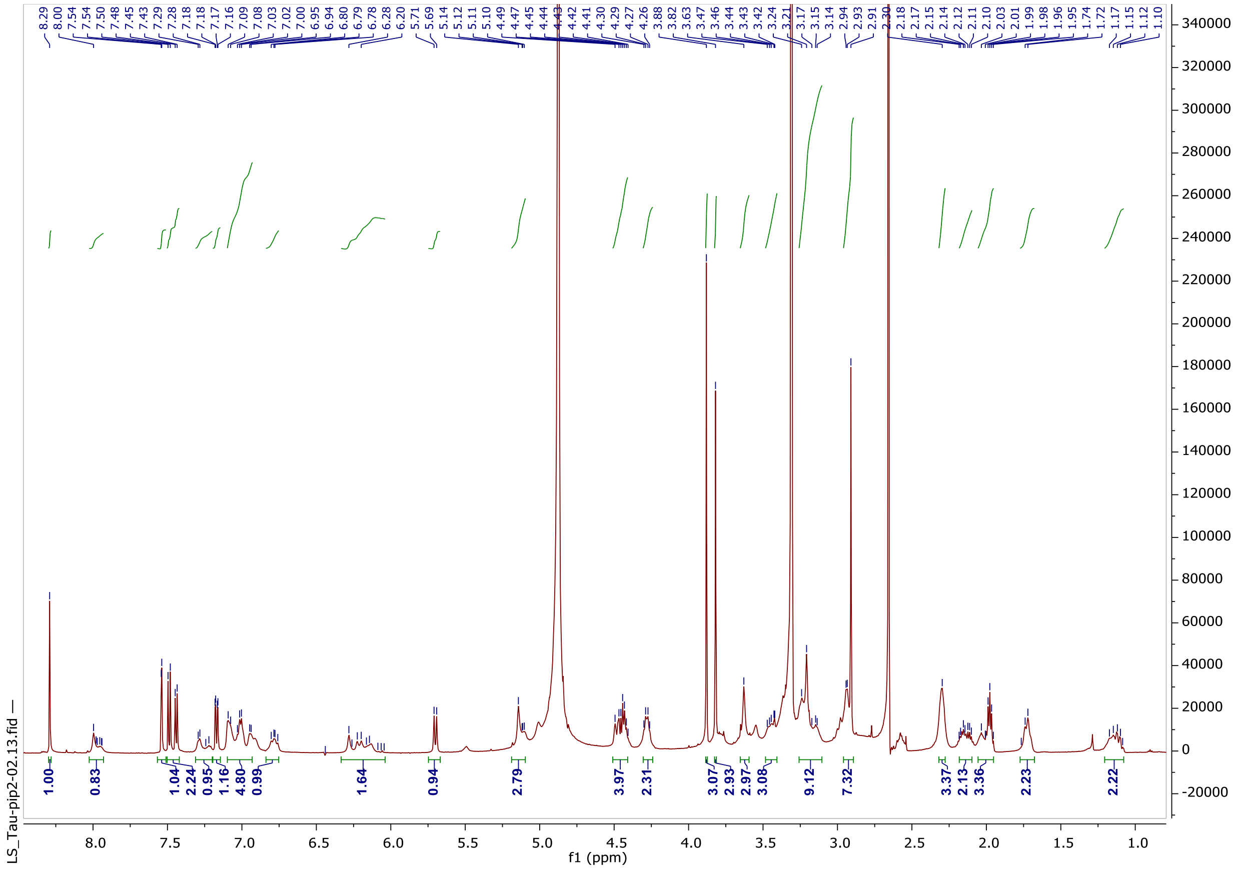

Analytical HPLC

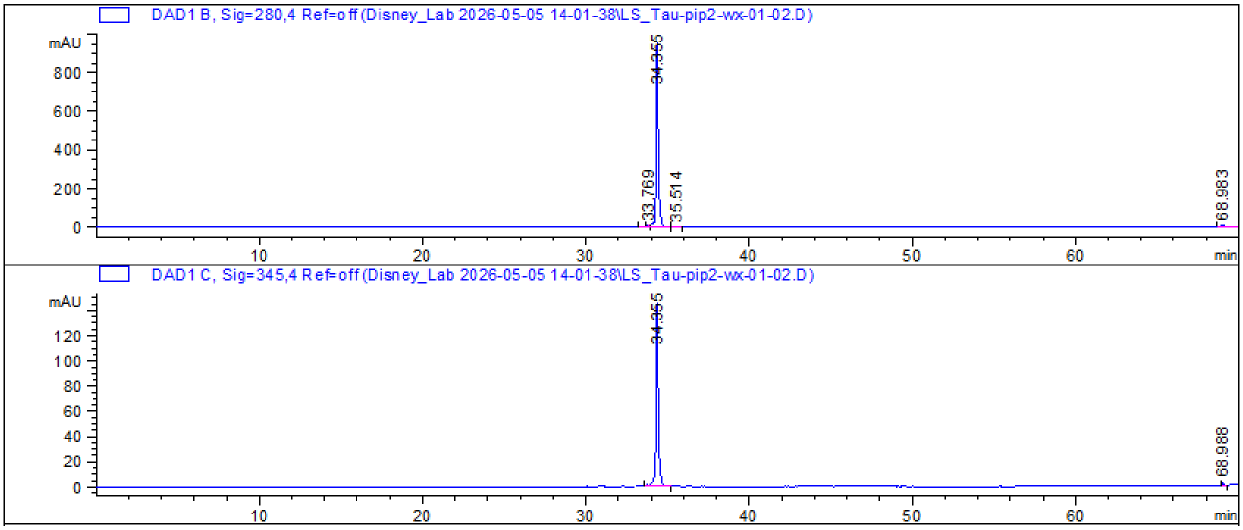

HRMS

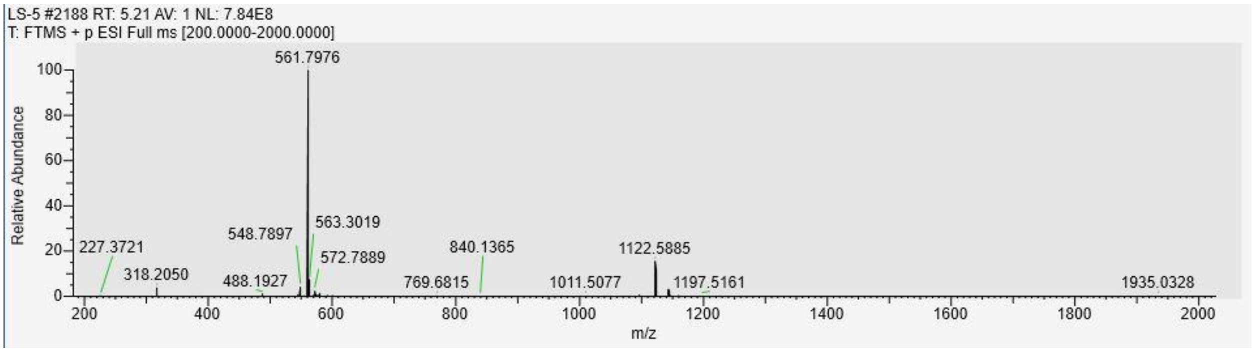
LS-5 #2188 RT: 5.21 AV: 1 NL: 7.84E8
T: FTMS + p ESI Full ms [200.0000-2000.0000]
Relative Abundance
m/z
561.7976
548.7897
563.3019
572.7889
227.3721
318.2050
488.1927
769.6815
840.1365
1011.5077
1122.5885
1197.5161
1935.0328

#### *o*-AP1867 Tracer

##### Analytical HPLC

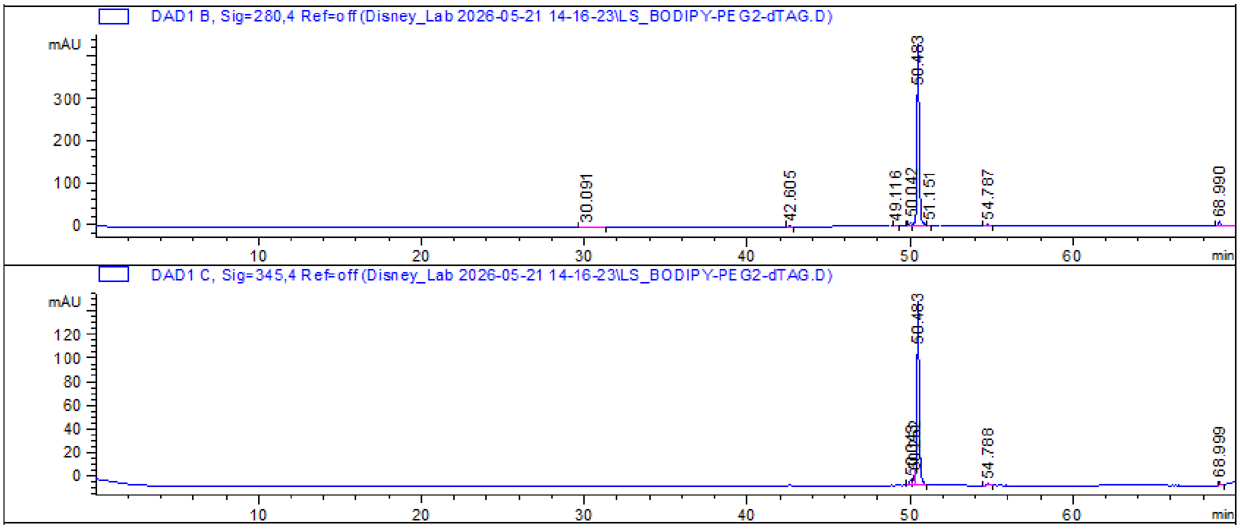


##### HRMS

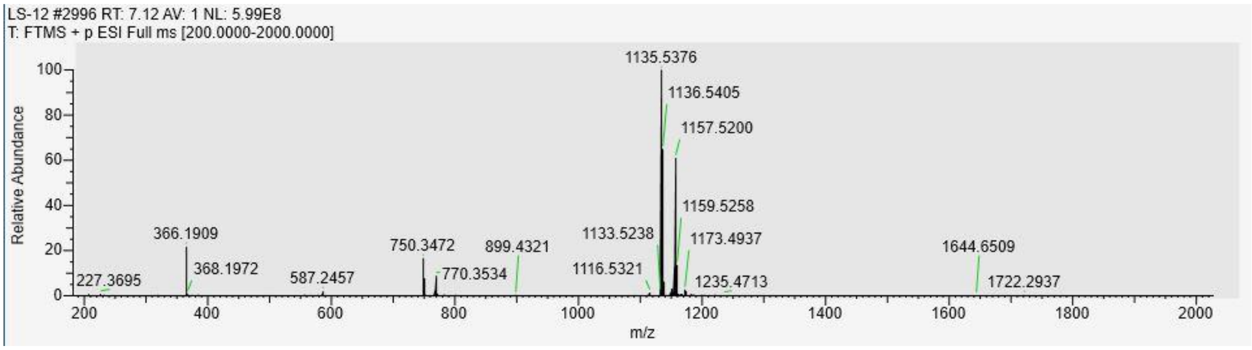
